# Identification and analysis of splicing quantitative trait loci across multiple tissues in the human genome

**DOI:** 10.1101/2020.05.29.123703

**Authors:** Diego Garrido-Martín, Beatrice Borsari, Miquel Calvo, Ferran Reverter, Roderic Guigó

## Abstract

We have developed an efficient and reproducible pipeline for the discovery of genetic variants affecting splicing (sQTLs), based on an approach that captures the intrinsically multivariate nature of this phenomenon. We employed it to analyze the multi-tissue transcriptome GTEx dataset, generating a comprehensive catalogue of sQTLs in the human genome. A core set of these sQTLs is shared across multiple tissues. Downstream analyses of this catalogue contribute to the understanding of the mechanisms underlying splicing regulation. We found that sQTLs often target the global splicing pattern of genes, rather than individual splicing events. Many of them also affect gene expression, but not always of the same gene, potentially uncovering regulatory loci that act on different genes through different mechanisms. sQTLs tend to be preferentially located in introns that are post-transcriptionally spliced, which would act as hotspots for splicing regulation. While many variants affect splicing patterns by directly altering the sequence of splice sites, many more modify the binding of RNA-binding proteins (RBPs) to target sequences within the transcripts. Genetic variants affecting splicing can have a phenotypic impact comparable or even stronger than variants affecting expression, with those that alter RBP binding playing a prominent role in disease.

## Introduction

Alternative splicing (AS) is the process through which multiple transcript isoforms are produced from a single gene^1^. It is a key mechanism that increases functional complexity in higher eukaryotes^2^. Often, its alteration leads to pathological conditions^3^. AS is subject to a tight regulation, usually tissue-, cell type- or condition-specific, that involves a wide range of *cis* and *trans* regulatory elements^4,5^. Since AS is generally coupled with transcription, transcription factors and chromatin structure also play a role in its regulation^6^.

In recent years, transcriptome profiling of large cohorts of genotyped individuals by RNA-seq has allowed the identification of genetic variants affecting AS, i.e. splicing quantitative trait loci or sQTLs^7–12^. sQTL analyses in a variety of experimental settings have helped to gain insight into the mechanisms underlying GWAS associations for a number of traits, such as adipose-related traits^13^, Alzheimer’s disease^10^, schizophrenia9 or breast cancer^14^, among others. sQTLs might actually contribute to complex traits and diseases at a similar or even larger degree than variants affecting gene expression^15^.

The vast majority of methods commonly used for sQTL mapping treat splicing as a univariate phenotype. They assess association between genetic variants and the abundance of individual transcripts^7,16^, or the splicing of individual exons^9,17^ or introns^12,15^. However, this approach ignores the strongly correlated structure of AS measurements (e.g. at constant gene expression level, higher levels of a splicing isoform correspond necessarily to lower levels of other isoforms). In contrast, we propose an approach that takes into account the intrinsically multivariate nature of alternative splicing: variants are tested for association with a vector of AS phenotypes, such as the relative abundances of the transcript isoforms of a gene or the intron excision ratios of an intron cluster obtained by LeafCutter^18^.

Based on this approach, we have developed a pipeline for efficient and reproducible sQTL mapping. We have employed it to leverage the multi-tissue transcriptome data generated by the Genotype-Tissue Expression (GTEx) Consortium, producing a comprehensive catalogue of genetic variants affecting splicing in the human genome. Downstream analyses of this catalogue uncover a number of relevant features regarding splicing regulation. Thus, consistent with the multivariate nature of splicing, we have observed that sQTLs tend to involve multiple splicing events. A substantial fraction of sQTLs also affects gene expression, a reflection of the intimate relationship between splicing and transcription. We have found, however, many cases in which the sQTL affects the expression of a gene other than the sQTL target. In these cases, the pleiotropic effect of the regulatory locus is not mediated by the interplay between the splicing and transcription processes, but it is exerted through different mechanisms, acting upon different genes that otherwise may not appear to be directly interacting. We have also found that sQTLs tend to be preferentially located in introns that are post-transcriptionally spliced: these introns would be consequently acting as hotspots for splicing regulation. While many variants affect splicing patterns by directly altering the sequence of splice sites, many more modify the binding of RNA-binding proteins (RBPs) to target sequences within the transcripts. We have observed that sQTLs often impact GWAS traits and diseases more than variants affecting only gene expression, confirming earlier reports which suggest that splicing mutations underlie many hereditary diseases^15,19^. For many conditions, GWAS associations are particularly strong for sQTLs altering RBP binding sites.

## Results

### Identification of *cis* splicing QTLs across GTEx tissues

For sQTL mapping, we developed `sQTLseekeR2`, a software based on `sQTLseekeR`^20^, which identifies genetic variants associated with changes in the relative abundances of the transcript isoforms of a given gene. `sQTLseekeR` uses the Hellinger distance to estimate the variability of isoform abundances across observations, and Anderson’s method^21,22^, a non-parametric analogue to multivariate analysis of variance, to assess the significance of the associations (see Methods and Supplementary Note 1). Among other enhancements, `sQTLseekeR2` improves the accuracy and speed of the *p* value calculation, and allows to account for additional covariates before testing for association with the genotype, while maintaining the multivariate statistical test in `sQTLseekeR`. It also implements a multiple testing correction scheme that empirically characterizes, for each gene, the distribution of *p* values expected under the null hypothesis of no association (see Methods and Supplementary Note 1). To ensure highly parallel, portable and reproducible sQTL mapping, we embedded `sQTLseekeR2` in a Nextflow^23^ (plus Docker, https://www.docker.com/) computational workflow named `sqtlseeker2-nf`, available at https://github.com/dgarrimar/sqtlseeker2-nf.

Here we extensively analyze the sQTLs identified by `sqtlseeker2-nf`, using the expression and genotype data produced by the GTEx Consortium. For most of the analyses, we employed isoform quantifications obtained from the V7 release (dbGaP accession *phs000424.v7.p2*), corresponding to 10,361 samples from 53 tissues of 620 deceased donors. 48 tissues with sample size ≥ 70 were selected for sQTL analyses. We tested variants in a *cis* window defined as the gene body plus 5 Kb upstream and downstream the gene boundaries. In addition, to demonstrate that the statistical framework of `sQTLseekeR2` is not restricted to the analysis of transcript abundances, but it can leverage other splicing-related multivariate phenotypes, we have also computed the sQTLs based on the intron excision ratios obtained by LeafCutter^18^ from the GTEx RNA-seq data (Supplementary Note 2). Finally, we also provide the sQTLs identified by `sqtlseeker2-nf` in GTEx V8 (Supplementary Note 3), which can be compared to the sQTLs produced by the GTEx Consortium in an upcoming publication^12^.

At a 0.05 false discovery rate (FDR), we found in GTEx V7 a total of 210,485 *cis* sQTLs affecting 6,963 genes (6,685 protein coding genes and 278 long intergenic non-coding RNAs, lincRNAs). On average, per tissue, we identified 1,158 sGenes (Table S1). 44% and 34% of all tested protein coding genes and lincRNAs, respectively, were found to be sGenes. In an analogous experimental setting, the GTEx Consortium reported genetic variants affecting expression (expression QTLs, eQTLs) for 95% and 71% of all tested protein coding genes and lincRNAs, respectively^24^. To illustrate the nature of the sQTLs identified with `sqtlseeker2-nf`, in Fig. 1 we show the example of the SNP rs2295682, an sQTL for the gene RBM23 shared across 46 tissues, with larger effect in brain subregions such as cortex. The SNP strongly affects the relative abundances of the AS isoforms of the target gene, the dominant isoform depending on the genotype at the sQTL.

**Figure 1.**
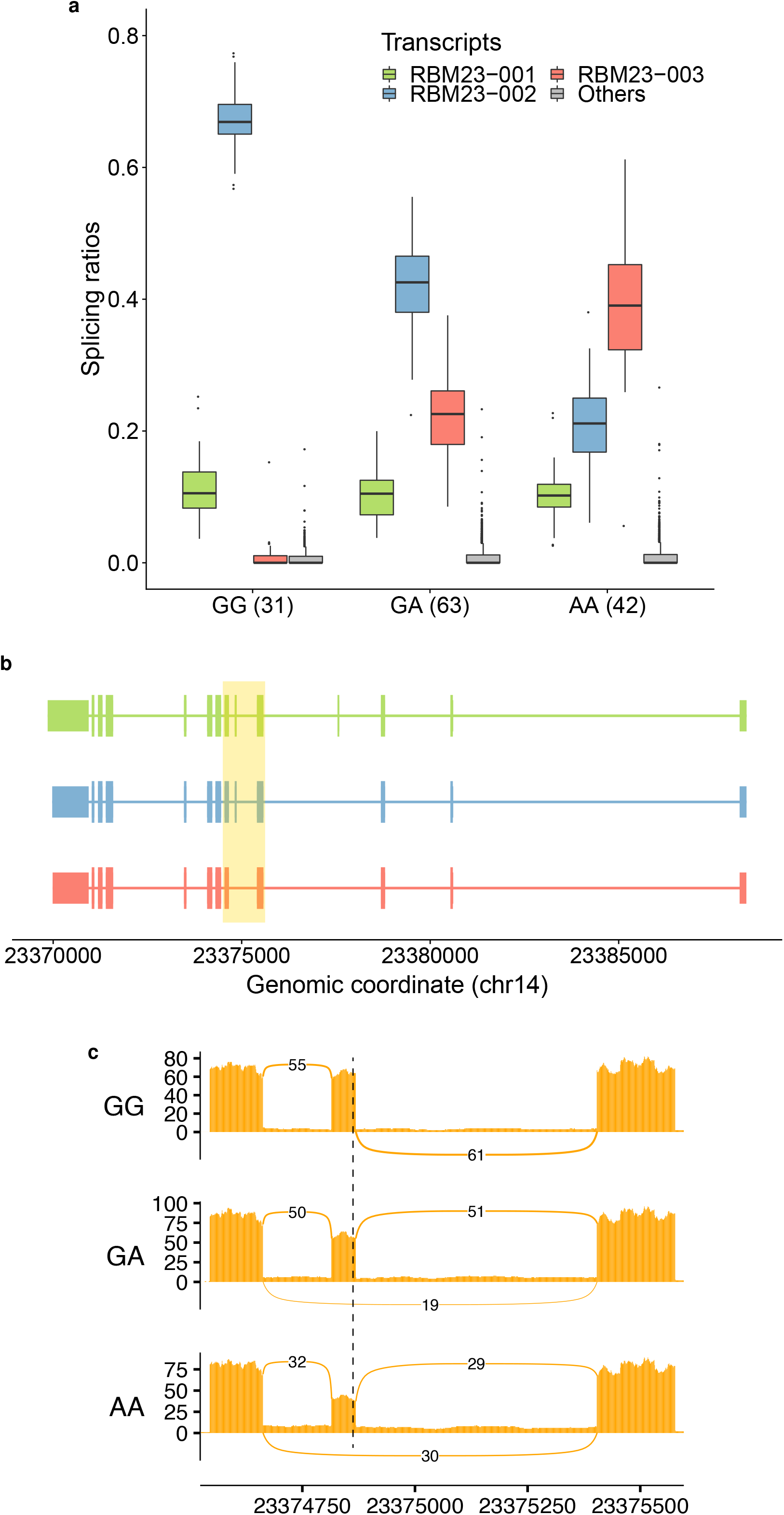
sQTL example. **a)** Relative abundances of the three most expressed isoforms in the brain cortex from the gene RBM23 (chr14:23,369,854-23,388,393, reverse strand, RBM23-001, RBM23-002 and RBM23-003, all protein coding), for each genotype group at the rs2295682 locus (chr14:23,374,862, G/A in the reverse strand). RBM23 encodes for an RNA-binding protein that may be itself involved in splicing. The least abundant isoforms are grouped in *Others*. The number of individuals in each genotype group is shown between parentheses. Individuals that are homozygous for the reference allele (GG) at the SNP locus, express preferentially RBM23-002 (blue), while they barely express RBM23-003 (red). In contrast, AA homozygous express preferentially RBM23-003 (red). Heterozygous individuals exhibit intermediate abundances. RBM23-001 (green) has similar levels in the three genotype groups. **b)** Exonic structure of the isoforms and location of exons 5, 6 and 7 (highlighted area). Compared to RBM23-001 (green), RBM23-002 (blue) lacks exon 6, and RBM23-003 (red), exons 4 and 6. **c)** Sashimi plot (corresponding to the highlighted area in b) displaying the mean exon inclusion of exon 6 of RBM23 across all brain cortex samples of each genotype group at rs2295682, obtained by `ggsashimi`^84^. The dashed line marks the location of the SNP. The number of reads supporting exon skipping increases with the number of copies of the alternative allele A, matching the changes observed in isoform abundances. This allele has been previously associated with increased skipping of exon 6^85^.

As expected, the number of sGenes over the number of tested genes grows with the tissue sample size (*r*^2^ = 0.91). This is explained by the gain of power to detect sQTLs as the number of samples increases (Fig. 2a). No signs of saturation are observed. Some tissues, such as skeletal muscle or whole blood (with less sQTLs than expected) and testis (with more sQTLs than expected) escape the general trend. This was also observed for eQTLs24. The cell type heterogeneity of the tissue, estimated using xCell25, does not seem to have a large impact on sQTL discovery compared to the tissue sample size (the partial correlation between the number of sGenes over the number of tested genes and the estimated cell type heterogeneity, controlling for the tissue sample size, is 0.23, *p* value 0.11, see Methods).

**Figure 2.**
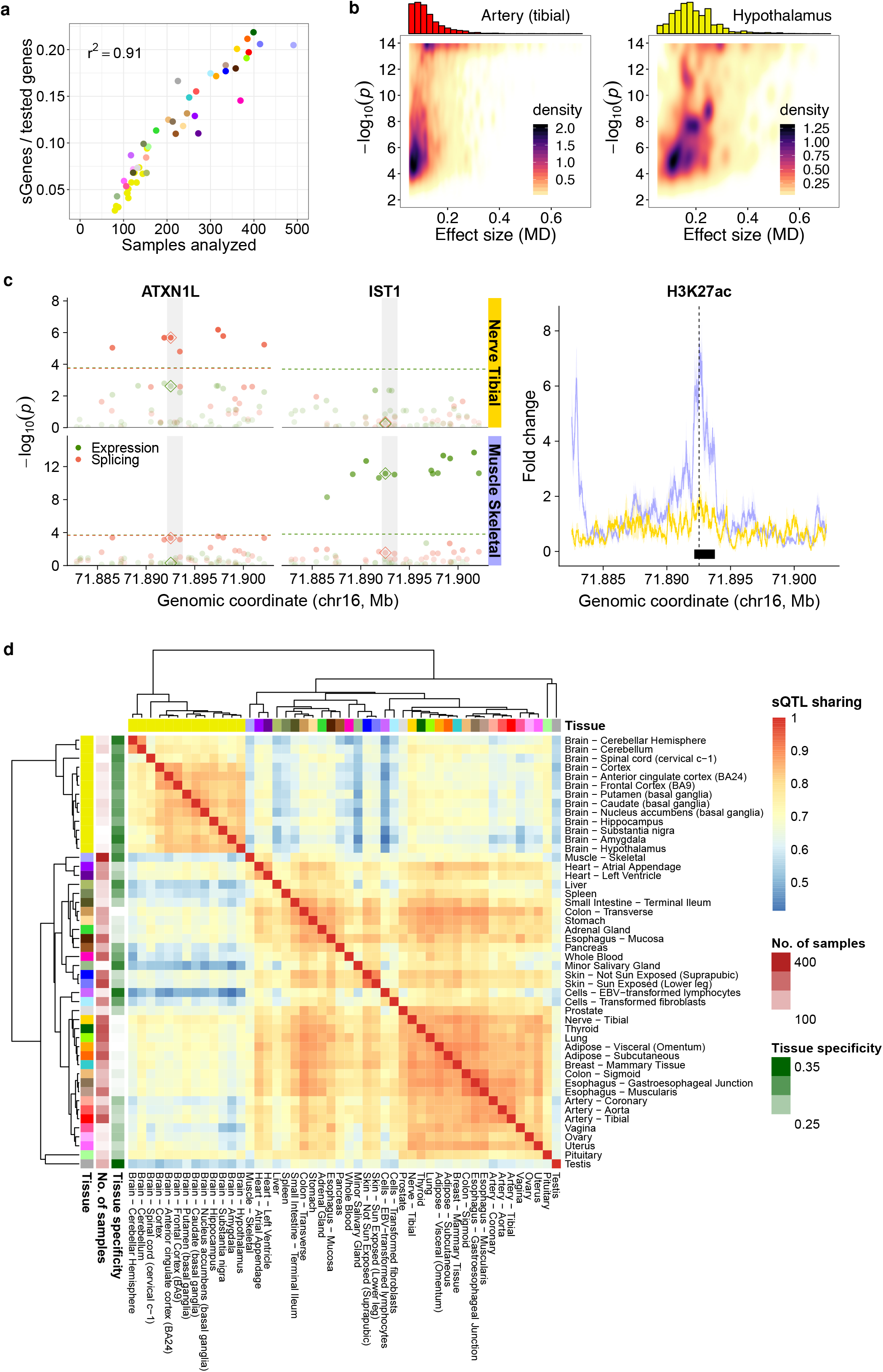
Overall results, heteropleiotropy and sQTL sharing across tissues. **a)** Proportion of sGenes (over tested genes) per tissue (y-axis) with respect to the tissue sample size (x-axis). Tissue color codes are shown in Table S1. **b)** For two tissues with markedly different sample sizes, such as tibial artery (upper panel, 388 samples) and hypothalamus (lower panel, 108 samples), we display the effect sizes (MD values, x-axis) of significant sQTLs vs the −*log*_10_ of their association *p* value with the target sGene (y-axis). The density of points is shown, together with the sQTL effect size distribution. Note that MD for sQTLs is bounded to [0.05, 1] (see Methods). **c)** Example of an heteropleiotropic locus. The SNP rs8046859 (chr16:71,892,531, C/T), an sQTL for the gene ATXN1L (chr16:71,879,894-71,919,171, forward strand) in Nerve Tibial, but not in Muscle Skeletal. The SNP is not an eQTL for ATXN1 in any of the two tissues. In contrast, the SNP is an eQTL for the gene IST1 (chr16:71,879,899-71,962,913, forward strand) in Muscle Skeletal, but not in Nerve Tibial. The SNP is not an sQTL for IST1 in any of the two tissues. In the left panel, the dots represent the −*log*_10_ *p* values of association with the expression (green) and splicing (red) of the two genes in the two tissues, for variants in a 20Kb window centered at rs8046859 (the −*log*_10_ *p* values corresponding to rs8046859 are highlighted by a diamond). The transparency of the dots depends on the −*log*_10_ *p* value. The significance level for each molecular trait, gene and tissue is shown as a colored, dashed horizontal line. When this line is not present, the gene-level *p* value is above the 0.05 FDR threshold and hence no variant is significantly associated with this molecular trait in this tissue (see Methods). The shaded area represents the position of a H3K27ac ChIP-seq peak (see below). The right panel shows the fold-change signal of the H3K27ac histone mark with respect to the input across ENTEx donors in Nerve Tibial and Muscle Skeletal, in the same genomic region of the left panel. The solid line and coloured area correspond to the mean signal and its standard error across 4 ENTEx donors, respectively. The location of the SNP and the overlapping ChIP-seq peak (intersection of the peaks in the 4 donors) are also displayed. **d)**Heatmap of sQTL sharing across GTEx tissues. Sharing estimates (see Methods) range from 0 (low sharing, blue) to 1 (high sharing, red). In addition, hierarchical clustering of the tissues based on sQTL sharing is displayed, together with the tissue sample sizes and tissue specificity estimates.

sQTL effect sizes, measured as the absolute maximum difference (MD) in adjusted transcript relative expression between genotype groups (see Methods), are generally low to moderate (MD from 0.05 to 0.20). Nevertheless, around 20% of sQTLs account for large effects (MD ≥ 0.20). As one would expect, the median effect size detected across tissues drops substantially with increasing sample sizes (Fig. S1), given that larger sample sizes allow the detection of smaller effects. Fig. 2b represents sQTL effect sizes (MD values) vs *p* values, together with the distribution of the former, for tibial artery (n = 388) and hypothalamus (*n* = 108).

GO enrichment analysis of sGenes shows a wide variety of biological processes, including cellular transport, immune response, mitochondrial functions and, interestingly, RNA processing (Fig. S2a). This might suggest some mechanism of splicing autoregulation, as it has been previously described^26^. In contrast, tested genes without sQTLs are enriched in functions related to signaling and, especially, development (Fig. S2b). This resembles the behaviour reported for genes without eQTLs^24^, as it does the fact that genes without sQTLs are less expressed than sGenes in all tissues (Wilcoxon Rank-Sum test *p* value < 10^−16^).

The sQTLs found here are highly replicated in other studies. We compared them with those obtained in the Blueprint Project^27^ for three major human blood cell types (CD14^+^ monocytes, CD16^+^ neutrophils, and naive CD4^+^ T cells, see Methods). The majority of GTEx sQTLs replicate at 5% FDR (from *π*_1_ = 0.80 in brain subregions to *π*_1_ = 0.96 in whole blood). As expected, whole blood displays the highest sQTL replication rate (Fig. S3).

We characterized the types of alternative splicing (AS) events associated with sQTLs (see Methods, Fig. S4a). Note that here we also account for other relevant sources of transcript diversity, such as alternative transcription initiation and termination^28^. sQTLs generally involve multiple events (on average 2.63). Around 34% of sQTLs are related to at least one AS event involving internal exons and/or introns. Among them, exon skipping is the most frequent simple event (7% to 10% of all events). In addition, 58% of sQTLs are associated with events affecting first/last exons and untranslated regions (UTRs). The landscape of AS events associated with sQTLs is very similar across tissues. However, brain subregions present some particularities when compared to non-brain tissues, such as a larger proportion of exon skipping events and a smaller proportion of complex events involving the 3’ gene terminus (see Fig. S4b,c for details).

We found that 52% of the identified sQTLs are also eQTLs for the same gene and tissue, although the top sQTL coincides with the top eQTL only in 3% of the cases. This relatively large overlap, which departs from that reported in some previous studies^15^, matches what was observed for `sQTLseekeR` sQTLs in the GTEx pilot^29^. This is partially due to our sQTLs being able to involve transcriptional termini, in addition to canonical splicing events. It also indicates a substantial degree of co-regulation of gene expression and splicing, either at the level of transcription (e.g. variants that impact transcription and thus, splicing), or at the level of transcript stability (e.g. variants that affect splicing, and as a consequence, transcript stability and gene expression).

We focused on a set of 148,618 variants that were tested for association with both the expression and splicing of two genes (i.e. *g*_1_ and *g*_2_) or more, in at least two tissues, and identified 6,552 cases in which the variant is only sQTL for gene *g*_1_, but not for gene *g*_2_, in one tissue, and it is only eQTL for gene *g*_2_, but not for gene *g*_1_, in a different tissue (Fig. S5a). These cases uncover regulatory loci in the genome that, either through the same causal variant or through different causal variants in linkage disequilibrium (LD), have different effects on different genes through likely different molecular mechanisms. We term this phenomenon heteropleiotropy. We found evidence supporting the dual regulatory behaviour of heteropleiotropic loci. We identified the ChIP-seq peaks corresponding to six histone modifications from the ENTEx Project overlapping the heteropleiotropic variants above (see Methods). We hypothesized that loci with different regulatory effects (i.e. splicing and expression) in different tissues would be differently marked by histone modifications in these tissues. Indeed, we observed histone modification changes in 24% of the heteropleiotropic variants (Table S2), compared to 19% of non-heteropleiotropic variants (Fisher’s exact test test *p* value 0.045, see Methods). Regardless of the underlying causal structure, heteropleiotropic loci would uncover genomic regions that allow the coordinated regulation of different processes and affect different genes which otherwise do not appear to interact directly with each other. While further work is required to establish the relevance and generality of this phenomenon, figures 2c and S5b show some potentially interesting examples.

### sQTLs are highly shared across tissues

The large number of tissues available in GTEx allowed us to evaluate tissue sharing and specificity of sQTLs. For every pair of tissues, we selected variant-gene pairs tested in both and found significant in at least one, and computed the Pearson correlation (*r*) between their effect sizes (MD values). Hierarchical clustering based on these correlations grouped tissues with similar sQTL sharing patterns (Fig. 2d). A comparable clustering was obtained when using the more stringent Jaccard index (Fig. S6). Brain sub-regions cluster together and apart from the rest of the tissues, which form a second major cluster. We observe a high degree of sQTL sharing within each of the two groups (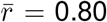 and 0.78, respectively), but lower between them 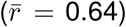. The same pattern was depicted for eQTLs in GTEx^24^. We further estimated tissue specificity as 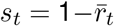, where 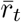 is the mean correlation between a given tissue t and the others (tissue specificity estimates shown in Fig. 2d). On average, brain sQTLs are more tissue-specific than non-brain sQTLs (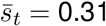 *vs* 0.25, Wilcoxon Rank-Sum test *p* value 9.32 · 10^−5^). Other tissues with relatively high tissue-specific sQTLs include testis (0.37), skeletal muscle (0.33) or liver (0.32).

sQTLs with large effects are more shared than those with smaller effects (Fig. S7a). As with eQTLs^24^, the detection of sQTLs with small effects requires larger sample sizes, thus sQTLs in tissues with small sample sizes tend to be more shared, while sQTLs identified in tissues with large sample sizes tend to be more tissue-specific (Fig. S7b). To rule out an effect of the sample size in the patterns of sQTL sharing, we downsampled the original dataset to 100, 200 and 300 samples per tissue, and evaluated again sQTL sharing. We found that the patterns of sQTL sharing above are replicated independently of the sample size (Fig. S8).

To capture more complex sharing patterns, we further designed a geometric approach that compares changes in the whole splicing phenotype due to sQTLs between tissues (see Methods and Fig. S9a). The derived tissue dendrogram (Fig. S9b) displayed high similarity with the ones generated by simpler approaches (i.e. based on MD values and *Jaccard* index), and also with the one obtained using *multivariate* adaptive shrinkage^30^ on `LeafCutter` sQTLs from GTEx V8^12^ (Fig. S9c). This strongly supports the robustness of the sQTL sharing patterns observed.

sGenes are also markedly shared: 66% of genes tested in all tissues are sGenes in at least two tissues. To identify tissue-specific sGenes, we computed *τ_s_*, a variation of the *τ* index^31^ based on sGene significance. We also employed the standard *τ* to determine the tissue specificity of sGene expression (see Methods). We found 469 genes under strong tissue-specific splicing regulation (highly tissue-specific sGenes), 81 of which did not display tissue-specific expression (Table S3). GO enrichment of these genes (universe: all sGenes) identified biological processes related to RNA processing and its regulation (three out of all five significant terms at FDR < 0.1: RNA splicing via transesterification reactions, regulation of RNA splicing, regulation of mRNA processing) suggesting again some mechanism of splicing autoregulation^26^.

### sQTLs are enriched in functional elements of the genome related to splicing and in high-impact variants

To shedlighton the mechanisms through which sQTLs may impact splicing, we built a comprehensive functional annotation of the human genome (see Methods). Overall, we observed a high density of functional elements in the proximity of sQTLs (Fig. S10). We next evaluated the enrichment of sQTLs in every functional category, with respect to a null distribution of similar variants not associated with splicing (Fisher’s exact test, FDR < 0.05). The top enrichments are summarized in Fig. 3a (the complete list, together with the statistical significance associated with each enrichment, is shown in Fig. S11).

As one would expect from bona fide variants affecting splicing, sQTLs are strongly enriched in splice sites (donors: OR = 12.98, adj. *p* value < 10^−16^; acceptors: OR = 12.23, adj. *p* value 1.22 · 10^−15^). They also display enrichments in exons, transcription factor (both activator and repressor) binding sites, RNA binding protein (RBP) binding sites, including several relevant splicing factors and spliceosomal components, and RNA Pol II sites. sQTLs tend to fall in open chromatin regions and show enrichments for several chromatin marks, particularly for H3K36me3 (OR = 2.85, adj. *p* value < 10^−16^). H3K27me3 regions, in contrast, are depleted of sQTLs (OR = 0.63, adj. *p* value < 10^−16^).

**Figure 3.**
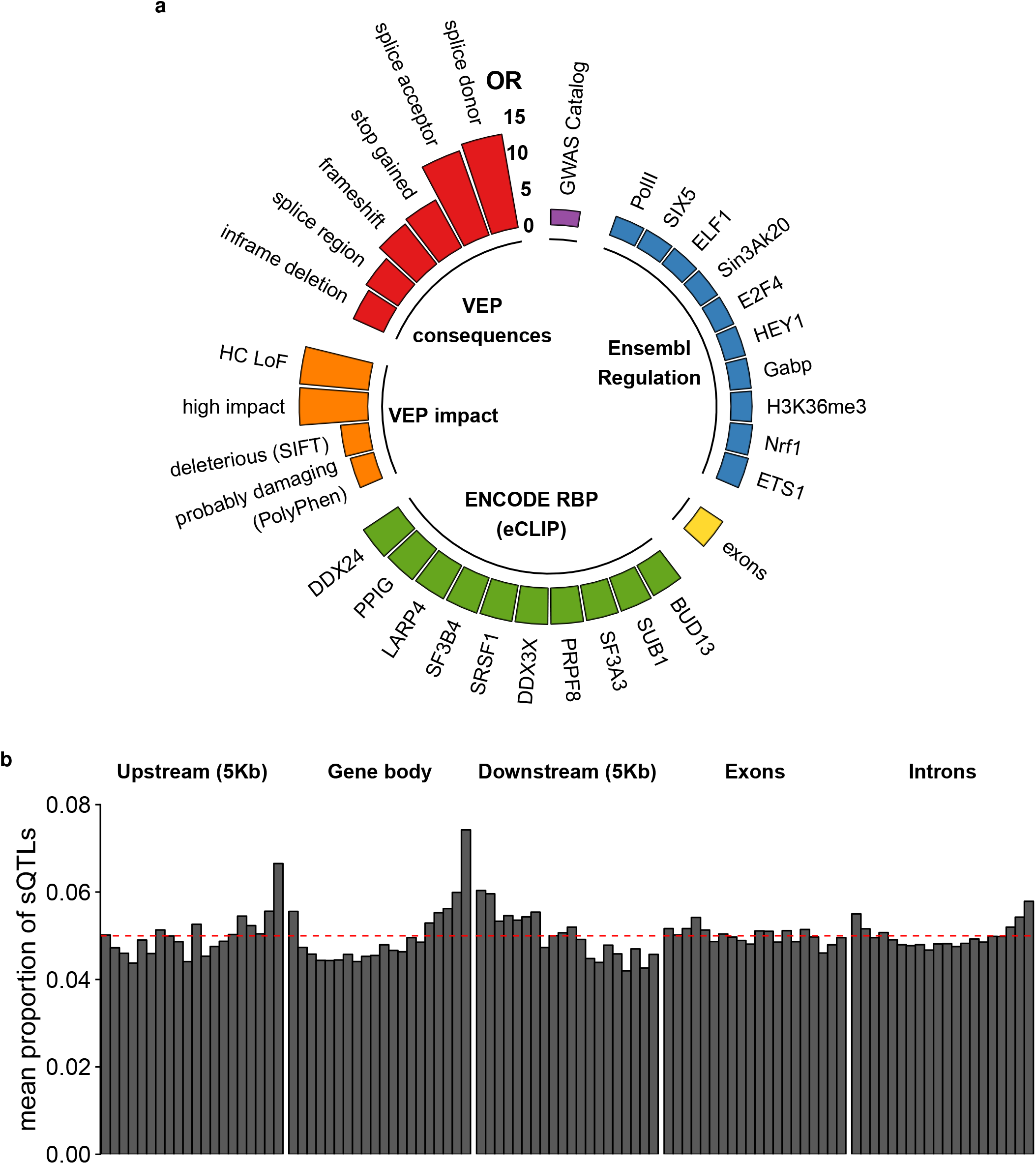
Functional enrichment and distribution of sQTLs. **a)** Top enrichments of sQTLs in functional annotations. The height of the bars represents the odds-ratio (OR) of the observed number of sQTLs to the expected number of variants that are not sQTLs overlapping a given annotation (see Methods): Variant Effect Predictor (VEP) categories (red) and impact (orange), ENCODE RNA-binding proteins (RBPs) eCLIP peaks (green), exons of GENCODE v19 protein coding and lincRNA genes (yellow), Ensembl Regulatory Build elements (blue) and GWAS catalog hits (purple). All these enrichments are significant at FDR < 0.05 and have OR confidence intervals not overlapping the range [1/1.50, 1.50]. **b)** Distribution of the mean proportion of sQTLs along the gene bodies of sGenes, their upstream and downstream regions, introns and exons. The red dashed line represents the expected distribution under a uniform model (see Methods).

sQTLs display large enrichments in predicted protein loss-of-function consequences (stop-gained, frameshift, VEP high impact variants, LOF-TEE high-confidence loss-of function variants (HC-LoF)) and potentially deleterious variants (according to Polyphen^32^ and SIFT^33^ scores). In addition, we found an enrichment in variants in high LD (*r*^2^ ≥ 0.80) with GWAS hits (OR = 2.08, adj. *p* value < 10^−16^). When performing stratified enrichments (see Methods), we found that large effect size sQTLs are more enriched in high impact variants, splice sites and GWAS hits, while small effect size sQTLs show larger enrichments in RBP binding sites, TFBS and open chromatin regions (Fig. S12).

In contrast to eQTLs, which tend to cluster around transcription start sites (TSS)^7,24^, we found sQTLs preferentially located towards transcription termination sites (TTS) (Fig. 3b), as previously observed^15^. In addition, while exonic sQTLs are uniformly distributed, intronic sQTLs are biased towards splice sites. Overall, sQTLs are closer to splice sites than non-sQTLs (Wilcoxon Rank-Sum test *p* value < 10^−16^, Fig. S13).

### sQTLs affect splice site strength and RNA-binding protein (RBP) binding

Enrichments in functional annotations (Fig. 3a) suggested several mechanisms through which sQTLs may affect splicing. One of them is the modification of splice site strength. Thus, for each variant within the sequence of an annotated splice site, we scored the site considering the reference and the alternative allele, using position weight matrices (see Methods). Overall, when compared to non-sQTL variants, a larger fraction of sQTLs modifies splice site strength (63% vs 49%, OR = 1.79, Fisher’s exact test *p* value < 10^−16^). The absolute difference in splice site strength is also larger for sQTLs (Wilcoxon Rank-Sum test *p* value 1.98 · 10^−7^), and increases with the sQTL effect size (Fig. 4a).

**Figure 4.**
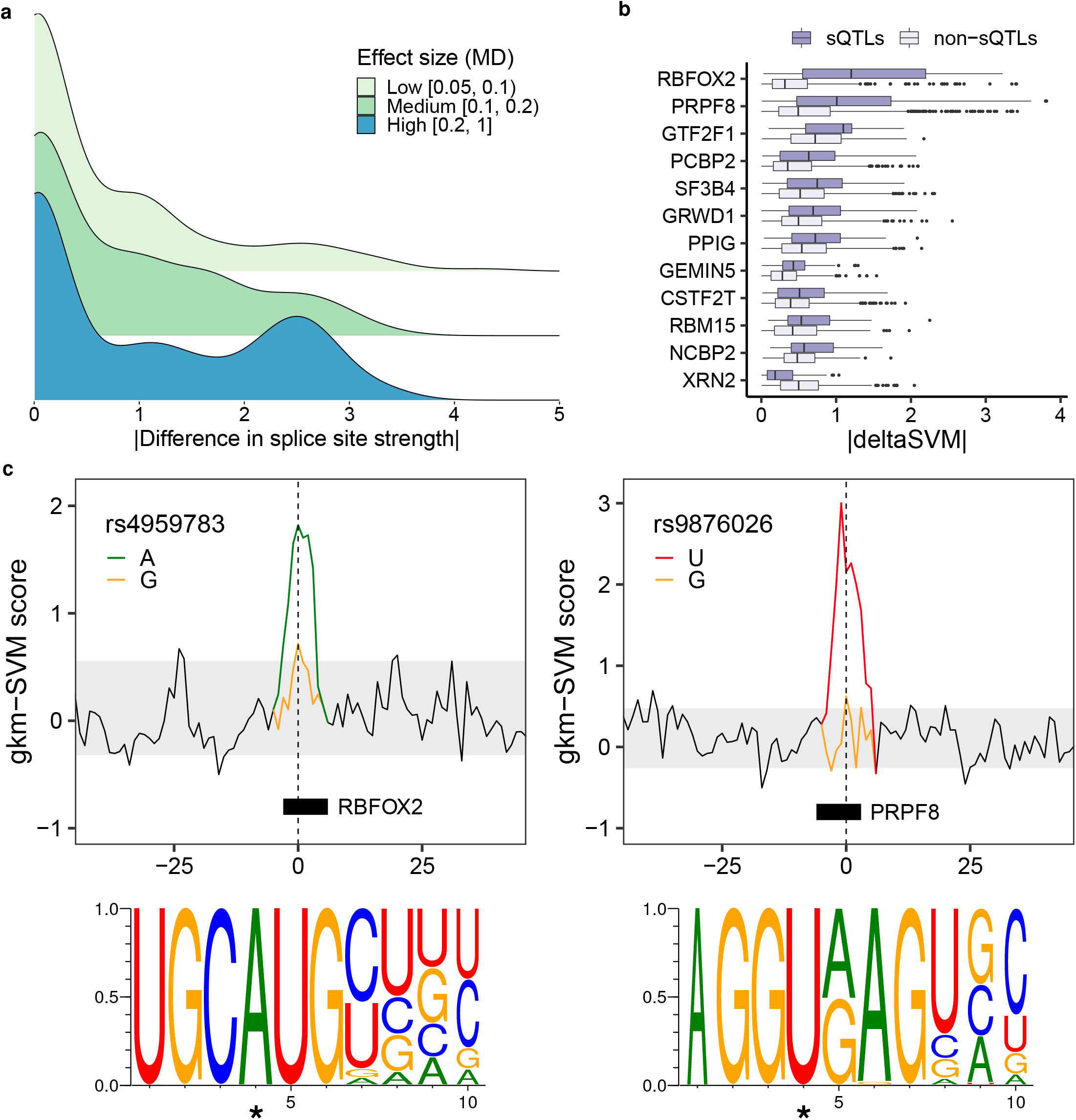
Impact of sQTLs on splice sites and RBP binding sites. **a)** Distribution of the absolute change in splice site strength for sQTLs with low, moderate and high effect sizes (MD value). **b)** Distribution of the absolute deltaSVM value (|deltaSVM|) of sQTLs and non-sQTLs, for RBPs with significantly different mean |deltaSVM| between sQTLs and non-sQTLs (FDR < 0.1). **c)** Modification of the binding sites of the RBPs RBFOX2 (left) and PRPF8 (right) by SNPs rs4959783 (chr6:3,260,093, G/A, |deltaSVM| = 2.48) and rs9876026 (chr3:11,849,807, T/G in the reverse strand, |deltaSVM| = 4.77), respectively. The lines represent the gkm-SVM scores of all possible (overlapping) 10-mers in a 100bp window around the SNP. Those corresponding to the 10-mers overlapping the SNP are colored according to the allele. SNP positions are marked with a dashed line. The grey area includes the 90% middle gkm-SVM scores of 10-mers not overlapping the variant. The relative location of the predicted RBP motifs and the corresponding sequence logos are also displayed. In the logos, the SNP position is marked with an asterisk.

Another mechanism through which sQTLs may affect splicing is the modification of RNA-binding protein (RBP) binding sites. To investigate it, we used eCLIP peaks of 114 RBPs available for HepG2 and K562 cell lines from the ENCODE project^34^. We employed a k-mer-based machine learning approach, which has been shown to outperform PWMs to identify transcription factor binding sites^35^ and provides a unique framework to assess the impact of genetic variants on the binding^36^. First we trained, for each RBP, a gapped k-mer support vector machine (gkm-SVM)^37^ on the sequences of high-confidence eCLIP peaks. 79 RBPs with a mean cross-validation ROC AUC ≥ 0.8 were kept. Then, we estimated the impact of all variants (whether sQTLs or not) overlapping the eCLIP peaks of each of these RBPs via the deltaSVM metric^36^, which measures the difference in predictive potential between the variant alleles (see Methods). To ensure the robustness of our results, we further restricted the analysis to RBPs with at least 30 sQTLs among the top 5% variants most predictive of the binding of the RBP at either allele, resulting in a final set of 32 RBPs (see Methods).

At FDR < 0.1, differences in |deltaSVM| between sQTLs and non-sQTLs were found significant for 12 RBPs (Fig 4b, the corresponding gkm-SVM ROC curves and motif logos shown in figures S14 and S15, respectively). Notably, for 11 of these proteins the |deltaSVM| values are larger for sQTLs than for non-sQTLs, as expected from variants regulating splicing. In addition, three of them (PPIG, SF3B4 and PRPF8) are among the top 10 RBPs whose binding sites are more enriched in sQTLs (Fig. 3a). In Fig. 4c, we show examples of the impact of the SNPs rs4959783 and rs9876026, which are sQTLs for the genes PSMG4 and TAMM41 (see also Fig. S16) and disrupt the binding sites of the RBPs RBFOX2 and PRPF8, respectively.

We further investigated whether allele-specific RBP binding (ASB) was occurring specifically at sQTLs. We obtained a set of ASB variants identified in the ENCODE eCLIP dataset using BEAPR (Binding Estimation of Allele-specific Protein-RNA interaction)^38^ and overlapped them with our sQTLs (see Methods). We found that sQTLs were highly enriched in ASB variants, when compared to non-sQTLs, across all RBPs (OR = 2.30, Fisher’s exact test *p* value < 10^−16^). When considering individual RBPs, at FDR < 0.05 we found a significant enrichment of sQTLs among ASB variants for 22 of them (Fig. S17), including 7 of the ones identified above with larger |deltaSVM| values for sQTLs. Altogether, these results suggest that sQTLs may affect splicing through allele-specific binding of RBPs.

Overall, the effect sizes (MD) of sQTLs in splice sites are larger than those of sQTLs overlapping RBP eCLIP peaks (Wilcoxon Rank-Sum test *p* value 1.98 · 10^−7^, Fig. S18), although the proportion of sQTLs in splice sites is much smaller (1.5% vs 8.3% out of all sQTLs). Often, both mechanisms may co-occur, as many RBPs bind near splice sites. This is the case of PRPF8, which binds specifically to the sequence of splice donors^39^. Indeed, the SNP rs9876026 (Fig. 4c), which modifies |deltaSVM| and has been identified as an allele-specific binding SNP for PRPF8 by BEAPR, also disrupts a donor splice site.

### sQTLs are preferentially located on post-transcriptionally spliced introns

Although splicing generally occurs co-transcriptionally (most introns are spliced prior to transcription termination and polyadenylation), there is a group of transcripts, often alternatively spliced, that tend to be processed more slowly, even post-transcriptionally^40^. We evaluated the role of genetic variants in the regulation of co- and post-transcriptional splicing (here referred to as cs and ps, respectively). In order to identify cs and ps introns, we determined the degree of splicing completion of annotated introns in nuclear and cytosolic RNA-seq data available for 13 cell lines from the ENCODE project (see Methods). We focused on a subset of introns consistently classified as eithercsor psin at least 10 of the analyzed cell lines (14,699 and 6,419 introns, respectively).

We observe a higher variant density in *ps* introns than in *cs* introns (4.38 vs 3.34 variants/Kb, differently distributed along the intron, Fig. S19a). The proportion of variants that are sQTLs in *ps* introns is larger than in *cs* introns (9.2% compared to 6.6%, OR = 1.47, Fisher’s exact test *p* value < 10^−16^). This enrichment is stronger when considering sQTLs that are not eQTLs for the same gene and tissue (OR = 1.67, *p* value < 10^−16^). Furthermore, sQTLs in ps introns display a substantial enrichment, with respect to sQTLs in cs introns, in RBPs and Pol II binding sites, and less markedly, in histone marks such as H3K36me3 and H3K4me3, open chromatin regions and TFBS (Fig. S19b). The proportion of sQTLs overlapping splice sites and GWAS hits is not significantly different between the two types of introns.

These results suggest that splicing regulation occurs preferentially at ps introns. This is expected, since these introns are retained longer within the primary transcript, offering more opportunities for regulation through the interaction with RBPs and other factors, including chromatin-related ones.

### sQTLs help to gain insight into disease and complex traits

To explore the relevance of regulatory variation affecting splicing in disease and complex traits, we assessed the overlap between GTEx sQTLs and the GWAS Catalog (https://www.ebi.ac.uk/gwas/), extended to include variants in high LD (*r*^2^ ≥ 0.80) with the GWAS hits. sQTLs display a substantial enrichment, when compared to non-sQTLs, in variants associated with a wide variety of GWAS traits and diseases (median OR = 3.23). Among the diseases with the largest sQTL enrichment, we find many for which alternative splicing has been previously related to their pathophysiology (Table S4). We integrated the enrichment information with estimates of semantic similarity between individual GWAS terms, computed from the Experimental Factor Ontology (EFO)^41^. Then, we applied multidimensional scaling (MDS) to summarize and represent the results (see Methods). This allowed us to identify the major phenotype groups related to sQTLs. Trait measurements (right-hand side of the MDS plot) and diseases (left-hand side) are the two main groups of enriched GWAS terms observed (Fig. 5a). Within the latter, we identify subgroups corresponding to cancer, autoimmune diseases and other disorders (neurological, cardiovascular, metabolic, etc.).

**Figure 5.**
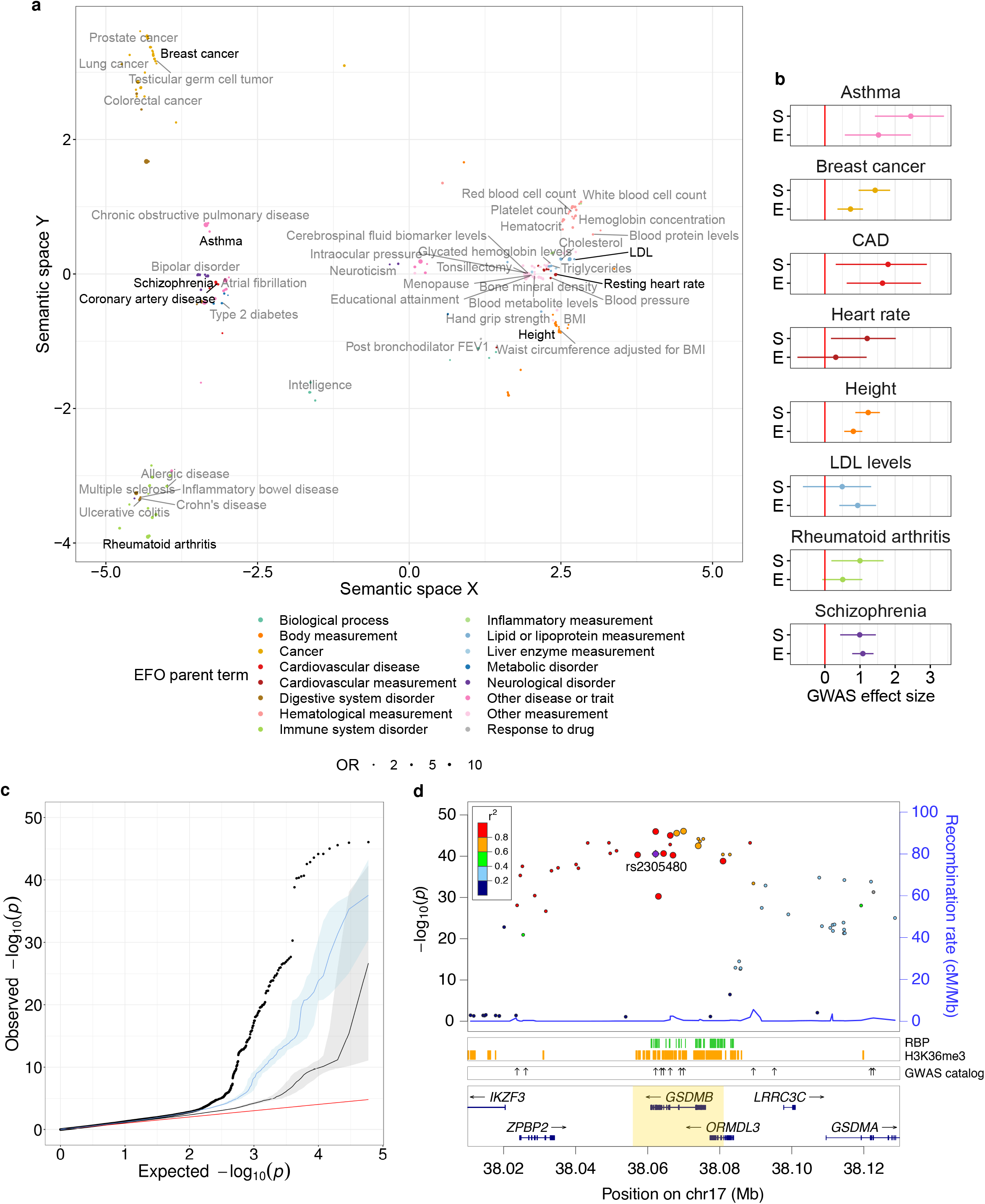
sQTLs and GWAS. **a)** Multidimensional scaling-based representation of the semantic dissimilarities between GWAS traits and diseases whose associated variants are enriched among sQTLs with respect to non-sQTLs (FDR < 0.05). Each GWAS term is represented by a dot, whose size corresponds to the enrichment odds-ratio (OR), and the color to the Experimental Factor Ontology (EFO) parent category the term belongs to. GWAS terms that lie close to each other are semantically similar. Eight representative traits with available summary statistics are highlighted. To help visualization, only the labels for the non-redundant, confidently and highly enriched terms are displayed (p value < 10^−8^, lower bound of the 95% confidence interval (CI) for the OR estimate > 1.5, width of the 95% CI for the OR estimate below the median). **b)** Maximum likelihood estimates and 95% CIs for the GWAS association effect size of variants affecting splicing (S) and expression, but not splicing (E), for eight traits and diseases. **c)** Quantile-quantile plot of *p* values for association with asthma in Demenais *et al*.^42^ for sQTLs (black dots), eQTLs without effects on splicing (blue), and variants with effects neither on expression nor on splicing (grey). Solid lines and coloured areas represent means and 95% CIs across 10,000 random samplings, respectively. The identity line is shown in red. **d)** *p* values for association with asthma in Demenais *et al*.^42^ (left y-axis) of variants in the region chr17:38,010,000-38,130,000, around the GSDMB gene (highlighted). The larger dots correspond to variants identified as sQTLs for the GSDMB gene in the lung. Linkage disequilibrium patterns (color-coded) and recombination rates are also displayed. The lower panels represent the location of RNA-binding protein (RBP) eCLIP peaks, H3K36me3 marked-regions and other GWAS-catalog associations with asthma (shown as arrows). The highlighted variant (rs2305480) is in perfect LD with rs11078928, previously shown to have an impact in GSDMB splicing^54^.

We also compiled genome-wide GWAS summary statistics for a subset of enriched traits representative of the observed clusters: asthma^42^, breast cancer^43^, coronary artery disease^44^, heart rate^45^, height^46^, LDL cholesterol levels^47^, rheumatoid arthritis^48^ and schizophrenia^49^. Overall, sQTLs show stronger GWAS associations than non-sQTL variants (Fig. S20). We further characterized the contribution to the disease phenotype of variants affecting splicing and variants affecting exclusively gene expression using fgwas^50^ (see Methods). For most of the traits analyzed, including asthma, breast cancer, coronary artery disease, heart rate, height and rheumatoid arthritis, we observe stronger effects among variants affecting splicing than among variants affecting only gene expression (Fig. 5b), suggesting that alterations in splicing play a relevant role in the molecular mechanisms underlying these traits. A few others, such as LDL levels or schizophrenia, display the opposite behaviour, pointing to a predominant effect of alterations in gene expression in the disease phenotype.

In addition, we observe that GWAS variants are especially enriched among sQTLs located in splice sites (OR = 2.66, Fisher’s exact test *p* value 1.02 · 10^−9^) or within RBP binding sites (OR = 1.78, Fisher’s exact test *p* value < 10^−16^). In particular, some of the traits and diseases with available summary statistics analyzed display stronger GWAS associations for sQTLs in RBP binding sites than for other sQTLs. Notably, this behaviour seems trait/disease and RBP-specific (Fig. S21).

An interesting example of how sQTL mapping can help to gain insight into the mechanisms underlying GWAS associations is the case of asthma and the gene gasdermin b (GSDMB). Asthma displays the largest effect size for sQTL variants (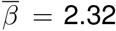, Fig. 5b), and stronger associations for sQTLs than for variants affecting only gene expression, or variants affecting neither expression nor splicing (Fig. 5c). Indeed we identified over 850 sQTLs co-localizing with known asthma loci, affecting the splicing patterns of genes related to immunity, including interleukins and immune cell’s receptors (IL13, TLSP, IL1RL1, TLR1), major histocompatibility complex components (HLA-DQA1, HLA-DQB1) or interferon-activated transcription factors (IRF1). However we also found other genes, such as GSDMB, with a priori less clear roles in the pathophysiology of the disease.

The GSDMB locus (17q21) has been consistently identified as a contributor to genetic susceptibility to asthma^42^ and other autoimmune diseases, such as type 1 diabetes^51^, ulcerative colitis^52^ or rheumatoid arthritis^53^. Although its exact function is unknown, GSDMB is highly expressed in human bronchial epithelial cells in asthma^54,55^, and it is known that overexpression of the human GSDMB transgene in mice induces an asthma phenotype^55^. In addition, the lipid-binding N-terminal domain of GSDMB and other gasdermins causes pyroptotic cell death^56^, potentially leading to the release of inflammatory molecules that trigger the asthma pathophysiology.

GSDMB is a sGene in 39 GTEx tissues, including lung (sGene FDR 1.42 · 10^−10^, median MD 0.22). Indeed, sQTLs for GSDMB are among the top associated variants with asthma in Demenais et al.^42^ (Fig. 5d). Allele C of the splice acceptor variant rs11078928 (chr17:38064469, T/C) has been shown to lead to the skipping of exon 6, which encodes 13 amino acids in the N-terminal domain, disrupting its pyroptotic activity^54^. While the major allele (T) is associated with a higher incidence of asthma, the C allele confers a lower asthma risk^54^. We have identified rs11078928 as an sQTL for GSDMB, whose alternative allele C precisely promotes expression of isoforms GSDMB-001 and GSDMB-002 (exon 6 skipping) vs isoform GSDMB-003 (exon 6 inclusion) (Fig. S22).

### Discussion

Using the unprecedented resource generated by the GTEx Consortium, we have obtained and analyzed a comprehensive set of genetic variants in the human genome affecting transcript isoform abundances (splicing QTLs, sQTLs). Unlike most methods for sQTL detection, we use a multivariate approach that monitors global changes in the relative abundances of a gene’s transcript isoforms, rather than targeting specific splicing events. Leveraging the correlated structure of isoform abundances is likely to result in increased power for sQTL mapping. Indeed, our approach has demonstrated the ability to detect sQTLs associated with complex splicing events that often escape univariate approaches^20^. In addition, we show that our method is not restricted to the analysis of transcript abundances, but can also accommodate other AS phenotypes, such as `LeafCutter` intron excision ratios^18^. A comparison of the resulting sQTLs obtained employing the two types of input data highlights the complementarity between global and local views of alternative splicing, especially regarding the types of splicing events identified^20,29^.

We have surveyed a large collection of tissues. Our analyses show thats QTLs tend to be highly shared, suggesting that there is a core set of variants that are involved in the regulation of splicing independently of the tissue or cell type. This has also been recently reported by the GTEx Consortium^12^. Among genes whose splicing is regulated by genetic variants (i.e. sGenes), there is a consistent enrichment of functions related to RNA processing, maybe reflecting splicing autoregulation. Indeed, several positive and negative autoregulation and cross-regulation mechanisms, such as coupling to nonsense-mediated decay, have been proposed for a large number of splicing factors^26^.

Overall, we found fewer genes regulated at splicing than at expression level. This is in line with recent reports^12^, and with the smaller contribution of splicing, compared to gene expression, to the global variability in transcript abundances across tissues and individuals^57,58^. Many variants, however, seem to be involved simultaneously in the regulation of both processes. This is not surprising, given that there is a substantial interplay between the molecular mechanisms underlying splicing and transcription, and because splicing often takes place co-transcriptionally^6^. In addition, variants altering splicing can affect RNA stability and, consequently, gene expression^59^.

In this regard, we have observed that introns that are spliced post-transcriptionally (ps) tend to be more enriched in sQTLs than introns that are spliced co-transcriptionally (cs). This is somehow expected, as ps introns are retained longer within the primary transcript, offering more opportunities for regulation. Consistent with this, sQTLs in ps introns display a larger enrichment, compared to sQTLs in cs introns, in RBP binding sites, but also in Pol II binding sites and histone marks. We note that chromatin-related features play a prominent role in co-transcriptional splicing, often through the regulation of transcription^6^. However, not-fully spliced but already 3’-end mature transcripts are present in the fraction of RNA attached to chromatin^60,61^. In this context, interactions between chromatin-side features and not-fully spliced transcripts can occur post-transcriptionally. Indeed, similar enrichments have been reported for exons that are spliced more slowly^40^. Overall, it seems that post-transcriptionally spliced introns play a larger role in splicing regulation that introns quickly spliced during transcription.

In addition to variants that are sQTLs and eQTLs for the same gene, we have found many variants that are sQTLs for a gene and eQTLs for a different one. In order to rule out indirect regulatory effects (e.g. when the variant directly affects the expression -splicing- of one gene, and the product of this gene directly affects the splicing -expression- of the other gene), we considered each effect (splicing or expression) occurring in different tissues, likely underestimating the number of such variants. Since our multivariate approach is not compatible with currently available co-localization methods (see below), we cannot distinguish the cases in which the two effects are indeed caused by the same variant or by two different variants in LD. Regardless of the underlying causal structure, these variants uncover regulatory loci, which we termed heteropleiotropic, that would be involved in the coordinated regulation, through different mechanisms, of different genes which otherwise do not appear to directly interact. Thus, heteropleiotropic loci could reveal regulatory relationships between genes that may not be easily captured by co-expression or splicing networks, highlighting the complexity of the gene regulation program in eukaryotes. While further work is required to establish the relevance and generality of this phenomenon, we believe that we have identified a number of convincing examples.

Our study also helps to understand the molecular mechanisms through which genetic variants impact splicing. Two such mechanisms appear to be the most relevant. On the one hand, direct impact on donor and acceptor splice sites. On the other hand, modification of binding sites of a wide variety of transcriptional regulators, especially RNA-binding proteins (RBPs), which are major players in RNA processing, transport and stability^5,62^. While the latter seems to occur in a larger number of cases, the former often leads to stronger effects on splicing. However, in many cases both mechanisms are likely to cooperate, given that RBPs often bind near splice sites.

Finally, our work provides new insights into the relationship between genetic variation, splicing and phenotypic traits. Specifically, we found that sQTLs are enriched in variants associated with a number of complex traits and diseases, some of them previously reported^9,10,14,15^. sQTLs display stronger GWAS associations than variants not associated with splicing and, for some traits, even larger effects than variants affecting exclusively gene expression. This grants splicing a key role in mediating the impact of genetic variation in human phenotypes^15^. Because gene expression is the main driver of biological function, we hypothesize that genetic variants affecting expression are likely to have a much larger biological impact than those affecting splicing: often, they could be lethal during development. In contrast, genetic variants affecting splicing may have subtler effects, therefore being better tolerated and leading more frequently to observable phenotypes. That genetic variants affecting splicing may underlay most human hereditary diseases has already been pointed out^19^. Especially relevant seems the implication of sQTLs in the mechanisms underlying autoimmune diseases, also supported by the overrepresentation of immune functions among sGenes. Actually, sQTLs have been recently proposed as relevant players in human immune response and its evolution^16^. In addition, sQTLs altering RBP binding seem to play a prominent role in disease. Indeed, the relevance of RBPs in human disorders has been often remarked^62^.

A more detailed analysis of the relationship between sQTLs and GWAS variants could be achieved by the usage of statistical methods to assess co-localization^63–65^, and subsequent fine-mapping of the sQTL candidates^66–68^ to assign causal probabilities. However, currently available methods are not directly applicable within our multivariate, non-parametric framework. In addition, recent works have demonstrated the utility of in silico splicing predictors to identify pathogenic variants affecting splicing, especially in the case of Mendelian disorders^69–71^. These methods provide a complementary view to RNA-seq-based approaches that measure splicing changes associated with genetic variants, such as `sQTLseekeR2`. Indeed, while the former target rare variants in the vicinity of splice sites with strong phenotypic effects, the latter focus on common regulatory variation, not restricted to the splice region nor necessarily pathogenic. Furthermore, the ability of pathogenicity predictors to account for features such as evolutionary conservation or exon importance provides valuable information about the relevance of individual alleles^71^, which may help prioritizing sQTLs in clinical settings.

Our implementation of an enhanced pipeline for sQTL mapping based on `sQTLseekeR2`, Nextflow and Docker will help sQTL discovery in multiple datasets, across different platforms, in a highly parallel and reproducible manner. Here we have employed it to identify sQTLs in the GTEx dataset. The extensive catalogue of sQTLs generated constitutes a highly valuable resource for the field. As our initial analyses already show, this resource will contribute to the understanding of the mechanisms underlying alternative splicing regulation and its implication in phenotypic traits, including disease risk.

## Data availability

All the data employed in this study is publicly available. GTEx data was obtained from dbGaP (www.ncbi.nlm.nih.gov/gap/, accessions *phs000424.v7.p2* and *phs000424.v8.p2*). ENCODE and ENTEx data was obtained from the ENCODE Portal (www.encodeproject.org, accession numbers provided in Supplementary Tables S5-7). The sQTL catalogue generated is available at https://public-docs.crg.es/rguigo/Data/dgarrido/sQTLs.GarridoMartin_etal.tar.gz.

## Code availability

Our pipeline for sQTL mapping is publicly available at https://github.com/dgarrimar/sQTLseekeR2-nf. Detailed information about the software can be found in Methods and Supplementary Note 1.

## Methods

### GTEx data

Transcript expression (transcripts per million, TPM) and variant calls (SNPs and short *indels*) were obtained from the V7releaseofthe Genotype-Tissue Expression (GTEx) Project (dbGaP accession *phs000424.v7.p2*). These correspond to 10,361 samples from 620 deceased donors with both RNA-seq in up to 53 tissues and Whole Genome Sequencing (WGS) data available. Metadata at donor and sample level and variant annotations (Ensembl’s Variant Effect Predictor, VEP, v83 (http://www.ensembl.org/info/docs/tools/vep) with the Loss-Of-Function Transcript Effect Estimator extension, LOFTEE, (https://github.com/konradjk/loftee)) were also retrieved. In GTEx V7, RNA-seq reads are aligned to the human reference genome (build hg19/GRCh37) using STAR^72^ v2.4.2a, based on the GENCODE v19 annotation (https://www.gencodegenes.org/human/release_19.html). Transcript-level quantifications are obtained with RSEM^73^ v1.2.22. WGS reads are aligned with BWA-MEM (http://bio-bwa.sourceforge.net) after base quality score recalibration and local realignment at known indels using Picard (http://broadinstitute.github.io/picard). Joint variant calling across all samples is performed using GATK’s HaplotypeCaller v3.4 (https://software.broadinstitute.org/gatk/documentation/tooldocs). Further details on GTEx data preprocessing and QC pipelines can be found on the GTEx Portal (https://gtexportal.org).

### sQTL mapping

#### Gene, transcript and variant filtering

48 tissues with sample size *n* ≥70 were selected for *cis* sQTL mapping. The *cis* window was defined as the gene body plus 5 Kb upstream and downstream the gene boundaries. We considered genes expressed ≥ 1 TPM in at least 80% of the samples (samples with lower gene expression were removed from the analysis of the gene), with at least two isoforms and a minimum isoform expression of 0.1 TPM (transcripts with lower expression in all samples were removed). These filters correspond to the default parameters of `sQTLseekeR2`. We analyzed only biallelic SNPs and short indels (autosomal + X) with MAF ≥ 0.01 and at least 10 samples per observed genotype group. In total, 3,588,609 variants and 16,010 genes (15,195 protein-coding, 815 lincRNA) were analyzed.

#### Covariate selection

To evaluate the impact of known technical and biological covariates at sample and donor level in expression data, we regressed the first ten principal components (PCs) of the gene expression per tissue onto each available covariate, determining the percentage of variance explained 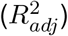. We selected donor ischemic time, gender and age, as well as sample RIN (RNA integrity number) as the most relevant covariates. We also included the first three genotype PCs (obtained from dbGap and computed as described in [24]), to control for population (i.e. ancestry) effects, and the genotyping platform employed (Illumina HiSeq 2000 or HiSeq X). Selected covariates were regressed out from the relative abundances of each gene’s transcript isoforms by `sQTLseekeR2` before testing for association with the genotype.

#### Software

For sQTL mapping we employed `sQTLseekeR2` v1.0.0, an enhanced version (see also Supplementary Note 1) of the `sQTLseekeR` R package^20^, which identifies genetic variants that are associated with multivariate changes in the relative abundances of agene’s transcript isoforms (i.e. splicing ratios). `sQTLseekeR2` was embedded in `sQTLseekeR2-nf`, a highly parallel, portable and reproducible pipeline for sQTL mapping developed using Nextflow^23^, a framework for computational workflows, and Docker container technology. `sQTLseekeR2` and `sQTLseekeR2-nf` are available at https://github.com/dgarrimar.

#### Details on significance assessment

We performed *cis* sQTL mapping on each tissue. Nominal *p* values were obtained using the function sqtl.seeker. To correct for the fact that multiple genetic variants in LD were tested pergene, an adaptive permutation scheme was applied (implemented in the function `sqtl.seeker.p`). A Benjamini-Hochberg false discovery rate (FDR) threshold of 0.05 was selected to identify sGenes. To retrieve all significant variant-gene pairs, we employed a procedure identical to the one described in [24] for expression QTLs (implemented in the function `sqtls.p`). See Supplementary Note 1 for details. In addition, as our test statistic is sensitive to the heterogeneity of the splicing ratios’ variability among genotype groups, a multivariate homoscedasticity test74 was also performed for each gene-variant pair. Pairs failing this test (FDR across all nominal tests > 0.05) were still assessed for significance and taken into account for multiple testing correction, but they were not reported as significant sQTLs.

#### Cell type heterogeneity

We employed xCell^25^ to estimate the enrichment of 64 reference cell types from the bulk expression profile of each GTEx sample. We applied the xCellAnalysis function in the xCell R package to the full gene expression TPM matrix (56,205 genes × 11,688 samples), in order to maximize tissue heterogeneity. We then applied the *τ* index^31^ (see also section *sQTL sharing*) to median xCell enrichments across samples per tissue. The cell type heterogeneity of a tissue was estimated as 1−*τ*. While these results should be interpreted with caution, as xCell is not a deconvolution method, but an enrichment method, they were generally biologically meaningful. For example, the most homogeneous tissues included brain subregions or transformed fibroblasts, and the most heterogeneous, spleen or whole blood. To determine the impact of the cell type heterogeneity of a tissue on sQTL discovery, we computed the partial correlation between the number of sGenes over the number of tested genes and the estimated cell type heterogeneity (i.e. 1−*τ*), controlling for the tissue sample size.

### sQTL effect size

We used the absolute maximum difference (MD) in mean adjusted transcript relative expression between genotype groups as a measure of the size of the effect. MD takes values in the interval [0, 1]. In practice, usual MD values belong to [0.01, 0.4]. As a general rule, we considered MD values < 0.1 as small effect sizes, MD values between 0.1 and 0.2 as moderate effect sizes and MD values greater than 0.2 as large effect sizes. sQTLs with MD values below 0.05 were not taken into account for further analyses (default in `sQTLseekeR2`).

### GO enrichment of sGenes

For each tissue, we obtained the corresponding set of sGenes, and performed hypergeometric tests to assess Gene Ontology (GO) Biological Process (BP) term over-representation, selecting as gene universe all the tested genes. We set a FDR threshold of 0.1 to identify significantly enriched terms. Similarly, we selected genes that were not sGenes in any tissue, and performed a hypergeometric test to assess GO BP term over-representation in this set (FDR <0.1, universe: all tested genes). Then, we employed REVIGO^75^ (http://revigo.irb.hr/, with parameters: allowed similarity = 0.9, database = *H. sapiens*, semantic metric = *SimRel*) to remove highly redundant terms and generate semantic similarity-based GO term representations for sGenes and non-sGenes.

### sQTL replication

To assess replication of GTEx sQTLs, we examined the *p* values for matched variant-gene pairs identified as splicing QTLs by `sQTLseekeR` for three immune cell types (CD14^+^ monocytes, CD16^+^ neutrophils, and naive CD4^+^ T cells) in the Blueprint Project (BP)^27^. Both studies have large differences in RNA sources (tissues in GTEx vs cell types in Blueprint), library preparation (unstranded polyA^+^ vs stranded Ribo-Zero), sequencing strategy (e.g. paired-end vs single-end in monocytes and neutrophils) and data processing pipelines (e.g. different transcript quantification software). *π*_1_ statistics, that provide an estimate of the proportion of true positives^76^, were computed for each pair GTEx tissue/BP celltype. A final replication rate for each GTEx tissue was calculated as the average *π*_1_ value across the three BP cell types.

### Alternative splicing events associated with sQTLs

To determine the nature of the splicing events related to sQTLs we selected, for each sQTL, the two isoforms of the target sGene that changed the most (in opposite directions) across genotypes. Then, we compared the exonic structure of both transcripts using the function `classify.events` of `sQTLseekeR`, which extends the classification proposed in AStalavista^77^. We considered the same event categories depicted in Monlong et al.20: exon skipping, alternative 5’ and 3’ splice sites, intron retention, mutually exclusive exons, alternative first and last exons, alternative 5’ and 3’UTR, tandem 5’ and 3’ UTRs, complex splicingevents(complexcombinations of events affecting internal exons) and complex 5’/3’ events (complex combinations of events affecting 5’/3’termini). Some of these events are not explicitly involving splicing, but alternative transcription initiation and termination sites. Note that each transcript pair, and therefore each sQTL, can be associated with more than one event.

### Heteropleiotropy and ENTEx histone modification analysis

Given a genetic variant *υ* and a pair of genes (i.e. *g*_1_ and *g*_2_) and tissues (i.e. *t*_1_ and *t*_2_), we consider *υ* heteropleiotropic with effects in different tissues if i) *υ* is an sQTL -but not an eQTL- for gene *g*_1_ in tissue *t*_1_, ii) *υ* is an eQTL -but not an sQTL- for gene *g*_2_ in tissue *t*_2_, iii) *υ* is neither an sQTL nor an eQTL for gene *g*_2_ in tissue *t*_1_ and iv) *υ* is neither an sQTL nor an eQTL for gene *g*_1_ in tissue *t*_2_. Out of 148,618 variants tested for association with both the expression and splicing of at least two genes in at least two tissues, we identified 6,552 heteropleiotropic cases. In order to evaluate whether changes at epigenetic level were occuring at these positions, we obtained ChIP-seq peaks corresponding to six histone modifications (H3K27ac, H3K4me1, H3K4me3, H3K36me3, H3K27me3 and H3K9me3) from the ENTEx data collection of the ENCODE Project^78,79^ (https://www.encodeproject.org/, accessed 2019-10-04, accession numbers provided in Table S5). ENTEx is a joint effort between GTEx and ENCODE consortia to deeply profile overlapping tissues from the same four donors (two male, two female) using shared technologies. The two tissues of interest were available for at least 3 out of 4 ENTEx donors for 2,855 heteropleiotropic variants. By overlapping these with the ChIP-seq peaks in the corresponding tissues, we identified 699 cases where one or more histone marks present in a tissue were absent in the other (in at least 3 donors). We compared this number with the one obtained for variants *υ′* affecting both the splicing and expression of the two genes (*g*_1_ and *g*_2_) in the two tissues (*t*_1_ and *t*_2_), using Fisher’s exact test for significance assessment.

### sQTL sharing

For every pair of tissues, we selected variant-gene pairs tested in both and found significant in at least one. We computed Pearson correlation (*r*) between their effect sizes (MD values). Tissue specificity was estimated as 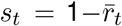, where 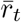 is the mean correlation between a given tissue t and the others. To determine the robustness of the observed sharing patterns with changes in the sample size, we randomly downsampled every original tissue dataset once to 100, 200 and 300 samples, ran our sQTL mapping pipeline again and re-evaluated the sharing patterns. Alternatively, we computed the Jaccard index on the sets of variant-gene pairs identified in every pair of tissues. In this case, tissue specificity estimates corresponded to 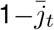, where 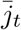 is the mean Jaccard index between a given tissue t and the others.

We further compared these approaches with a third strategy, aimed at evaluating the changes in the whole splicing phenotype due to sQTLs between differenttissues, ratherthan relying on MD values or sQTL presence/absence. This allows more flexibility, likely resulting in an increased ability to capture complex sharing patterns. In short, we focused on variant-gene pairs tested in all tissues and found significant in at least one. For every tissue *t_i_*, variant-gene pair *j* ∈ {1… *p*}, and genotype group *k* ∈ {0,1,2}, we computed the centroid of the adjusted (square root transformed, covariate corrected) splicing ratios, *c_t_i_jk_*. Then, we obtained:

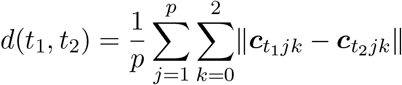

where *d* measures the distance between any two tissues (*t*_1_ and *t*_2_) in terms of sQTL sharing, as the mean (across variant-gene pairs) of the sum (across genotype groups) of the Euclidean distance between centroids (||**x**|| represents the Euclidean norm of vector **x**). Small values of d correspond to large sQTL sharing, and vice versa (Fig. S9a illustrates the behaviour of d for a single variant-gene pair evaluated in 4 tissues). A distance matrix built upon d values was then employed as input for hierarchical clustering.

To compare the tissue clusters obtained using different approaches we computed Baker’s Gamma (Γ), a metric of similarity between two dendrograms given by the rank correlation between the stages at which pairs of objects combine in each of the two trees^80^. Γ ranges from −1 to 1, with values close to 1 corresponding to high similarity between both dendrograms. To assess the significance of this similarity, we performed a permutation test (shuffling the labels of one tree 10,000 times, keeping tree topologies constant). We also employed Baker’s Gamma to compare ourtreeswith theone obtained using mashR30 for `LeafCutter` sQTLs in GTEx V812, available at https://github.com/broadinstitute/gtex-v8.

Of note, we employed pairwise approaches to study sQTL sharing, rather than methods to analyze QTL sharing jointly across tissues (such as mashR, to cite an example), given that the latter, to the best of our knowledge, cannot be applied in our multivariate, non-parametric setting.

In addition, for each sGene tested in all tissues and found significant in at least one, we determined tissue specificity of the sGene expression, using the *τ* index^31^:

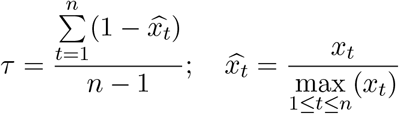

where *x_t_* is the expression of the gene in tissue *t* and *n* the number of tissues. *τ* takes values between 0 (i.e. genes equally expressed in all tissues) and 1 (i.e. tissue-specific genes). We calculated *τ* using median gene expression across tissues. In addition, to assess tissue specificity of splicing regulation, we computed a variation of *τ, τ_s_*, where *x_t_* was the −*log*_10_(FDR) of the sGene in tissue t. For sGenes in the top 20 percentile of the distribution of τs values, and the bottom 20 percentile of the distribution of *τ* values, we evaluated GO BP term over-representation (hypergeometric test, FDR < 0.1, universe: all sGenes).

### Functional enrichment of sQTLs

ChIP-seq peaks (transcription factor binding sites, histone marks) and open-chromatin regions were obtained from the Ensembl Regulation dataset (ftp://ftp.ensembl.org/pub/grch37/release-86/regulation/homo_sapiens). eCLIP peaks in HepG2 and/or K562 cell lines for 114 RNA-binding proteins (RBPs)^34^ were obtained from the ENCODE Project^78,79^ (https://www.encodeproject.org, see section *Splice site strength and sQTL impact on RBP binding sites* for details). Disease and complex-trait associated variants were retrieved from the GWAS catalog (https://www.ebi.ac.uk/gwas, accessed 2018-09-18), extended to GTEx variants in high linkage disequilibrium (*r*^2^ > 0.8) with the GWAS hits. Protein coding and lincRNA exons were derived from the GENCODE v19 annotation. The coordinates of these functional elements were overlapped with all the tested variants (either sQTLs or not) to obtain a functional annotation per variant. The functional consequences of each variant (stop-gained, frameshift, etc.), computed by the Variant Effect Predictor (VEP, http://www.ensembl.org/info/docs/tools/vep), were obtained from dbGap (accession *phs000424.v7.p2*). Note that the VEP leverages the Ensembl Variation dataset, which contains data from a wide variety of sources (https://www.ensembl.org/info/genome/variation/species/sources_documentation.html). From the VEP result we also identified variants with *HIGH* impact or in the categories *probably damaging* (PolyPhen, http://genetics.bwh.harvard.edu/pph2), *deleterious* (SIFT, https://sift.bii.a-star.edu.sg), *pathogenic* (ClinVar, https://www.ncbi.nlm.nih.gov/clinvar) and *high-confidence loss-of-function* (LOFTEE, https://github.com/konradjk/loftee).

The top 10 most significant sQTLs per gene and tissue were compared to a null distribution of 1000 sets of randomly sampled variants not associated with splicing (FDR > 0.05, *non-sQTLs*), with the same size of the sQTL set. The top 10 were selected to ensure the coverage of the less common annotations. Non-sQTLs were matched to sQTLs in terms of relative location within the gene and minor allele frequency (MAF). Specifically, we selected non-sQTLs so that they were located in the same bins (see section sQTL *location*) within the genes for which they were not sQTLs, as sQTLs within the genes for which they were sQTLs, and had MAFs equal to the sQTLs’ MAFs +/- 0.02. The enrichment was calculated as the odds ratio (OR) of the frequency of a certain annotation among sQTLs to the mean frequency of the same annotation across the 1000 non-sQTLs sets. To ensure enrichment reliability, we filtered out annotations with a mean frequency across the non-sQTLs sets lower than 5. The significance of each enrichment was assessed using a Fisher’s exact test. *p* values were corrected for false discovery rate, selecting a threshold of FDR < 0.05. Enrichments in a subset of relevant features, such as high impact/potentially damaging variants, splice sites, GWAS hits, exons, TFBS (all TFs pooled together), RBP binding sites (all RBPs pooled together), Pol II binding sites, HK36me3 and open chromatin regions, were also carried out separately for high effect size (MD ≥ 0.2) and low effect size sQTLs (MD < 0.1).

### sQTL location

We divided every sGene body into 20 bins of equal size and assigned each sQTL to the corresponding bin according to its location. The number of bins (20) was selected in order to provide a good balance between granularity and bin size. We computed the mean proportion of sQTLs (with respect to the total number of sQTLs for the gene) on each bin. An identical procedure was applied to exons, introns, downstream and upstream regions. In each case, to ensure a minimum bin size, we filtered out the 20% shortest regions. Under the null hypothesis of no preference in location, a uniform distribution for the mean proportion of sQTLs across bins was expected.

### Splice site strength and sQTL impact on RBP binding sites

To estimate the impact of genetic variants on splice sites, for each variant (either sQTL or not) within the sequence of an annotated splice site we scored the site considering the reference and the alternative allele, using position weight matrices (PWMs) built upon human splice sites^81^. High scores corresponded to common/strong splice sites, while low scores corresponded to rare/weak sites, probably leading to less efficient splicing. Then we estimated the change in splice site strength as the absolute value of the difference between alternative and reference scores.

To estimate the impact of genetic variants on RBP binding sites,we obtained eCLIP peaks in HepG2 and K562 cell lines for 114 RBPs^34^ from the ENCODE Project^78,79^ (https://www.encodeproject.org/, accessed 2018-04-16, accession numbers provided in Table S6). For each RBP, we selected the peaks significant at FDR < 0.01 and with a fold-change (FC) with respect to the mock input ≥ 2. We further required a minimum overlap between replicates (50% of the length of the union of a given pair of peaks). This constituted our positive set of RBP-binding sequences. We generated an equally-sized negative set of matched (in terms of GC content, length and repeats) sequences, not overlapping eCLIP peaks from the same RBP. We combined both sets of sequences to build our training set. To achieve feasible memory usage and running times, we limited the size of the training set to 30,000 sequences.

We then trained a gapped k-mer support vector machine (gkm-SVM)^37^ with default parameters (word length `l` = 10, informative columns `k` = 6), as recommended for our training set size range^36^. Other choices of `l` and `k` barely changed the overall performance (Fig. S23). The option `addRC` (add reverse complementary) was set to FALSE as we were working with RNA sequences. The classification performance was evaluated using a 5-fold cross-validation. 79 RBPs with a mean cross-validation area under the Receiver Operating Characteristic curve (ROC AUC) ≥ 0.8 were kept. To predict the impact of variants in RBP binding, for all the variants overlapping the eCLIP peaks (FDR < 0.01, FC ≥ 2) of a given RBP, we computed the deltaSVM metric^36^. The gkmSVM assigns a weight to each possible 10-mer, quantifying its contribution to the prediction of RBP binding. Each variant is given a score computed as the sum of the weights of the 10-mers overlapping it (10-mer SVM scores were used as a proxy for weights). deltaSVM computes the difference between the score of the alternative and the reference allele, quantifying their difference in predictive potential. Here we used the minor and the major allele instead of the alternative and the reference allele, respectively.

We focused on the most predictive variants of the binding of each RBP (score of the variant at either allele among the 5% highest scores for this RBP). This was done to target those variants lying on sequences likely to be highly relevant for RBP binding (i.e. potential binding sites). To ensure the robustness of our results, we further required at least 30 sQTLs with deltaSVM values per RBP, resulting in a final set of 32 RBPs. Of these, for 12 RBPs with significantly different |deltaSVM| values between sQTLs and non-sQTLs (Wilcoxon Rank-Sum test, FDR < 0.1), we obtained the 100 highest-scoring 10-mers, aligned them using mafft v7.407 (high accuracy mode *L-INS-1*)^82^, removed the columns of the alignment with more than 50% of gaps and built sequence logos using WebLogo standalone v3.6.0 (http://weblogo.threeplusone.com/).

To evaluate allele-specific RBP binding (ASB), we obtained the ASB variants identified in the same eCLIP dataset using BEAPR (Binding Estimation of Allele-specific Protein-RNA interaction), available from Yang et al.^38^. In short, BEAPR is a method to identify ASB events in protein-RNA interactions from eCLIP data. It accounts for crosslinking-induced sequence propensity and variability between replicates, outper-forming commonly used count-based approaches. We only considered ASB variants for which the same alleles had been genotyped in GTEx. We focused on sQTLs, non-sQTLs and ASB variants overlapping eCLIP peaks (FDR < 0.01, FC ≥ 2) for any of the 114 RBPs of interest in HepG2 and/or K562 cell lines. We assessed the significance of the difference in the proportion of sQTLs and non-sQTLs overlapping ASB variants across RBPs using Fisher’s exact test. We also performed this analysis separately for each RBP, using false discovery rate for multiple testing correction (FDR < 0.05).

### co- and post-transcriptional splicing

We obtained RNA-seq data from nuclear and cytoplasmic fractions (2 replicates/fraction) corresponding to 13 cell lines available from the ENCODE project^78^,79 (https://www.encodeproject.org/, accessed 2018-05-25, accession numbers provided in Table S7). A nextflow implementation of the Integrative Pipeline for Splicing Analyses (IPSA), developed in house (https://github.com/guigolab/ipsa-nf), was employed to determine the number of reads supporting splicing completion and splicing incompletion, for each intron annotated in GENCODE v19. We excluded from this analysis introns that overlapped either exons or non-identical introns in terms of chromosome, start and end positions. To assess the significance of the difference in the proportion of reads supporting splicing completion between nuclear and cytoplasmic compartments we employed Fisher’s exact test. False discovery rate was employed for multiple testing correction (FDR < 0.05). Introns with significantly larger proportions of reads supporting splicing completion in the cytoplasm were classified as post-transcriptionally spliced (here referred to as ps). Introns that did not pass the FDR threshold were labelled as either unprocessed (i.e. intron retention events) or co-transcriptionally spliced (here referred to as cs), depending on the degree of splicing completion in both cellular compartments. We focused on introns consistently classified as either ps or cs in at least 10 of the analyzed cell lines. We computed variant density (number of variants per Kb of intron) at 10 bins of equal size along both types of introns (10 was selected to ensure that enough variants were present in each bin). We also assessed the enrichment in functional elements of sQTLs in ps introns with respect to sQTLs in cs using Fisher’s exact test. False discovery rate was employed for multiple testing correction (FDR < 0.05).

### GWAS analyses

We downloaded the GWAS catalog, including the Experimental Factor Ontology (EFO) annotations for the GWAS terms (https://www.ebi.ac.uk/gwas, accessed 2018-09-18). We used LiftOver (https://genome.ucsc.edu/cgi-bin/hgLiftOver) to convert variant coordinates from hg38 to hg19 and PLINK v1.90b6.2 (https://www.cog-genomics.org/plink2) to extend the catalog to the variants in high linkage disequilibrium (*r*^2^ ≥ 0.8) with the GWAS hits. The sQTL enrichment was calculated as the odds ratio (OR) of the frequency of GWAS variants among sQTLs to the mean frequency of GWAS variants across 1000 matched non-sQTL sets (see section Functional enrichment of sQTLs). In parallel, we obtained the complete EFO ontology (https://www.ebi.ac.uk/efo/) in Open Biomedical Ontologies (OBO) format. For the GWAS terms with an OR > 1, we used the `ontologySimilarity` R package^83^ to compute the pairwise semantic similarity (method = resnik) between the enriched GWAS terms, and built a similarity matrix, *S*. From it, we derived a distance matrix, *D*, as *max*(*S*) – *S*, and performed multidimensional scaling (MDS). This is an analogous strategy to the one employed in REVIGO75 to visualize GO terms.

We further compiled genome-wide GWAS summary statistics for 8 traits representative of the clusters observed in the MDS representation: asthma42, breast cancer43, coronary artery disease44, heart rate45, height46, LDL cholesterol levels47, rheumatoid arthritis48 and schizophrenia^49^. In each case, we employed fgwas50 v0.3.6 (https://github.com/joepickrell/fgwas, default parameters, except for window size set to 2500bp to ensure convergence) to obtain the maximum likelihood estimate and 95% confidence interval for the association effect size, both for i) sQTLs (variants affecting splicing, independently of their effect on expression), and ii) variants affecting expression, but not splicing (GTEx V7 eQTLs tested also in our setting and not identified as sQTLs). To display the regional GWAS association results for the GSDMB locus we employed LocusZoom standalone v1.4 (https://github.com/statgen/locuszoom-standalone).

## Acknowledgements

We thank the donors and their families for their generous gifts of organ donation for transplantation and tissue donations for the GTEx research study. We thank Manuel Muñoz, Emilio Palumbo, Aliaksei Holik, François Aguet and Kristin Ardlie for useful discussions. We thank Romina Garrido for administrative support. This project was supported by the National Human Genome Research Institute of the National Institutes of Health under grants R01MH101814 and 5U24HG009446, as well as by the BIO2015-70777-P grant from the Spanish Ministry of Economy, Industry and Competitiveness (MEIC). The Genotype-Tissue Expression (GTEx) project was supported by the Common Fund of the Office of the Director of the National Institutes of Health (http://commonfund.nih.gov/GTEx). D.G-M. is supported by a ‘la Caixa’-Severo Ochoa pre-doctoral fellowship (LCF/BQ/SO15/52260001). B.B. is supported by the fellowship 2017FI_B 00722 from the Secretaria d’Universitats i Recerca del Departament d’Empresa i Coneixement (Generalitat de Catalunya) and the European Social Fund (ESF). We thank the ENCODE Consortium and, in particular, Thomas Gingeras’, Gene Yeo’s and Bradley Bernstein’s laboratories for data production. We also acknowledge support of the Spanish Ministry of Economy, Industry and Competitiveness (MEIC) to the EMBL partnership, ‘Centro de Excelencia Severo Ochoa’, the CERCA Programme / Generalitat de Catalunya and the European Regional Development Fund (ERDF).

## Author Information

### Contributions

D.G-M. and R.G. conceived and designed the study. D.G-M. implemented the software and analyzed the data. B.B. contributed to several analyses, provided analysis tools and helped with the interpretation of the results. M.C. and F.R. contributed ideas and statistical advice, helping with the design of the software. D.G-M. and R.G. wrote the original draft. All the authors reviewed the final manuscript.

