## Supplementary material for "Identification and analysis of splicing quantitative trait loci across multiple tissues in the human genome": supp.notes1-3.pdf

### Supplementary Note 1

#### sQTLseekeR and sQTLseekeR2

`sQTLseekeR`[1] identifies genetic variants that are associated with changes in the relative abundances of a gene's transcript isoforms (i.e. splicing ratios). The splicing ratio of isoform $i$ in individual $j$ is $f_{ij} = x_{ij}/\sum_{i=1}^{q} x_{ij}$, where $x_{ij}$ is the expression of isoform $i$ in individual $j$ and $q$ is the number of isoforms of the gene. Note that splicing ratios configure a multivariate phenotype, with as many values as there are transcripts for a given gene. `sQTLseekeR` uses the Hellinger distance between splicing ratios to estimate their variability across individuals. The Hellinger distance between the splicing ratios of individuals $j$ and $k$ is given by:

$$d_H(j,k) = \sqrt{\sum_{i=1}^{q}(\sqrt{f_{ij}} - \sqrt{f_{ik}})^2}$$

For a given gene and genetic variant, `sQTLseekeR` compares the variability of the gene's splicing ratios, calculated as a sum of squares, within and between genotypes at the variant (i.e. 0, 1, 2). The comparison is performed computing a pseudo F score as defined by Anderson[2]:

$$\tilde{\mathrm{F}} = \frac{SS_B}{SS_W} = \frac{SS_T - SS_W}{SS_W}$$

where $SS_T$ is the total variability, $SS_B$ is the variability between genotypes, and $SS_W$ the variability within genotypes. If $N$ is the total number of individuals,

$$SS_T = \frac{1}{N}\sum_{j=1}^{N-1}\sum_{k=j+1}^{N} d_H^2(j,k)$$

where $d_H^2(j,k)$ is the squared Hellinger distance between the splicing ratios of individuals $j$ and $k$, and

$$SS_W = \sum_{g=1}^{p}\frac{1}{n_g}\sum_{j=1}^{N-1}\sum_{k=j+1}^{N} d_H^2(j,k)\epsilon_{g,j,k}$$

where $p$ is the number of genotypes at the variant, $n_g$ the number of individuals with genotype $g$, and $\epsilon_{g,j,k} = 1$ if individuals $j$ and $k$ have genotype $g$ at the variant, otherwise $\epsilon_{g,j,k} = 0$.

To assess significance, `sQTLseekeR` relies on the asymptotic distribution of pseudo F scores. Anderson derived the asymptotic distribution for the numerator of the pseudo F score, which is a linear combination of independent $\chi^2$ variables with $n_g-1$ degrees of freedom, where the coefficients are derived from the matrix built upon the distances between every pair of individuals[3]. For any set of permutations of a given data set, the sum of squares of the numerator is a monotonic function of the pseudo F score, thus the permutation *p* values computed on the numerator and on the pseudo F score are equivalent. Essentially, this approach is a nonparametric analogue to multivariate analysis of variance (MANOVA). `sQTLseekeR2`, available at `https://github.com/dgarrimar/sQTLseekeR2`, is a completely rewritten, largely enhanced version of `sQTLseekeR` (see below), although the statistical framework implemented remains the same. Notably, this framework is not necessarily restricted to the analysis of relative transcript abundances, and can be applied to other multivariate AS phenotypes (see also Supplementary Note 2).

#### Enhancements in sQTLseekeR2

##### *p* value computation

To approximate the asymptotic distribution of the test statistic, `sQTLseekeR` relied on the Monte Carlo generation of $\chi^2$ distributed values. Therefore, the smallest achievable *p* value was $1/(M+1)$, where $M$ was the number of Monte Carlo generations. Due to memory and time limitations, in practice $M$ takes values up to $10^7$. In contrast, `sQTLseekeR2` relies on Davies algorithm[4] to perform this task. The algorithm allows to approximate the CDF of the asymptotic distribution of the test statistic with high accuracy through the numerical inversion of the characteristic function. This has considerably decreased the *p* value precision limit (up to $10^{-14}$) and speeded up the *p* value computation.

##### Correction for potential confounders

`sQTLseekeR2` allows the incorporation of potential technical or biological confounders (either numerical or categorical) as covariates to be regressed out from the splicing ratios before testing for association with the genotype.

##### Multiple testing correction scheme

In *cis* sQTL mapping, all possible variant-gene pairs in *cis* are tested for association. This implies two multiple-testing levels: i) multiple variants are tested per gene and ii) multiple genes are tested genome-wide. To account for (i), `sQTLseekeR2` implements a permutation scheme that empirically characterizes, for each gene, the distribution of nominal *p* values expected under the null hypothesis of no association[5]. This null distribution is then modeled using a beta distribution as in `FastQTL`[6]. In short, the minimal nominal *p* value ($p_{min}$) per gene is used as the test statistic. Permutations are performed by randomizing sample labels for the splicing ratios. By default, a maximum of 1000 permutations is performed, with a stopping criteria of having at least 100 permuted $p'_{min}$ values lower than the nominal $p_{min}$. A maximum likelihood-fitted beta distribution from the permutations is used to compute gene-level empirical *p* values. To account for (ii) and identify sGenes, `sQTLseekeR2` uses false discovery rate (FDR). Due to the computational burden of the permutations, a simpler approach (originally in `sQTLseekeR`) based on performing FDR on pooled nominal *p* values is also available. This can be further enhanced in `sQTLseekeR2` by clustering variants in high linkage disequilibrium (see section *Linkage disequilibrium-based variant clustering*).

To identify all significant variant-gene pairs, `sQTLseekeR2` implements a procedure identical to the one depicted in [7] for expression QTLs: i) it defines a genome-wide empirical *p* value threshold, $p_t$, as the gene-level *p* value closest to the 0.05 FDR threshold; ii) for each gene, it computes a nominal *p* value threshold based on the beta distribution model of the minimum *p* value distribution $f(p_{min})$ (obtained from the permutations for this gene, see above), as $F^{-1}(p_t)$ (where $F^{-1}$ is the inverse cumulative distribution); and iii) for each gene, variants with a nominal *p* value below the gene-level threshold are considered significant.

##### Linkage disequilibrium-based variant clustering

A substantial fraction of the variants tested in *cis* (up to 50% for a window of 5 Kb around the gene) are in high ($r^2 \geq 0.8$) or even complete ($r^2 = 1$) linkage disequilibrium (LD). `sQTLseekeR2` allows clustering variants above a user-defined LD threshold, so that only a representative of each cluster is tested for association. This reduces the number of dependent tests and therefore the running time and stringency of the multiple testing correction. It can be combined with simple FDR as a fast, suboptimal alternative to permutations for multiple testing correction.

##### Reduced running time

The more efficient *p* value computation, along with other minor improvements to speed up the preprocessing steps, allowed to achieve a substantial reduction (between 3 and 4 times) of the running time of a nominal pass of `sQTLseekeR2` with respect to `sQTLseekeR` (Fig. S24).

#### Simulation study

To study the type I error of `sQTLseekeR2`, we simulated the splicing ratios of 10,000 genes with $q$ transcript isoforms in $n$ individuals, under the null hypothesis of no association with genetic variants. We evaluated different scenarios modifying the values of $q \in \{5, 10, 15\}$ (selected matching the distribution of the number of splicing isoforms per gene considered in our sQTL analysis) and $n \in [100, 500]$ (selected matching the tissue sample sizes in GTEx V7). We simulated 1 SNP per gene, the probability of an individual belonging to each genotype group (i.e. 0, 1, 2) being 0.6, 0.3 and 0.1, respectively. Splicing ratios were simulated as vectors of proportions in the $q-1$ simplex with mean $\boldsymbol{c} = \boldsymbol{q}^{-1}$ (i.e. the center of the simplex) for all genotype groups. To generate observations in the simplex with certain variability around $\boldsymbol{c}$ we performed $q$ random displacements of size $\sim \mathcal{N}(0, \sigma^2)$ from $\boldsymbol{c}$ towards the simplex vertices. $\sigma^2$ was selected so that the mean standard deviation of splicing ratios was constant and approximately equal to 0.03 across different values of $q$, ensuring that the resulting vectors of proportions were elements of the simplex. We estimated type I error for each combination of $q$ and $n$ as the fraction of tests (out of 10,000) found significant at $\alpha = 0.05$. Our results show that `sQTLseekeR2` presents overall a controlled type I error rate (Fig. S25a).

To study power, we simulated splicing ratios under the alternative hypothesis. First, we generated vectors of proportions in the simplex as above. Then we incremented the splicing ratio of the first transcript isoform in an amount $\Delta$ (decreasing accordingly the splicing ratios of the other transcript isoforms in $\Delta/(q-1)$) for individuals with genotype 0 at the SNP of interest. We did the opposite for individuals with genotype 1 at this SNP. We explored values of $\Delta$ in the range (0, 0.02]. In practice, $\Delta =$ MD/2, where MD is the sQTL effect size (see MD definition in Methods). To estimate power, for each combination of $n$, $q$ and MD, we computed the fraction of tests significant at $\alpha = 0.05$. Our results show that `sQTLseekeR2` presents a very large power to detect differences in splicing ratios across genotype groups (Fig. S25b), even when effect sizes are very small (e.g. with $q = 10$ and $n = 300$, MD $= 0.02$ is detected with power 0.92).

##### Nextflow pipeline

sQTL mapping in large sequencing projects such as GTEx, with a large number of samples, a high density genotyping and millions of statistical tests, requires a parallelization strategy to absorb such a computational load in reasonable running times. However, any given implementation of an sQTL mapping pipeline may be restricted to a particular computing platform, hindering reproducibility, portability and scalability. Nextflow is a domain-specific language (DSL) for parallel computational pipelines[8]. It allows to execute a pipeline on multiple platforms without changes, and supports container technologies such as Docker (`https://www.docker.com/`) or Singularity (`https://singularity.lbl.gov/`), ensuring reproducibility. Taking advantage of these features, we have developed an sQTL mapping pipeline using `sQTLseekeR2`, Nextflow and Docker, in order to map sQTLs in GTEx. Our pipeline, named `sqtlseeker2-nf`, is available at `https://github.com/dgarrimar/sqtlseeker2-nf`.

### Supplementary Note 2

#### LeafCutter

A number of studies have recently employed `LeafCutter` to identify sQTLs[1–3]. `LeafCutter`[4] is a method to quantify alternative splicing (AS) from short-read RNA-seq data. It uses split-mapped reads to identify alternatively-excised introns and groups them into clusters. Then, intron usage can be expressed as proportions (i.e. intron excision ratios: the number of reads supporting an intron over the total number of reads supporting the cluster to which the intron has been assigned). To test for association with genetic variants, individual intron excision ratios are often modeled using univariate linear regression (e.g. as implemented in `FastQTL`[5]).

The vector of intron excision ratios obtained by `LeafCutter`, however, can also be modeled as a multivariate phenotype, and therefore it can be employed as input for `sQTLseekeR2` to identify sQTLs. Actually, multivariate modeling through a Dirichlet-multinomial generalized linear model (GLM) is available within `LeafCutter`, but only for differential splicing analyses. Given the popularity of `LeafCutter`, we have run `sqtlseeker2-nf` with `LeafCutter` quantifications in GTEx V7, and compared the resulting sQTLs with the set obtained using `RSEM` transcript quantifications.

#### LeafCutter AS quantification and sQTL mapping

We quantified AS based on the intron excision phenotypes defined by `LeafCutter` as follows: first, we used the `bam2junc.sh` script to quantify intron usage and then, we employed the `leafcutter_cluster.py` script (with options `--min_clu_reads 30 --min_clu_ratio 0.001 --max_intron_len 500000`) to define intron clusters. Both scripts are provided with `LeafCutter` software. To map `LeafCutter` clusters to genes, we employed the `map_clusters_to_genes.R` script available from `https://github.com/broadinstitute/gtex-pipeline`, with exon coordinates derived from GENCODE v19.

The resulting `*_perind_numers.counts.gz` files were modified to include two ID columns: i) *intron:cluster:gene* and ii) *cluster:gene*, and used as input for `sqtlseeker2-nf`, in place of tissue transcript expression files. Note that these columns replace the transcript and gene ID columns required when using transcript quantifications. By setting `--min_transcript_expr 5 --min_gene_expr 30` in `sQTLseekeR2`, we considered clusters supported by $\geq$ 30 reads in at least 80% of the samples (samples with lower cluster read support were removed from the analysis of the cluster), with at least two introns and a minimum intron read count of 5 (introns with lower read support in all samples were removed). The remaining settings (covariate correction, variant filtering, FDR threshold, etc.) were identical to the run using transcript quantifications (see section *sQTL mapping* in Methods). Note that for each cluster, the *cis* window was defined as the corresponding gene body plus 5 Kb upstream and downstream the gene boundaries. In total, 2,507,537 variants and 14,045 genes (13,347 protein coding, 698 lincRNA) were analyzed. At a 0.05 false discovery rate (FDR), we found a total of 263,827 *cis* sQTLs affecting 7,471 sGenes (7,046 protein coding genes and 425 lincRNAs) (Table S8).

#### Comparison between sQTLs identified using RSEM transcript quantifications and Leafcutter intron excision ratios

To obtain comparable results, genes that were not tested in both runs were filtered out, and the multiple testing correction (process `permuted_mtc` in `sqtlseeker2-nf`) was repeated on the common set of genes. In addition, the selection of the same *cis* window and parameters regarding variant filtering in both runs ensured that the set of variants tested was virtually identical. We compared the pairs sQTL-sGene identified by i) both approaches (i.e. common sQTLs), ii) `LeafCutter` quantifications only (i.e. LC-exclusive sQTLs), and iii) transcript quantifications only (i.e. TQ-exclusive sQTLs). Note that in the case of `LeafCutter`, we considered that an sQTL affects a given gene if it is associated with changes in intron excision ratios of at least one of the intron clusters assigned to this gene.

Overall, we observed a moderate overlap between the sQTL sets identified by the two approaches (median *Jaccard* index of sGenes across tissues 0.34, median $\pi_1$ of TQ in LC sQTL-sGene associations across tissues 0.73, Fig. S26a). The number of LC-exclusive sQTLs was consistently larger than the number of TQ-exclusive sQTLs. This could be explained by the fact that `LeafCutter` is able to detect novel introns, whereas transcript quantification relies only on GENCODE v19 annotation. Indeed, 44% (median across tissues) of the most-changing introns within intron clusters associated with LC-exclusive sQTLs are novel. However, we observed that the proportion of LC-exclusive sGenes decreased with the tissue sample size, while the proportion of common and TQ-exclusive sGenes increased (Fig. S26b). Furthermore, LC-exclusive sQTLs are associated with intron clusters with lower read support, when compared to common sQTLs (Fig. S26c). Hence, these could also indicate an inflated false positive rate associated with `LeafCutter` quantifications, which has already been observed in differential splicing analyses[6]. For large sample sizes, the number of sQTLs identified by `sQTLseekeR2` using `LeafCutter` or RSEM quantifications is very similar.

We also compared sQTL effect sizes (MD values). In the case of `LeafCutter` sQTLs, MD values (MD$_{LC}$) correspond to the absolute maximum difference in mean adjusted intron excision ratios of a given intron cluster (rather than in mean adjusted transcript relative expression, MD$_{TQ}$, as when using transcript quantifications) between genotype groups. In the case of sGenes with more than one significant intron cluster, sQTL effect sizes were computed as the median MD$_{LC}$ value across all significant intron clusters. As expected, MD$_{TQ}$ values were smaller (median reduction across tissues 7%) for TQ-exclusive sQTLs than for common sQTLs. The same behaviour was observed for the MD$_{LC}$ values of LC-exclusive sQTLs and common sQTLs, although in this case the reduction was much larger (median reduction across tissues 22%). This suggests that `LeafCutter` quantifications allow to identify smaller effects, and contributes to explain the larger number of LC-exclusive sQTLs with respect to TQ-exclusive sQTLs found. In addition, despite the marked differences in the splicing phenotypes employed, the effect sizes of common sQTLs obtained by `LeafCutter` (MD$_{LC}$) and transcript quantifications (MD$_{TQ}$) displayed a substantial *Pearson* correlation (median $r$ across tissues of 0.51).

Given that `LeafCutter` does not provide information about the flanking exons, we could not characterize the AS events associated with LC-exclusive sQTLs. Nevertheless, we computed the AS events associated with TQ-exclusive and common sQTLs (see section *Alternative splicing events associated with sQTLs* in Methods) and compared them. We observed that the nature of the AS events identified was different between TQ-exclusive and common sQTLs (Fig. S26d). We further explored individual events using the Wilcoxon Rank-Sum test to assess the significance of the differences. We found that common sQTLs displayed larger proportions of events involving internal exons, including mutually exclusive exons ($p$ value $6.70 \cdot 10^{-10}$) or alternative acceptor ($p$ value $4.51 \cdot 10^{-5}$). In contrast, TQ-exclusive sQTLs showed larger proportions of AS events affecting the gene termini, such as alternative first exon ($p$ value $1.56 \cdot 10^{-5}$) or alternative 3' UTR ($p$ value $4.74 \cdot 10^{-4}$), as well as intron retention ($p$ value $7.51 \cdot 10^{-6}$), an event that cannot be identified by `LeafCutter`. This highlights the complementarity of `LeafCutter` and transcript quantifications for alternative splicing measurement in the context of sQTL mapping.

Finally, we computed sQTL location (see section *sQTL location* in Methods) for both LC-exclusive and TQ-exclusive sQTLs. Overall, both types of sQTLs displayed a similar distribution along exons, introns and upstream/downstream regions. However, we found a slightly more marked enrichment of *TQ*-exclusive sQTLs towards the transcription termination site (Fig. S26e). This may relate to the larger proportion of AS events affecting the gene termini associated with TQ-exclusive sQTLs.

### Supplementary Note 3

#### sQTL mapping in GTEx V8

Transcript expression (transcripts per million, TPM) and variant calls (SNPs and short *indels*) were obtained from the V8 release of the GTEx Project (dbGaP accession *phs000424.v8.p2*). These correspond to 15,253 samples from 838 deceased donors with both RNA-seq in up to 54 tissues and Whole Genome Sequencing (WGS) data available. In GTEx V8, RNA-seq reads are aligned to the human reference genome (build hg38/GRCh38) using STAR[1] v2.5.3a, based on the GENCODE v26 annotation (`https://www.gencodegenes.org/human/release_26.html`). Transcript-level quantifications are obtained with RSEM[2] v1.3.0. WGS reads are aligned with BWA-MEM (`http://bio-bwa.sourceforge.net`) after base quality score recalibration and local realignment at known *indels* using Picard (`http://broadinstitute.github.io/picard`). Joint variant calling across all samples is performed using GATK's HaplotypeCaller v3.5. (`https://software.broadinstitute.org/gatk/documentation/tooldocs`). Further details on GTEx data preprocessing and QC pipelines can be found in [3].

49 tissues with sample size $n \geq 70$ were selected for *cis* sQTL mapping (48 already present in the V7 analysis, plus *Kidney Cortex*). Gene, transcript and variant filtering was performed as in V7 (see Methods). In total, 4,074,385 variants and 16,202 genes (15,319 protein coding, 883 lincRNA) were analyzed. Analogously, the covariates selected were donor ischemic time, gender and age, sample RIN (RNA integrity number), five genotype PCs (see [3] for details), and the WGS platform (Illumina HiSeq 2000 or HiSeq X) and library construction protocol (PCR-based or PCR-free) employed. We performed sQTL mapping on each tissue using `sqtlseeker2-nf`.

At a 0.05 false discovery rate (FDR), we found in GTEx V8 a total of 344,211 *cis* sQTLs affecting 9,051 sGenes (8,662 protein coding genes and 389 lincRNAs). Results are summarized in Table S9. As expected, the number of sGenes over the number of tested genes increases with the tissue sample size ($r^2 = 0.88$, Fig S27a). In contrast to the V7 analysis, here we observe some signs of saturation for larger sample sizes. The overlap between V8 and V7 sQTLs, evaluated considering the variant-gene-tissue trios tested in both runs, was substantial (median *Jaccard* index of sGenes across tissues 0.54, Fig. S27b). Indeed, V7 sQTLs are essentially a subset of V8 sQTLs (median minimum overlap across tissues 0.87). The increased sample sizes available in GTEx V8, with respect to GTEx V7, led to the identification of a larger number of sQTLs, with overall smaller effect sizes (Fig. S27c).

We evaluated the overlap between the V8 `sQTLseekeR2` sQTLs and the ones produced by the GTEx Consortium using intron excision ratios obtained by `LeafCutter` as splicing phenotypes and univariate linear regression, as implemented in `FastQTL`, for association testing[3]. Despite the large differences between the two analyses (regarding the set of variants and genes tested, the filters applied, the length of the the *cis* windows selected, the splicing phenotypes and covariates used, the methodology for association testing employed, etc.), we observe a moderate overlap between both sQTL sets (median *Jaccard* index of sGenes across tissues 0.33, median $\pi_1$ of `sQTLseekeR2` in `LeafCutter` sQTL-sGene associations across tissues 0.42, Fig. S27d). Note that the overlap was evaluated considering the variant-gene-tissue trios tested in both runs. Overall, the number of sQTLs identified by `LeafCutter` was larger than the number of sQTLs identified by `sQTLseekeR2`. Nevertheless, `sQTLseekeR2` added 53,420 new sQTLs and 1,136 new sGenes to those reported by the GTEx Consortium, capturing a population of events that tend to escape detection using the approach based on `LeafCutter` (see Supplementary Note 2).
