## Supplementary material for "Identification and analysis of splicing quantitative trait loci across multiple tissues in the human genome": supp.tables_figures.pdf

### Supplementary Tables and Figures

**Table S1.** Number of samples, variants and genes tested; variant-gene associations, sQTLs and sGenes identified across tissues (Benjamini-Hochberg FDR $< 0.05$) in GTEx V7.

**Table S2.** Histone mark changes at heteropleiotropic loci. For each heteropleiotropic variant, we report the associated sGene (gene $g_1$ in tissue $t_1$), eGene (gene $g_2$ in tissue $t_2$), histone mark and tissue where the variant overlaps the mark (either $t_1$ or $t_2$).

**Table S3.** Values of $\tau_s$ and $\tau$ corresponding to 469 sGenes under strong tissue-specific splicing regulation ($\tau_s > 0.9325$), 81 of which do not display tissue-specific expression ($\tau < 0.1965$). The $\tau_s$ and $\tau$ thresholds correspond, respectively, to the top 20 percentile of the distribution of $\tau_s$ values and the bottom 20 percentile of the distribution of $\tau$ values (see Methods).

**Table S4.** Top 10 diseases with the highest sQTL enrichment, with respect to non-sQTLs, on the corresponding GWAS loci. We focused on diseases with at least 30 non-sQTLs overlapping their GWAS hits. The odds-ratio (OR) for the enrichment is reported. All the enrichments are significant with FDRs $< 2.17 \cdot 10^{-16}$. In each case, we provide references of previous works relating alternative splicing to the disease pathophysiology.

**Table S5.** ENTEx ChIP-seq experiments for 6 histone marks across 29 tissues. The accession numbers provided allow to uniquely identify the samples on the ENCODE portal (`https://www.encodeproject.org/`).

**Table S6.** ENCODE eCLIP experiments for 114 target RNA-binding proteins in HepG2 and K562 cell lines. The accession numbers provided allow to uniquely identify the samples on the ENCODE portal (`https://www.encodeproject.org/`).

**Table S7.** ENCODE RNA-seq experiments on nuclear and cytoplasmic fractions of 13 human cell lines. The accession numbers provided allow to uniquely identify the samples on the ENCODE portal (`https://www.encodeproject.org/`).

**Table S8.** Number of samples, variants and genes tested; variant-gene associations, sQTLs and sGenes identified across tissues (Benjamini-Hochberg FDR $< 0.05$) in GTEx V7 using the vector of intron excision ratios of an intron cluster obtained by `LeafCutter` as splicing phenotype.

**Table S9.** Number of samples, variants and genes tested; variant-gene associations, sQTLs and sGenes identified across tissues (Benjamini-Hochberg FDR $< 0.05$) in GTEx V8. Note that *Cells - Transformed fibroblasts* from previous releases has been modified to *Cells - Cultured fibroblasts*.

#### Table S1

| Tissue | Samples | Variants | Genes | Associations | sQTLs | sGenes |
|---|---|---|---|---|---|---|
| Adipose - Subcutaneous | 385 | 1,407,668 | 10,425 | 58,152 | 57,347 | 2,206 |
| Adipose - Visceral (Omentum) | 313 | 1,263,199 | 10,454 | 44,103 | 43,410 | 1,794 |
| Adrenal Gland | 175 | 986,571 | 10,272 | 26,289 | 25,863 | 1,164 |
| Artery - Aorta | 267 | 1,205,692 | 10,365 | 42,011 | 41,378 | 1,609 |
| Artery - Coronary | 152 | 958,484 | 10,511 | 18,877 | 18,637 | 887 |
| Artery - Tibial | 388 | 1,379,506 | 10,114 | 54,533 | 53,769 | 1,994 |
| Brain - Amygdala | 88 | 719,074 | 10,075 | 5,943 | 5,884 | 314 |
| Brain - Anterior cingulate cortex (BA24) | 109 | 869,601 | 10,337 | 9,559 | 9,446 | 477 |
| Brain - Caudate (basal ganglia) | 144 | 991,073 | 10,382 | 14,479 | 14,195 | 695 |
| Brain - Cerebellar Hemisphere | 125 | 916,855 | 10,729 | 15,582 | 15,249 | 731 |
| Brain - Cerebellum | 154 | 1,041,020 | 10,807 | 25,631 | 25,202 | 1,017 |
| Brain - Cortex | 136 | 1,017,886 | 10,560 | 16,007 | 15,799 | 778 |
| Brain - Frontal Cortex (BA9) | 118 | 957,206 | 10,633 | 12,447 | 12,316 | 613 |
| Brain - Hippocampus | 111 | 853,881 | 10,300 | 8,228 | 8,148 | 422 |
| Brain - Hypothalamus | 108 | 885,872 | 10,589 | 10,013 | 9,914 | 490 |
| Brain - Nucleus accumbens (basal ganglia) | 130 | 957,657 | 10,493 | 12,775 | 12,661 | 604 |
| Brain - Putamen (basal ganglia) | 111 | 801,461 | 9,872 | 11,656 | 11,478 | 502 |
| Brain - Spinal cord (cervical c-1) | 83 | 669,619 | 10,327 | 6,942 | 6,873 | 334 |
| Brain - Substantia nigra | 80 | 656,998 | 10,141 | 5,037 | 4,970 | 280 |
| Breast - Mammary Tissue | 251 | 1,175,934 | 10,475 | 39,953 | 39,299 | 1,558 |
| Cells - EBV-transformed lymphocytes | 117 | 590,120 | 8,421 | 16,282 | 16,102 | 731 |
| Cells - Transformed fibroblasts | 300 | 1,127,243 | 9,464 | 43,890 | 43,293 | 1,652 |
| Colon - Sigmoid | 203 | 1,102,171 | 10,577 | 32,983 | 32,563 | 1,321 |
| Colon - Transverse | 246 | 1,115,378 | 10,321 | 34,833 | 34,415 | 1,359 |
| Esophagus - Gastroesophageal Junction | 213 | 1,101,011 | 10,480 | 32,313 | 31,778 | 1,288 |
| Esophagus - Mucosa | 358 | 1,298,073 | 10,568 | 48,733 | 47,941 | 1,901 |
| Esophagus - Muscularis | 335 | 1,358,213 | 10,477 | 50,653 | 49,826 | 1,920 |
| Heart - Atrial Appendage | 264 | 1,137,415 | 9,958 | 33,263 | 32,723 | 1,284 |
| Heart - Left Ventricle | 272 | 984,193 | 8,991 | 24,583 | 24,122 | 991 |
| Liver | 153 | 738,737 | 9,282 | 12,635 | 12,471 | 630 |
| Lung | 383 | 1,497,417 | 11,309 | 56,078 | 55,251 | 2,159 |
| Minor Salivary Gland | 85 | 625,502 | 10,330 | 6,782 | 6,655 | 442 |
| Muscle - Skeletal | 491 | 1,182,336 | 8,835 | 49,633 | 48,982 | 1,810 |
| Nerve - Tibial | 361 | 1,501,244 | 11,075 | 58,707 | 57,990 | 2,224 |
| Ovary | 122 | 854,281 | 10,433 | 16,883 | 16,714 | 748 |
| Pancreas | 220 | 941,010 | 9,505 | 24,737 | 24,545 | 1,044 |
| Pituitary | 157 | 1,093,565 | 11,282 | 24,929 | 24,647 | 1,083 |
| Prostate | 132 | 936,634 | 11,030 | 15,593 | 15,393 | 809 |
| Skin - Not Sun Exposed (Suprapubic) | 335 | 1,303,847 | 10,730 | 49,220 | 48,263 | 1,899 |
| Skin - Sun Exposed (Lower leg) | 414 | 1,440,899 | 10,836 | 60,061 | 58,953 | 2,232 |
| Small Intestine - Terminal Ileum | 122 | 811,821 | 10,648 | 15,218 | 15,089 | 726 |
| Spleen | 146 | 894,283 | 10,532 | 22,998 | 22,781 | 1,043 |
| Stomach | 237 | 1,044,155 | 10,020 | 27,184 | 26,905 | 1,184 |
| Testis | 225 | 1,513,533 | 12,687 | 58,402 | 57,589 | 2,111 |
| Thyroid | 399 | 1,523,381 | 11,133 | 64,120 | 62,919 | 2,434 |
| Uterus | 101 | 781,865 | 10,720 | 12,755 | 12,685 | 635 |
| Vagina | 106 | 790,813 | 10,768 | 12,140 | 11,923 | 578 |
| Whole Blood | 369 | 633,825 | 6,327 | 21,566 | 21,318 | 920 |
| **Average** | 214 | 1,034,130 | 10,304 | 28,446 | 28,034 | 1,158 |
| **Total** | 10,294 | 3,588,609 | 16,010 | 216,961 | 210,485 | 6,963 |

#### Table S8

| Tissue | Samples | Variants | Genes | Associations | sQTLs | sGenes |
|---|---|---|---|---|---|---|
| Adipose - Subcutaneous | 385 | 758,333 | 5,975 | 44,148 | 41,083 | 1,625 |
| Adipose - Visceral (Omentum) | 313 | 645,188 | 5,801 | 32,899 | 30,526 | 1,340 |
| Adrenal Gland | 175 | 357,993 | 4,302 | 19,987 | 18,129 | 842 |
| Artery - Aorta | 267 | 580,788 | 5,284 | 31,441 | 29,361 | 1,209 |
| Artery - Coronary | 152 | 431,275 | 5,266 | 18,085 | 17,140 | 824 |
| Artery - Tibial | 388 | 711,276 | 5,499 | 41,560 | 38,451 | 1,472 |
| Brain - Amygdala | 88 | 182,912 | 3,261 | 5,876 | 5,371 | 298 |
| Brain - Anterior cingulate cortex (BA24) | 109 | 243,497 | 3,642 | 9,462 | 8,682 | 437 |
| Brain - Caudate (basal ganglia) | 144 | 304,661 | 3,919 | 12,104 | 11,104 | 571 |
| Brain - Cerebellar Hemisphere | 125 | 405,495 | 5,232 | 23,494 | 19,834 | 952 |
| Brain - Cerebellum | 154 | 461,286 | 5,327 | 30,505 | 26,398 | 1,141 |
| Brain - Cortex | 136 | 364,844 | 4,398 | 15,213 | 13,878 | 667 |
| Brain - Frontal Cortex (BA9) | 118 | 311,326 | 4,025 | 11,854 | 10,918 | 558 |
| Brain - Hippocampus | 111 | 218,704 | 3,402 | 7,940 | 7,250 | 385 |
| Brain - Hypothalamus | 108 | 254,314 | 3,938 | 9,055 | 8,315 | 456 |
| Brain - Nucleus accumbens (basal ganglia) | 130 | 273,022 | 3,794 | 11,931 | 10,988 | 571 |
| Brain - Putamen (basal ganglia) | 111 | 214,012 | 3,318 | 8,450 | 7,645 | 436 |
| Brain - Spinal cord (cervical c-1) | 83 | 206,183 | 3,731 | 6,777 | 6,415 | 363 |
| Brain - Substantia nigra | 80 | 166,463 | 3,286 | 4,716 | 4,356 | 257 |
| Breast - Mammary Tissue | 251 | 610,844 | 5,895 | 35,075 | 32,411 | 1,371 |
| Cells - EBV-transformed lymphocytes | 117 | 315,504 | 4,442 | 22,263 | 21,048 | 933 |
| Cells - Transformed fibroblasts | 300 | 589,043 | 4,990 | 33,247 | 30,955 | 1,218 |
| Colon - Sigmoid | 203 | 462,574 | 5,134 | 27,428 | 25,914 | 1,108 |
| Colon - Transverse | 246 | 456,973 | 4,909 | 21,475 | 19,961 | 969 |
| Esophagus - Gastroesophageal Junction | 213 | 441,111 | 4,824 | 25,416 | 23,862 | 1,057 |
| Esophagus - Mucosa | 358 | 477,846 | 4,524 | 26,603 | 25,044 | 1,115 |
| Esophagus - Muscularis | 335 | 565,549 | 4,914 | 36,681 | 33,985 | 1,353 |
| Heart - Atrial Appendage | 264 | 478,621 | 4,480 | 22,583 | 21,026 | 928 |
| Heart - Left Ventricle | 272 | 270,243 | 2,900 | 14,371 | 13,401 | 600 |
| Liver | 153 | 190,231 | 2,963 | 11,037 | 10,085 | 523 |
| Lung | 383 | 715,284 | 6,003 | 36,182 | 34,324 | 1,527 |
| Minor Salivary Gland | 85 | 206,445 | 4,097 | 7,132 | 6,801 | 416 |
| Muscle - Skeletal | 491 | 516,129 | 4,100 | 27,539 | 26,127 | 1,089 |
| Nerve - Tibial | 361 | 753,162 | 6,180 | 48,704 | 45,256 | 1,790 |
| Ovary | 122 | 355,980 | 5,204 | 17,135 | 16,103 | 778 |
| Pancreas | 220 | 281,415 | 3,336 | 15,084 | 13,873 | 703 |
| Pituitary | 157 | 393,827 | 5,170 | 26,231 | 24,316 | 1,062 |
| Prostate | 132 | 338,794 | 5,044 | 16,227 | 15,103 | 819 |
| Skin - Not Sun Exposed (Suprapubic) | 335 | 617,127 | 5,628 | 36,497 | 34,260 | 1,485 |
| Skin - Sun Exposed (Lower leg) | 414 | 685,673 | 5,732 | 40,761 | 38,375 | 1,632 |
| Small Intestine - Terminal Ileum | 122 | 311,399 | 4,876 | 12,592 | 11,946 | 642 |
| Spleen | 146 | 328,298 | 4,525 | 19,206 | 18,026 | 895 |
| Stomach | 237 | 369,489 | 4,150 | 15,420 | 14,494 | 777 |
| Testis | 225 | 789,414 | 7,769 | 89,009 | 83,724 | 2,739 |
| Thyroid | 399 | 735,487 | 6,029 | 48,250 | 44,783 | 1,807 |
| Uterus | 101 | 334,515 | 5,323 | 14,358 | 13,405 | 711 |
| Vagina | 106 | 286,538 | 4,763 | 12,357 | 11,497 | 624 |
| Whole Blood | 369 | 295,509 | 3,020 | 18,687 | 17,085 | 732 |
| **Average** | 214 | 422,179 | 4,673 | 23,397 | 21,730 | 954 |
| **Total** | 10,294 | 2,507,537 | 14,045 | 286,349 | 263,827 | 7,471 |

**Table S9**

| Tissue | Samples | Variants | Genes | Associations | sQTLs | sGenes |
|---|---|---|---|---|---|---|
| Adipose - Subcutaneous | 581 | 1,849,202 | 10,494 | 101,905 | 100,331 | 2,990 |
| Adipose - Visceral (Omentum) | 469 | 1,759,158 | 10,604 | 79,846 | 78,490 | 2,487 |
| Adrenal Gland | 233 | 1,248,945 | 10,393 | 43,530 | 42,912 | 1,555 |
| Artery - Aorta | 387 | 1,634,096 | 10,520 | 72,071 | 70,843 | 2,272 |
| Artery - Coronary | 213 | 1,269,445 | 10,646 | 36,838 | 36,244 | 1,367 |
| Artery - Tibial | 584 | 1,808,194 | 10,208 | 92,154 | 90,652 | 2,720 |
| Brain - Amygdala | 129 | 997,280 | 10,155 | 14,306 | 14,141 | 580 |
| Brain - Anterior cingulate cortex (BA24) | 147 | 1,135,042 | 10,448 | 17,973 | 17,782 | 754 |
| Brain - Caudate (basal ganglia) | 194 | 1,269,547 | 10,425 | 28,549 | 28,010 | 1,052 |
| Brain - Cerebellar Hemisphere | 175 | 1,205,236 | 10,814 | 33,739 | 33,013 | 1,236 |
| Brain - Cerebellum | 209 | 1,355,112 | 10,902 | 44,581 | 43,584 | 1,581 |
| Brain - Cortex | 205 | 1,414,047 | 10,693 | 36,624 | 36,217 | 1,321 |
| Brain - Frontal Cortex (BA9) | 175 | 1,303,066 | 10,749 | 25,322 | 25,105 | 996 |
| Brain - Hippocampus | 165 | 1,139,777 | 10,250 | 20,053 | 19,871 | 756 |
| Brain - Hypothalamus | 170 | 1,233,848 | 10,594 | 24,432 | 24,157 | 883 |
| Brain - Nucleus accumbens (basal ganglia) | 202 | 1,303,678 | 10,457 | 28,187 | 27,832 | 1,042 |
| Brain - Putamen (basal ganglia) | 170 | 1,096,223 | 9,910 | 22,676 | 22,358 | 891 |
| Brain - Spinal cord (cervical c-1) | 126 | 971,511 | 10,456 | 17,001 | 16,865 | 669 |
| Brain - Substantia nigra | 114 | 890,437 | 10,080 | 11,457 | 11,340 | 515 |
| Breast - Mammary Tissue | 396 | 1,651,603 | 10,583 | 77,202 | 76,009 | 2,381 |
| Cells - EBV-transformed lymphocytes | 483 | 1,521,528 | 9,381 | 79,240 | 77,956 | 2,339 |
| Cells - Transformed fibroblasts | 147 | 743,422 | 8,435 | 28,101 | 27,708 | 1,045 |
| Colon - Sigmoid | 318 | 1,564,926 | 10,720 | 62,501 | 61,303 | 1,969 |
| Colon - Transverse | 368 | 1,555,065 | 10,466 | 65,543 | 64,518 | 2,082 |
| Esophagus - Gastroesophageal Junction | 330 | 1,566,911 | 10,625 | 63,217 | 62,335 | 2,011 |
| Esophagus - Mucosa | 497 | 1,700,995 | 10,651 | 78,535 | 77,340 | 2,499 |
| Esophagus - Muscularis | 465 | 1,811,046 | 10,620 | 83,766 | 82,229 | 2,537 |
| Heart - Atrial Appendage | 372 | 1,523,837 | 10,069 | 58,573 | 57,816 | 1,896 |
| Heart - Left Ventricle | 386 | 1,284,915 | 9,072 | 44,520 | 43,827 | 1,436 |
| Kidney - Cortex | 73 | 540,123 | 9,741 | 6,701 | 6,622 | 336 |
| Liver | 208 | 948,583 | 9,366 | 26,424 | 26,090 | 1,003 |
| Lung | 515 | 1,937,020 | 11,391 | 92,364 | 90,862 | 2,803 |
| Minor Salivary Gland | 144 | 1,017,962 | 10,674 | 22,910 | 22,486 | 966 |
| Muscle - Skeletal | 706 | 1,482,994 | 8,910 | 84,962 | 83,542 | 2,440 |
| Nerve - Tibial | 532 | 2,007,715 | 11,196 | 104,184 | 102,300 | 3,075 |
| Ovary | 167 | 1,101,628 | 10,571 | 32,306 | 31,812 | 1,176 |
| Pancreas | 305 | 1,213,421 | 9,490 | 44,223 | 43,518 | 1,549 |
| Pituitary | 237 | 1,490,248 | 11,386 | 54,580 | 53,717 | 1,795 |
| Prostate | 221 | 1,385,232 | 11,198 | 42,801 | 42,063 | 1,556 |
| Skin - Not Sun Exposed (Suprapubic) | 517 | 1,785,930 | 10,836 | 88,168 | 86,855 | 2,710 |
| Skin - Sun Exposed (Lower leg) | 605 | 1,864,176 | 10,901 | 99,943 | 98,230 | 3,032 |
| Small Intestine - Terminal Ileum | 174 | 1,118,322 | 10,838 | 29,823 | 29,563 | 1,179 |
| Spleen | 227 | 1,266,159 | 10,739 | 48,529 | 47,709 | 1,745 |
| Stomach | 324 | 1,387,919 | 10,133 | 45,025 | 44,442 | 1,552 |
| Testis | 322 | 2,033,706 | 12,712 | 111,059 | 109,218 | 3,087 |
| Thyroid | 574 | 1,992,634 | 11,235 | 107,543 | 105,641 | 3,174 |
| Uterus | 129 | 982,504 | 10,823 | 22,514 | 22,209 | 908 |
| Vagina | 141 | 1,011,962 | 10,940 | 22,196 | 21,866 | 870 |
| Whole Blood | 670 | 799,480 | 6,132 | 44,875 | 44,132 | 1,476 |
| **Average** | 310 | 1,370,935 | 10,380 | 51,501 | 50,687 | 1,679 |
| **Total** | 15,201 | 4,074,385 | 16,202 | 356,682 | 344,211 | 9,051 |

**Fig. S1.** Median sQTL effect size (absolute maximum difference in adjusted transcript relative expression between genotype groups, MD), computed for each tissue (y-axis), with respect to the tissue sample size (x-axis).

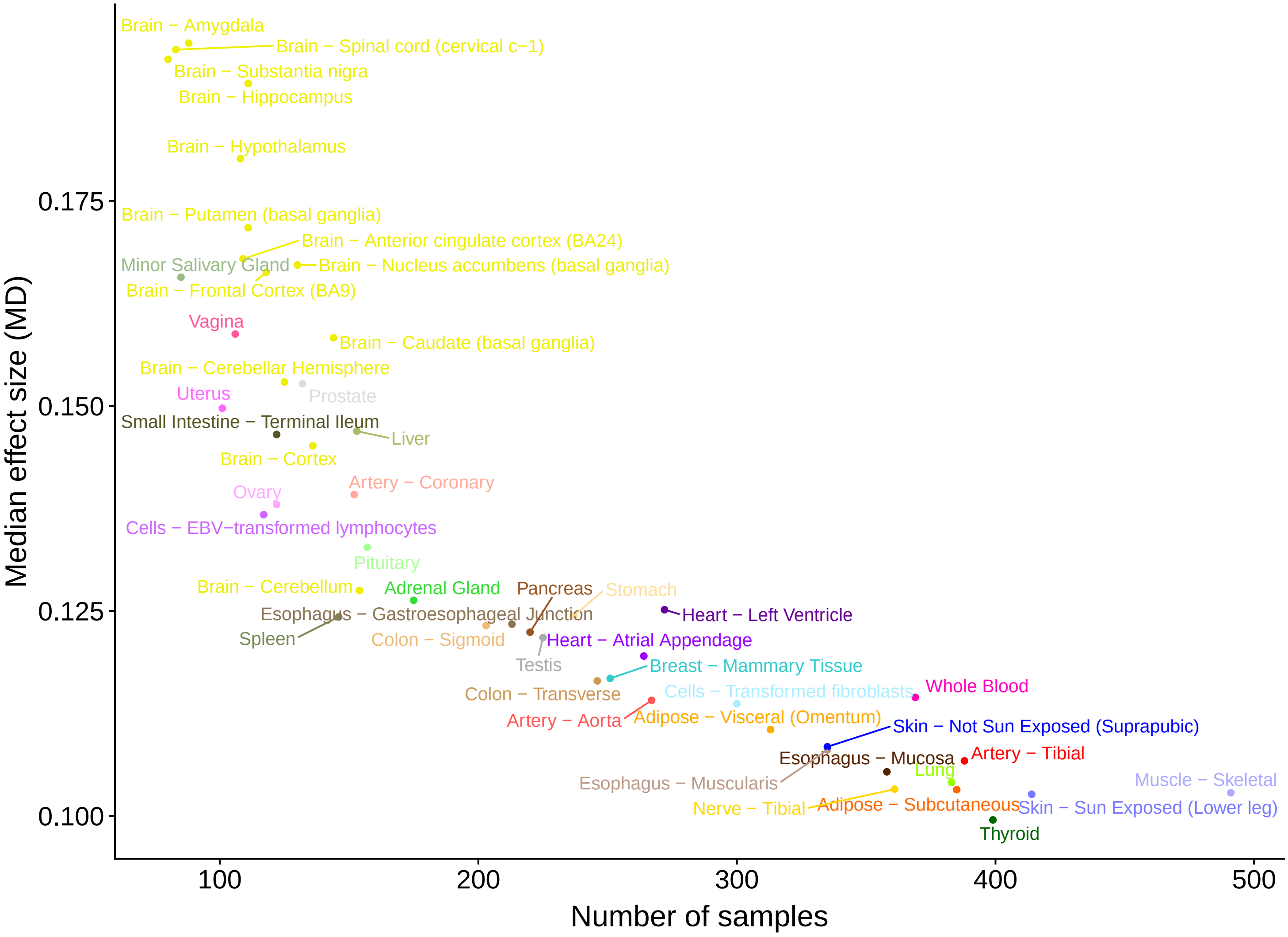

**Fig. S2. a)** Multidimensional scaling-based representation of the semantic dissimilarities between non-redundant Gene Ontology (Biological Process) terms enriched among sGenes. Each term is represented by a circle, being its size the number of tissues in which the term is enriched and its color the minimum $-log_{10}$ *p* value (hypergeometric test) for the term enrichment across tissues. GO terms that lie close to each other are semantically more similar. **b)** Analogous representation for genes without sQTLs in any tissue.

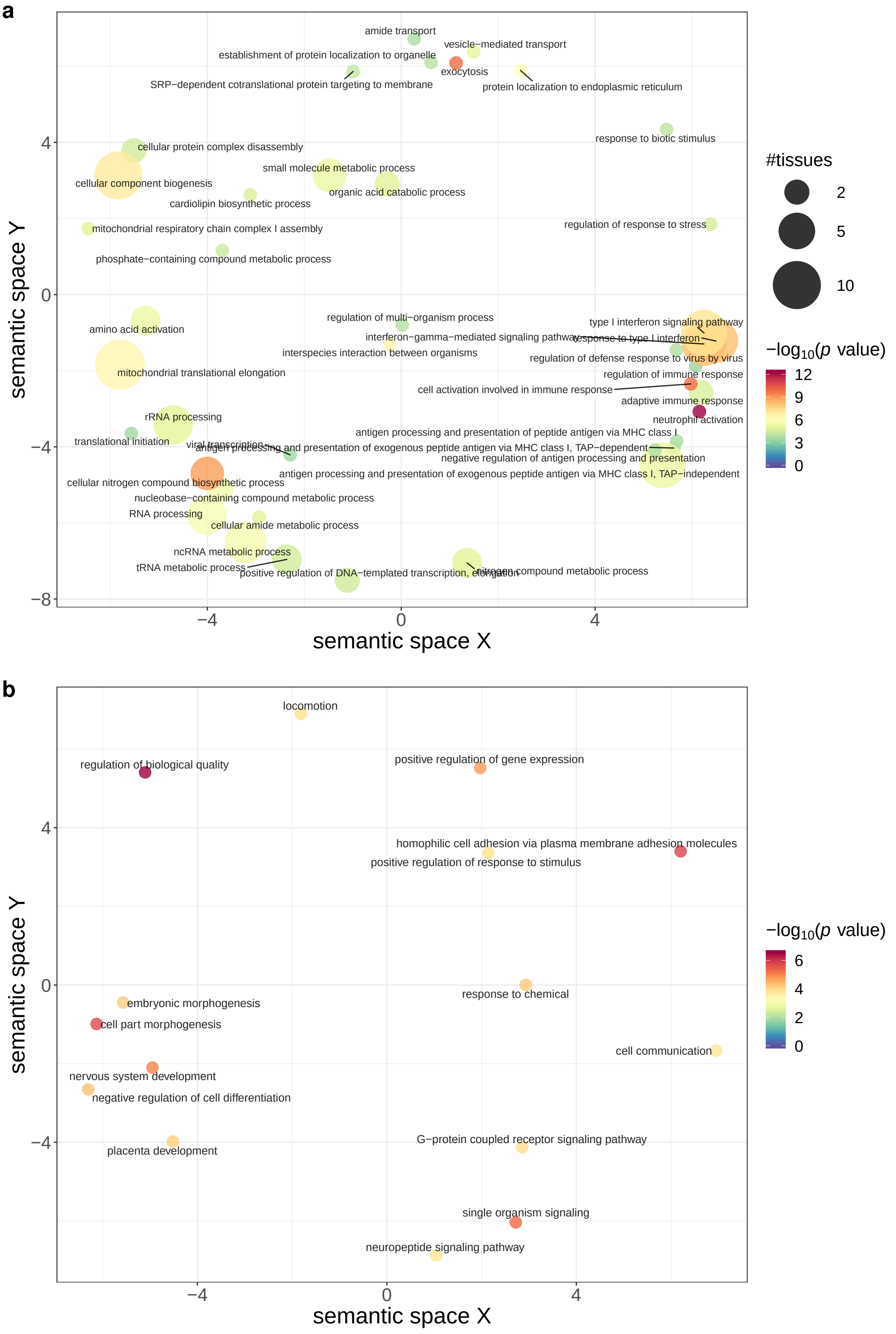

**Fig. S3**. Histogram of the replication *p* values of GTEx whole blood sQTLs in three immune cell types (CD14$^+$ monocytes, CD16$^+$ neutrophils, and naive CD4$^+$ T cells) from the Blueprint Project (BP). $\pi_1$ statistics are also shown.

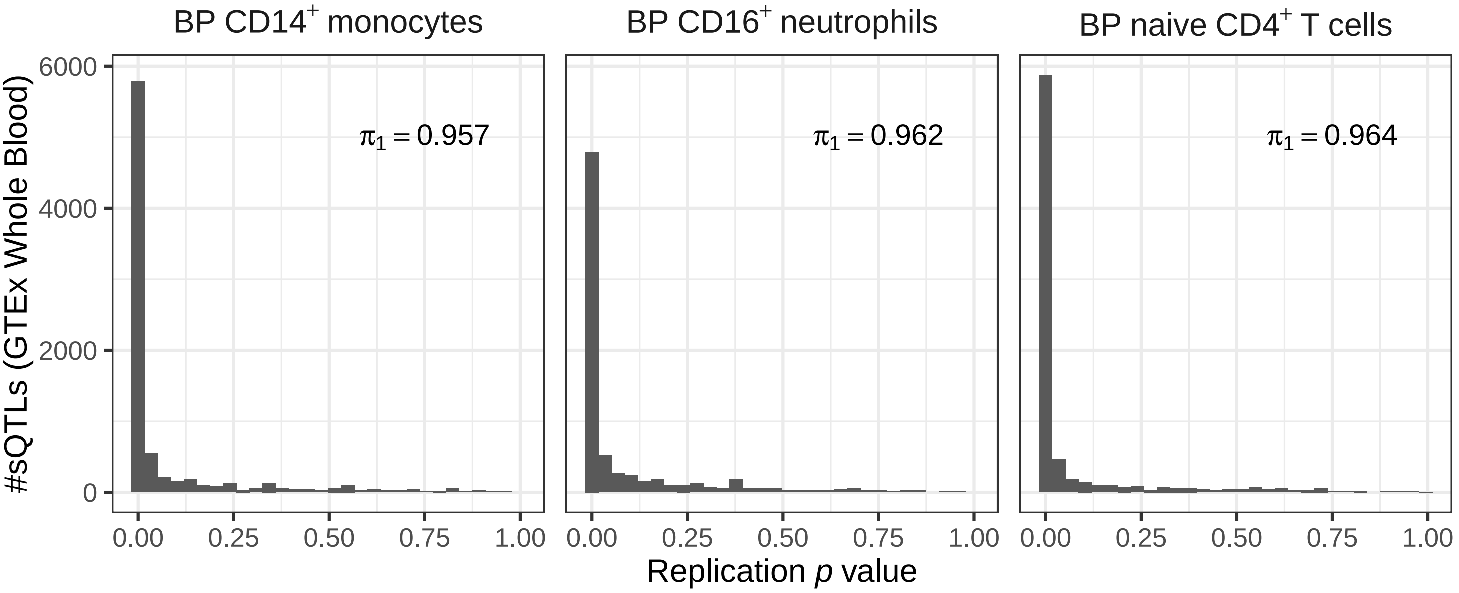

**Fig. S4**. **a)** Types of alternative splicing (AS) events associated to sQTLs. The height of each bar corresponds to the average proportion of AS events of a given type associated to sQTLs across GTEx tissues (see Methods). Error bars represent standard deviation. **b)** Representation in two dimensions, for each tissue, of the vector of proportions of the different types of AS events, obtained by t-SNE. Tissues with a similar pattern of AS events associated to sQTLs are closer in the bi-dimensional space, with brain subregions forming a distinct cluster. **c)** Comparison of the distribution of the proportions of exon skipping and complex 3' events associated to sQTLs in brain (yellow) and non-brain tissues (white). Brain subregions display a larger proportion of simple events affecting internal exons and introns, such as exon inclusion (Wilcoxon Rank-Sum test $p$ value $6.94 \cdot 10^{-10}$), and a smaller proportion of events affecting first/last exons and UTRs, such as complex 3' events (Wilcoxon Rank-Sum test $p$ value $3.30 \cdot 10^{-6}$).

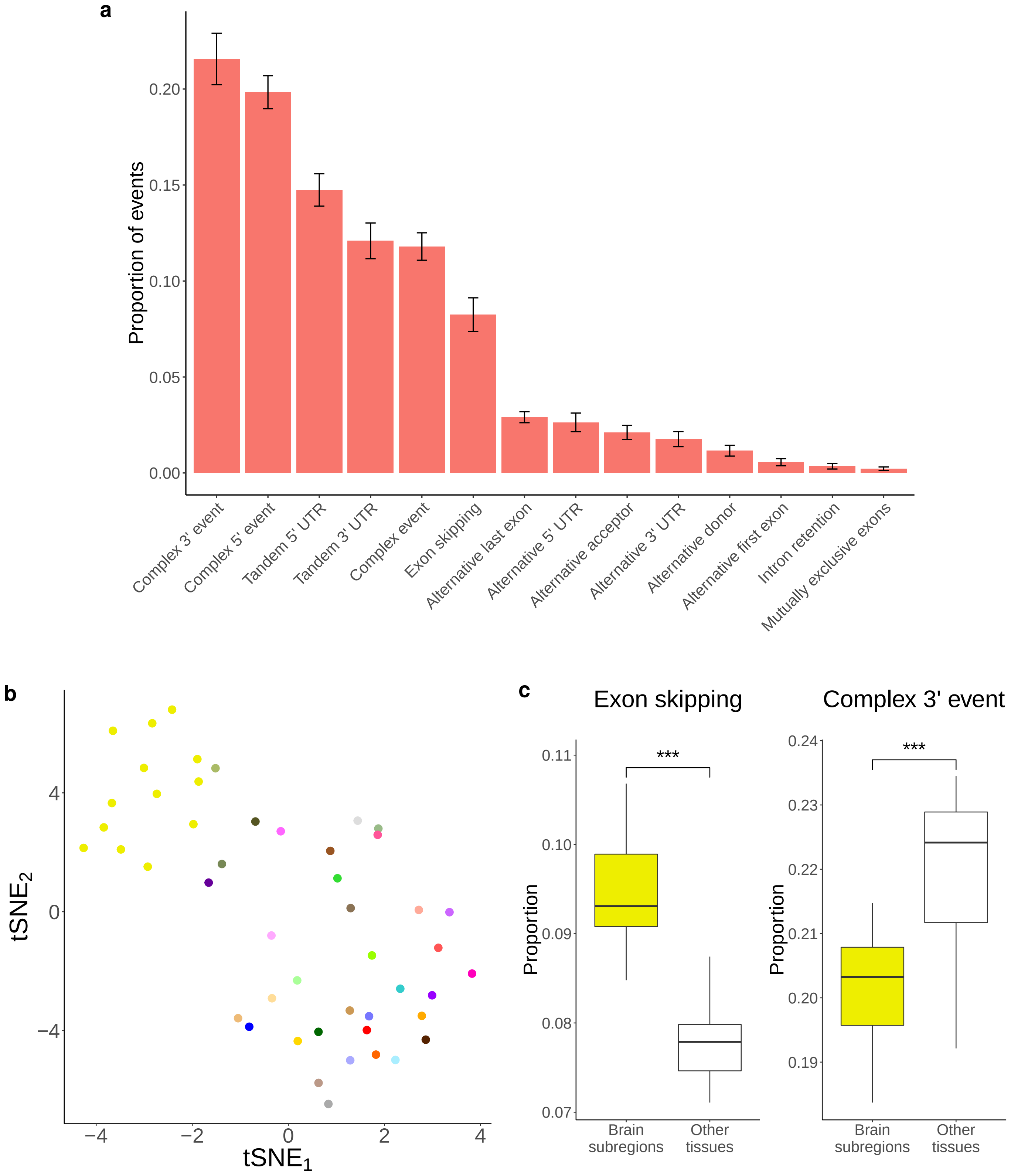

**Fig. S5.** **a)** A variant $v$ is considered heteropleiotropic in two different tissues (i.e. $t_1$ and $t_2$) if: i) it is an sQTL, but not an eQTL, for gene $g_1$ in tissue $t_1$, ii) it is an eQTL, but not an sQTL, for gene $g_2$ in tissue $t_2$, iii) it is neither an sQTL nor an eQTL for gene $g_2$ in tissue $t_1$, iv) it is neither an sQTL nor an eQTL for gene $g_1$ in tissue $t_2$. **d)** Example of an heteropleiotropic locus. The SNP rs11603538 (chr11:63,986,713, C/A), an sQTL for the gene FERMT3 (chr11:63,974,150-63,991,354, forward strand) in Spleen, but not in Muscle Skeletal. The SNP is not an eQTL for FERMT3 in any of the two tissues. In contrast, the SNP is an eQTL for the gene TRPT1 (chr11:63,991,272-63,993,726, reverse strand) in Muscle Skeletal, but not in Spleen. The SNP is not an sQTL for TRPT1 in any of the two tissues. In the left panel, the dots represent the $-log_{10}$ *p* values of association with the expression (green) and splicing (red) of the two genes in the two tissues, for variants in a 20Kb window centered at rs11603538 (the $-log_{10}$ *p* values corresponding to rs11603538 are highlighted by a diamond). The transparency of the dots depends on the $-log_{10}$ *p* value. The significance level for each molecular trait, gene and tissue is shown as a colored, dashed horizontal line. When this line is not present, the gene-level *p* value is above the 0.05 FDR threshold and hence no variant is significantly associated with this molecular trait in this tissue (see Methods). The shaded area represents the position of a H3K36me3 ChIP-seq peak (see below). The right panel shows the fold-change signal of the H3K36me3 histone mark with respect to the input across 3 ENTEx donors in Spleen and Muscle Skeletal, in the same genomic region of the left panel. The solid line and coloured area correspond to the mean signal and its standard error across 3 ENTEx donors, respectively. The location of the SNP and the overlapping ChIP-seq peak (intersection of the peaks in the 3 donors) are also displayed.

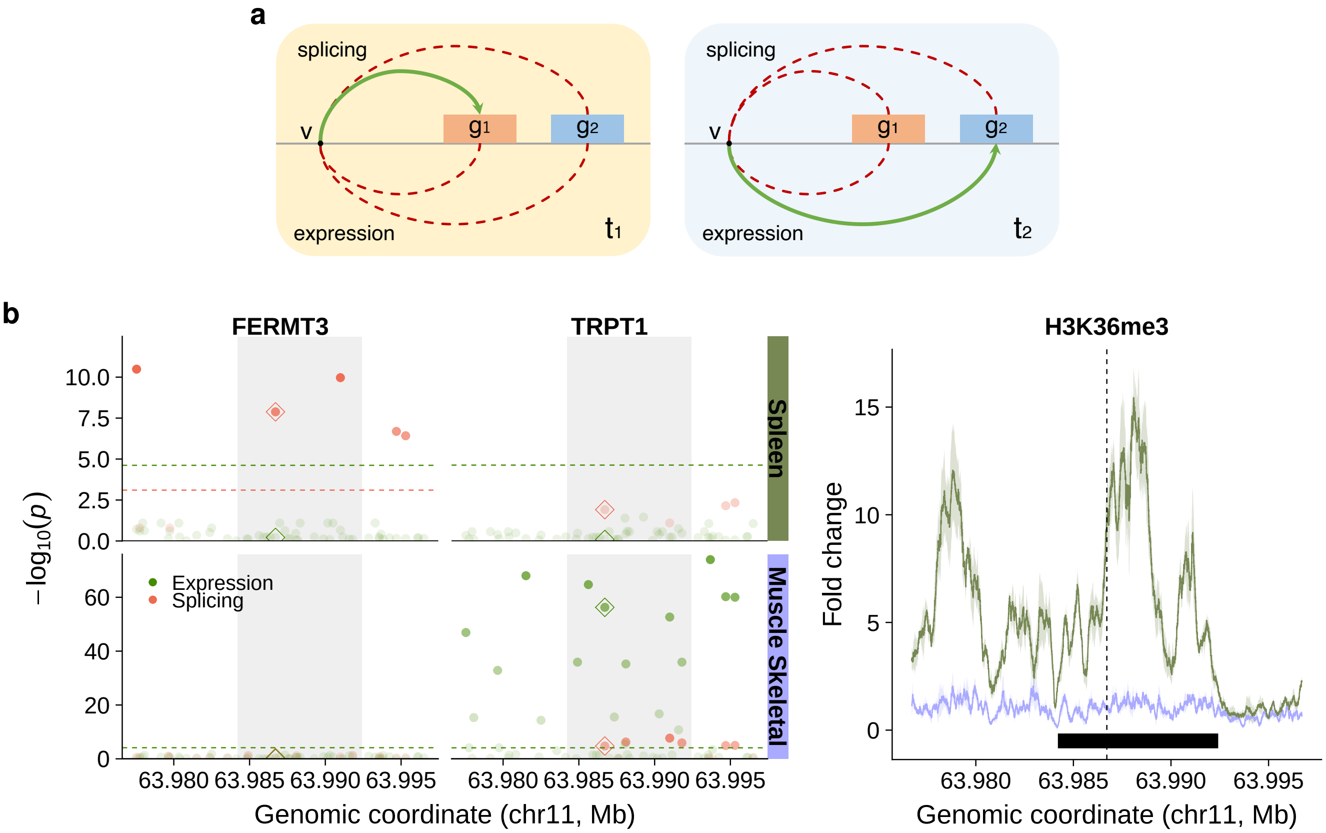

**Fig. S6**. Heatmap showing sQTL sharing patterns across GTEx tissues. *Jaccard* index computed on exact variant-gene pairs is employed as sharing estimate. Tissue specificity is estimated as $1 - \bar{j}_t$, where $\bar{j}_t$ is the mean *Jaccard* index between a given tissue $t$ and the others. Hierarchical clustering of the tissues based on the sharing patterns is also displayed, together with the corresponding tissue colors, sample sizes and tissue specificity estimates.

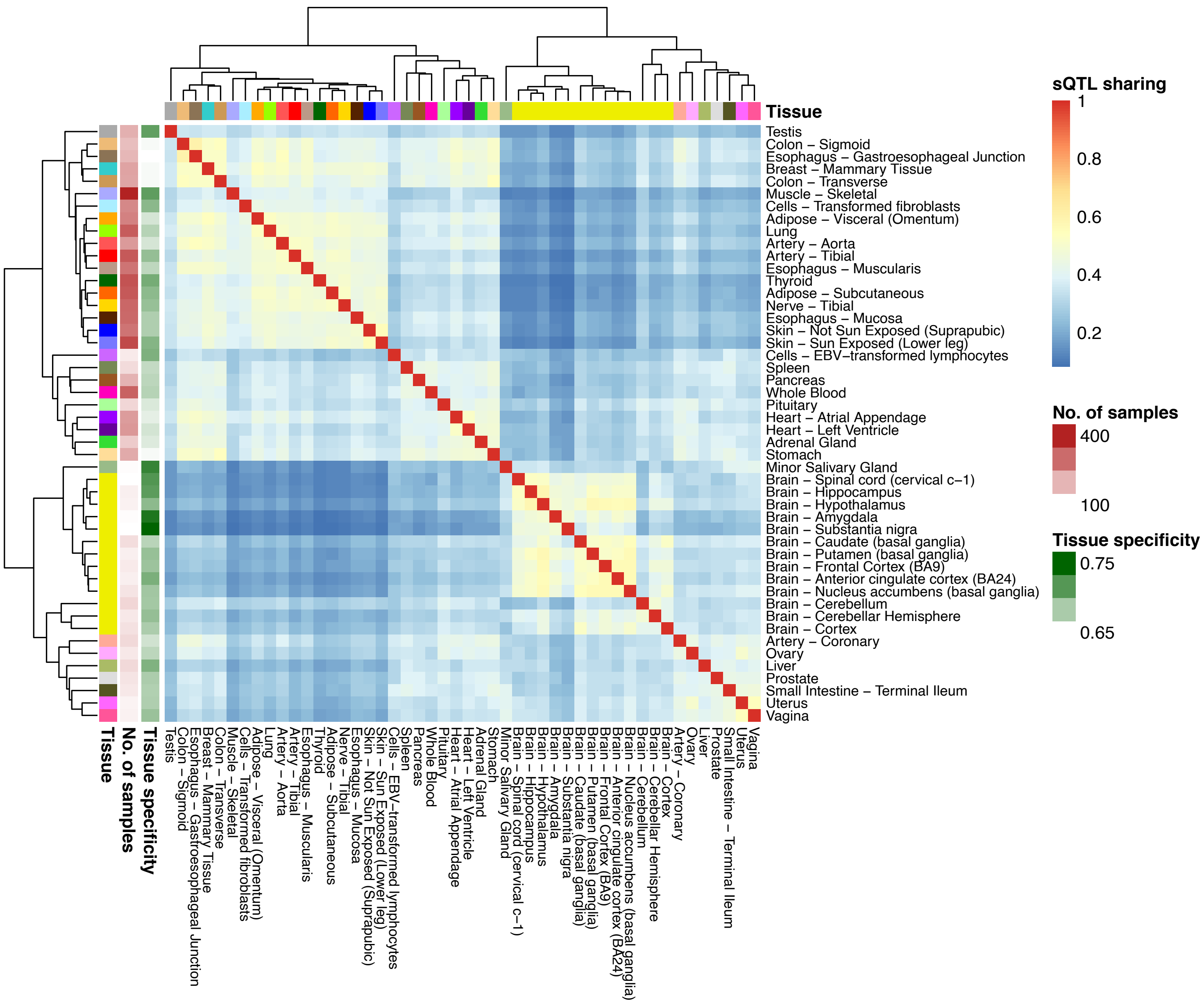

**Fig. S7.** For all variant-gene pairs tested in all GTEx tissues with $n \geq 70$, we show the distribution of the number of tissues in which the variants are identified as sQTLs for the target genes separately **a)** for high (MD in [0.2, 1]) and low (MD in [0.05, 0.1]) effect size sQTLs and **b)** for each quantile of the sample size distribution (note that the same variant-gene pair can be included in more than one quantile group). In each case, the median number of tissues is displayed as a black horizontal line.

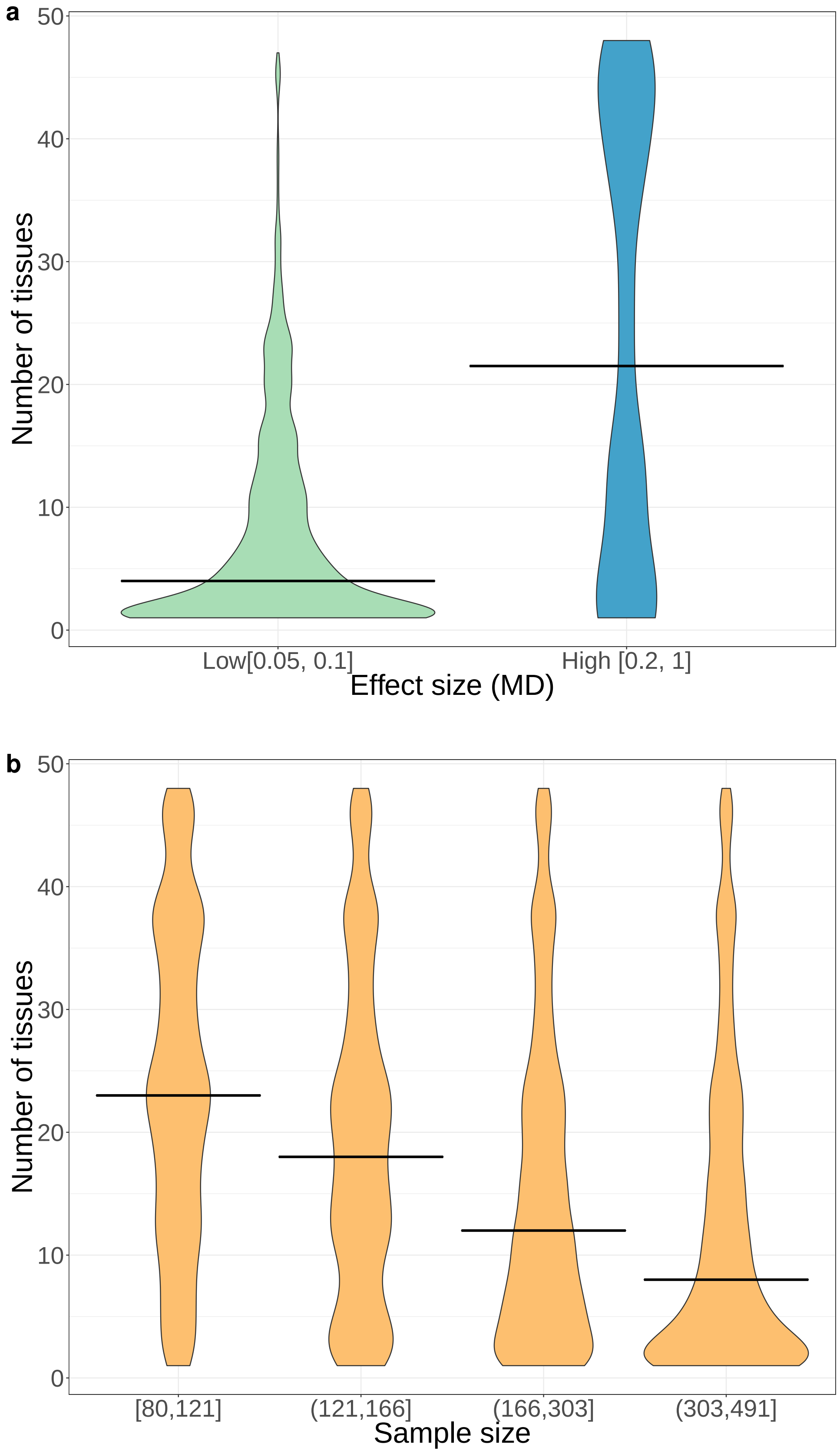

**Fig. S8**. Heatmap of sQTL sharing for downsampled GTEx datasets. GTEx tissues were randomly downsampled once to **a)** 100 **b)** 200 and **c)** 300 samples, respectively (see Methods). Sharing estimates range from 0 (low sharing, blue) to 1 (high sharing, red). In addition, we display the hierarchical clustering of the tissues based on sQTL sharing, together with the tissue specificity estimates.

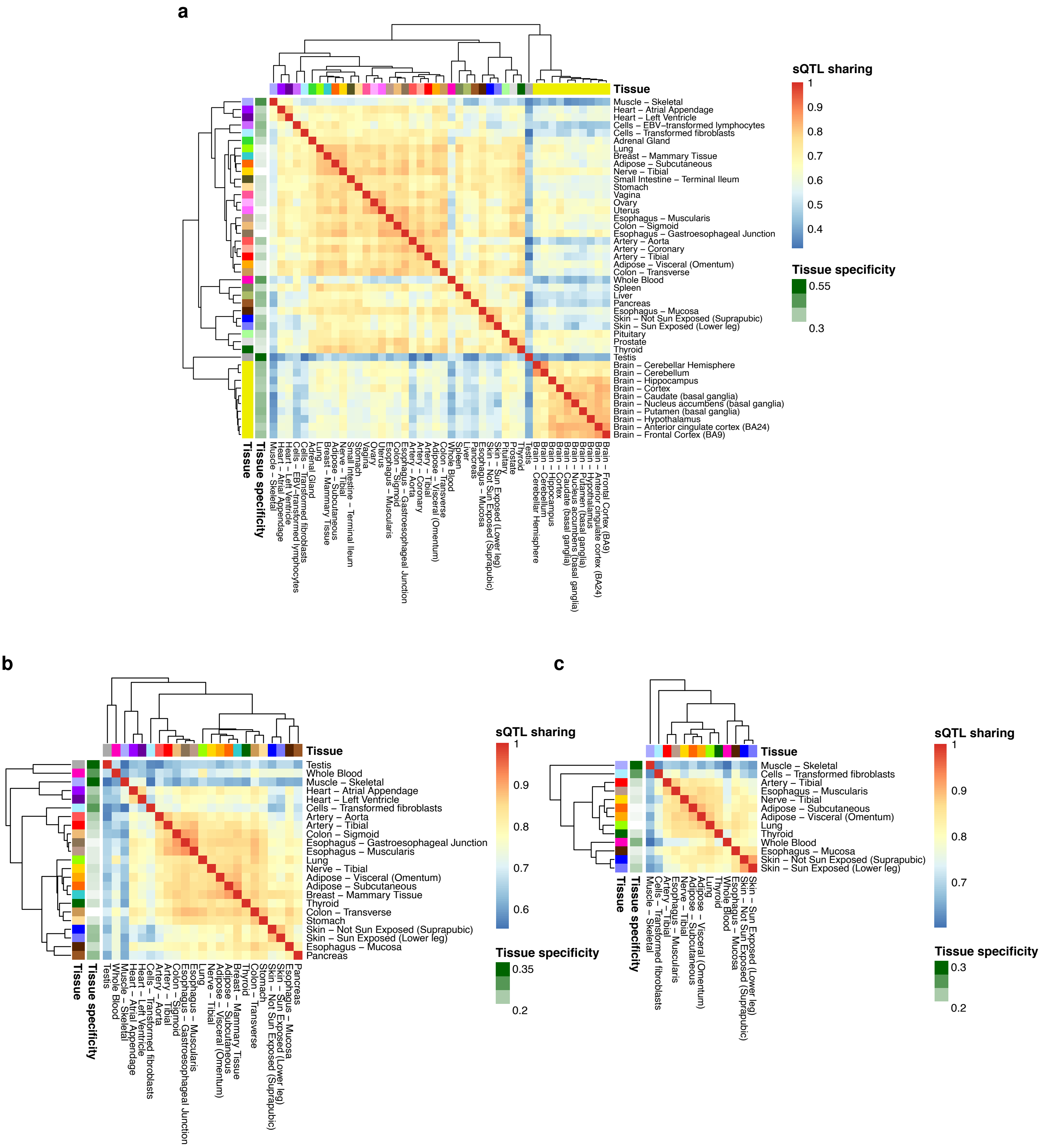

**Fig. S9**. **a)** Illustrative example of the centroid distance approach for sQTL sharing (see Methods). A SNP with alleles A>G is an sQTL for a gene with 3 isoforms in a given tissue $t_1$ (red). The average individual (centroid) of each genotype group can be represented as a point in a three-dimensional space, whose coordinates are the mean relative abundances of each isoform across individuals in this genotype group. As relative abundances add up to one, these points are located on a two-dimensional simplex (i.e. a triangle). The left panel shows a SNP that is a sQTL for the same gene in tissue $t_2$ (green), affecting isoform abundances in the same way as in tissue $t_1$ (red). Note that here we consider a single variant-gene pair ($p = 1$), and $d$ is given by the sum of the lengths of the dashed lines. Hence, in this case $d(t_1, t_2)$ would be small. In the middle panel, the SNP is not a sQTL for the same gene in tissue $t_3$ (blue), and therefore $d(t_1, t_3)$ would be large. In the right panel, the SNP is a sQTL for the same gene in tissue $t_4$ (yellow), but the associated change in splicing isoform abundances is different, and $d(t_1, t_4)$ would be large. **b)** Hierarchical clustering built using the centroid distance approach. **c)** Estimates of similarity (Baker's Gamma) between tissue dendrograms obtained using MD correlations, the *Jaccard* index and the centroid distance approach. The dendrogram obtained with `mashR` on `LeafCutter` sQTLs from GTEx V8 is also compared. All similarity estimates are significantly different from 0 (permutation test *p* value $< 10^{-4}$).

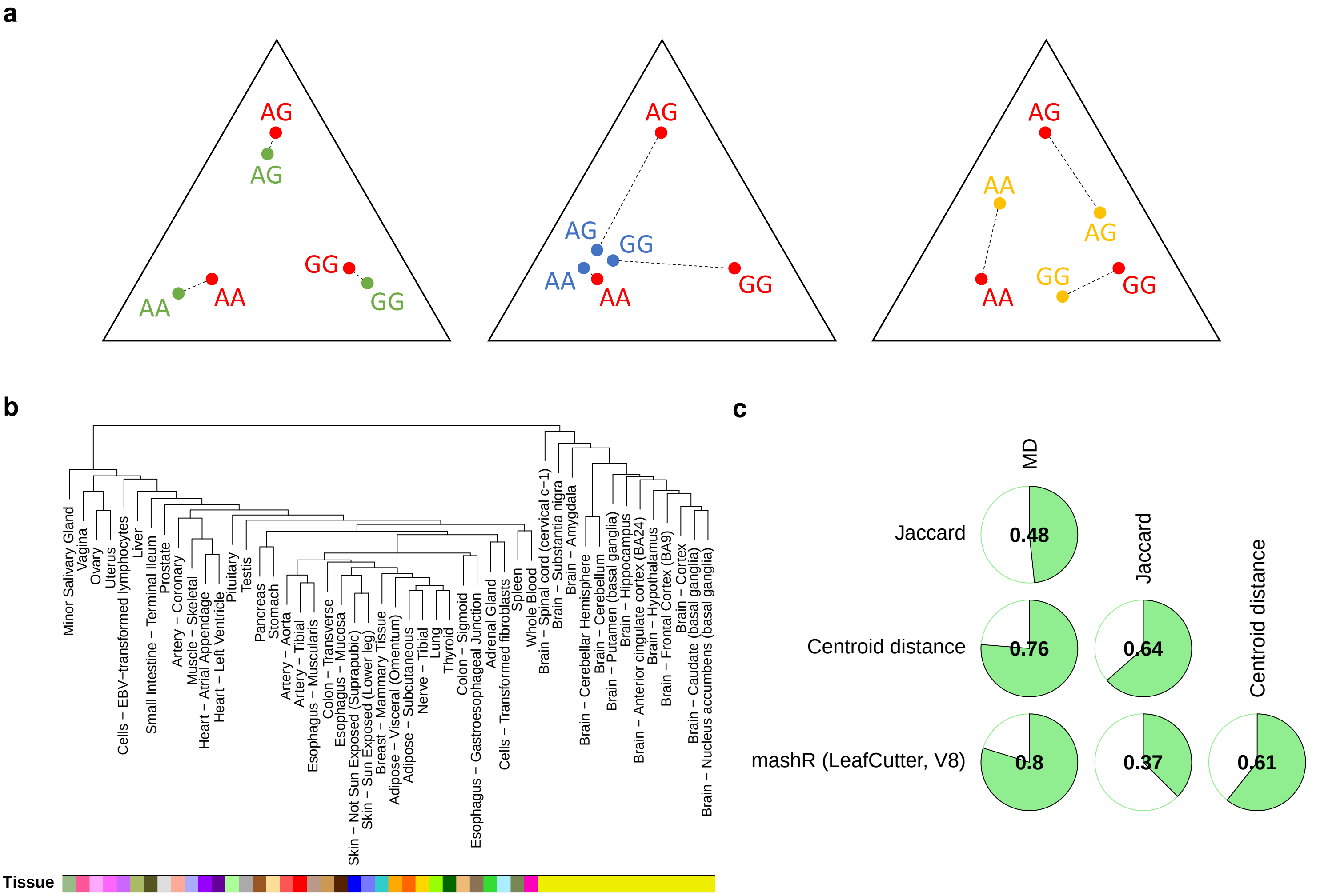

**Fig. S10**. Density of functional annotations (mean number of annotations per Kb, y-axis) with respect to the distance to sQTLs (x-axis). The location of the different functional elements was obtained from the Ensembl Regulation dataset (see Methods). The distance 0 corresponds to the functional annotations that overlap sQTLs.

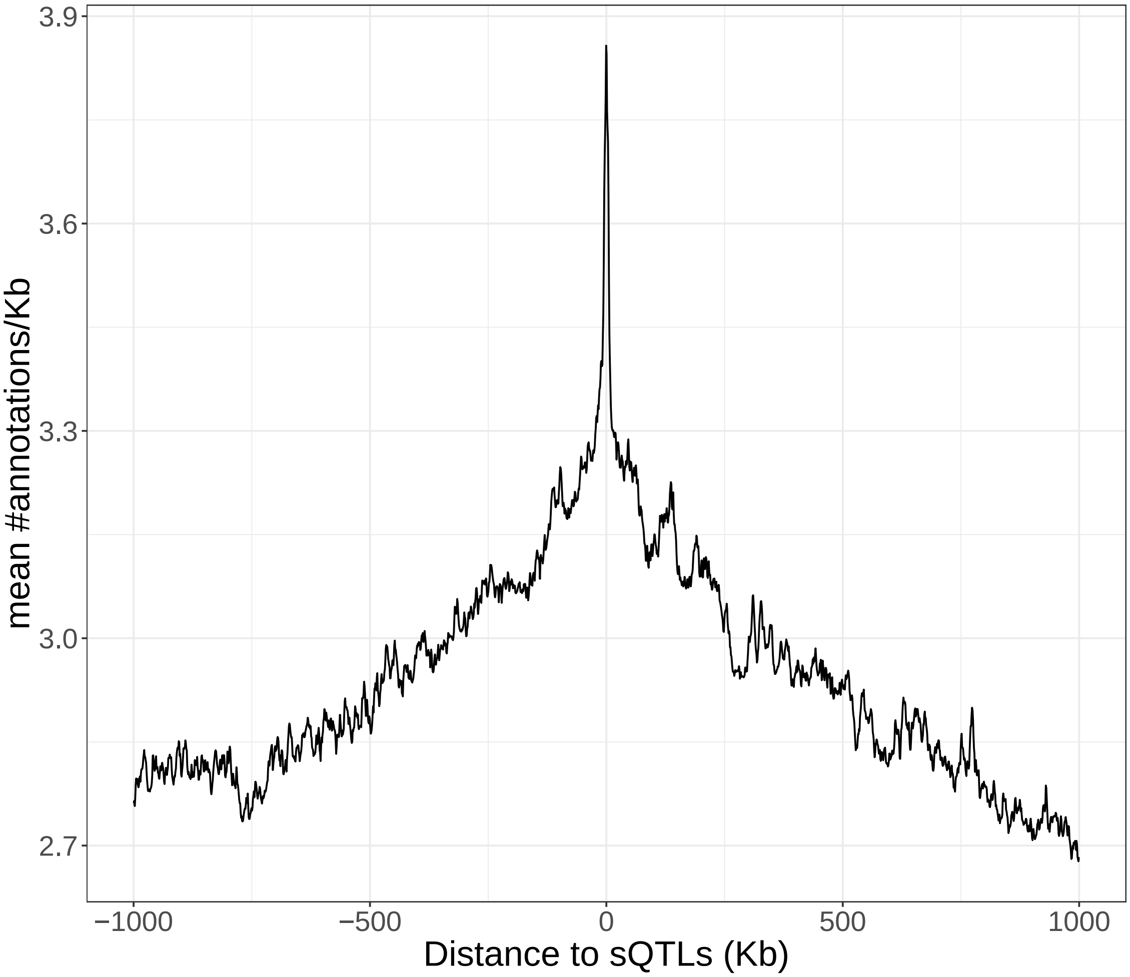

**Fig. S11.** Significant enrichments of sQTLs with respect to non-sQTLs (Fisher's exact test FDR $< 0.05$) in several functional annotations: **a)** Variant Effect Predictor categories and impact, GENCODE v19 protein coding and lincRNA exons and GWAS catalog; **b)** ENCODE RNA-binding protein eCLIP peaks and **c)** Ensembl Regulation data (see Methods). For each functional element, estimates and 95% confidence intervals for the enrichment $log_2$ odds-ratio (OR) are shown. FDR values are color-coded.

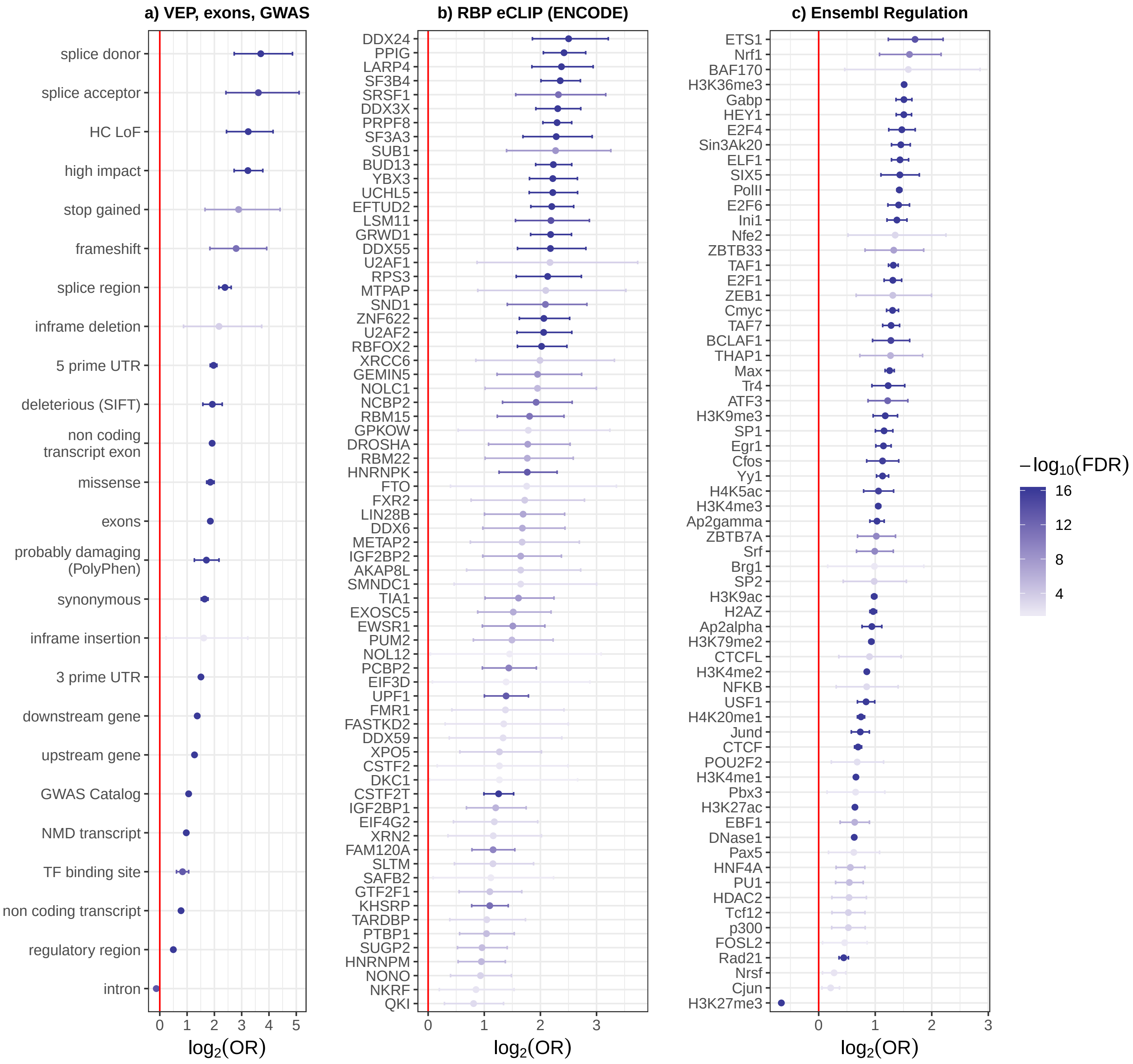

**Fig. S12.** Comparison of the enrichment of high (MD $\geq$ 0.2) and low (MD $<$ 0.1) effect size sQTLs (with respect to non-sQTLs, Fisher's exact test FDR $<$ 0.05) in a set of functional categories. For each effect size group and functional category, estimates and 95% confidence intervals for the enrichment $log_2$ odds-ratio (OR) are shown. TFBS and RBP correspond to pooled transcription factor and RNA-binding protein (RBP) binding sites, respectively.

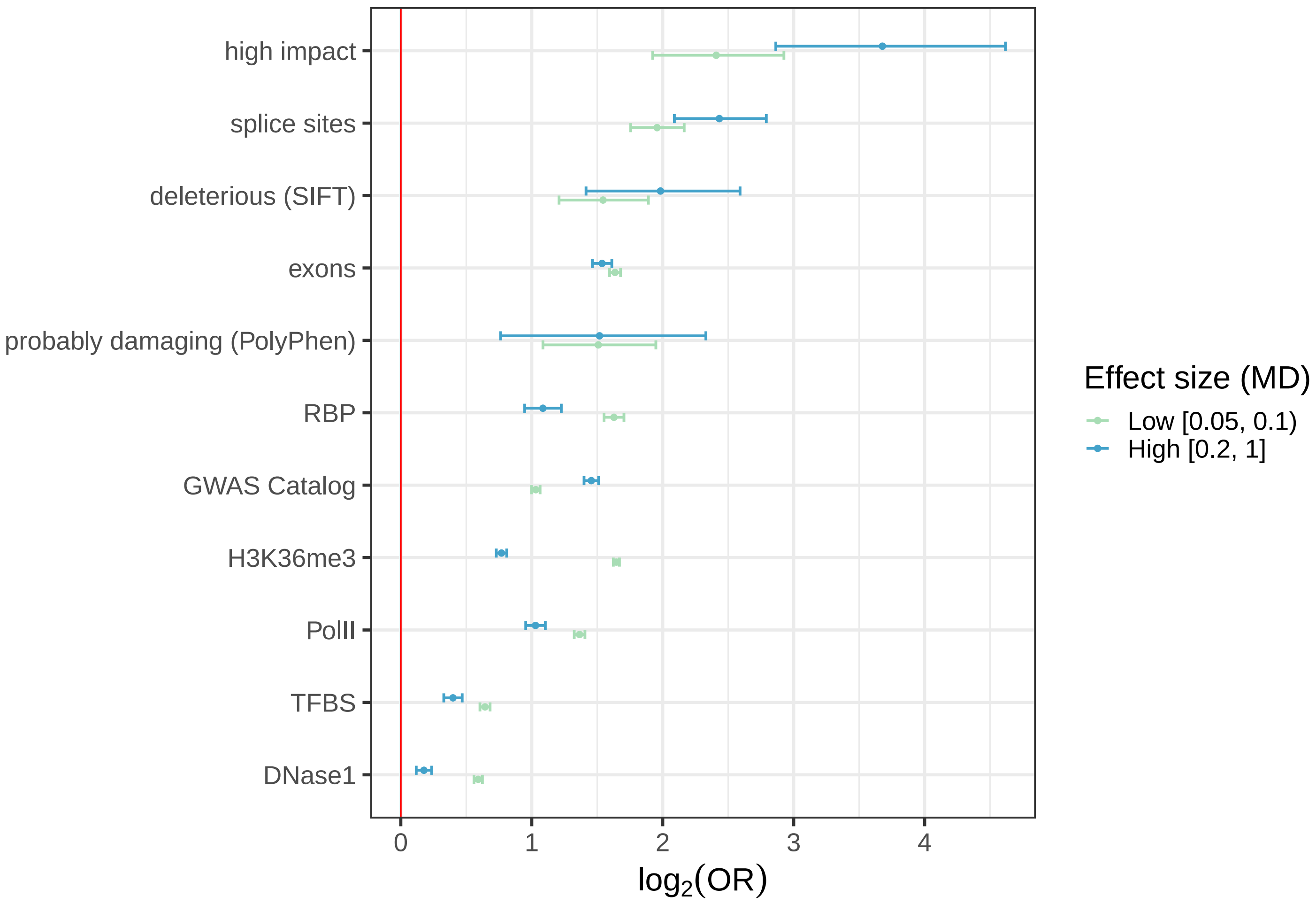

**Fig. S13**. Cumulative distribution of the distance to the closest splice donor or acceptor site from protein coding and lincRNA genes annotated in GENCODE v19, both for sQTLs and non-sQTLs.

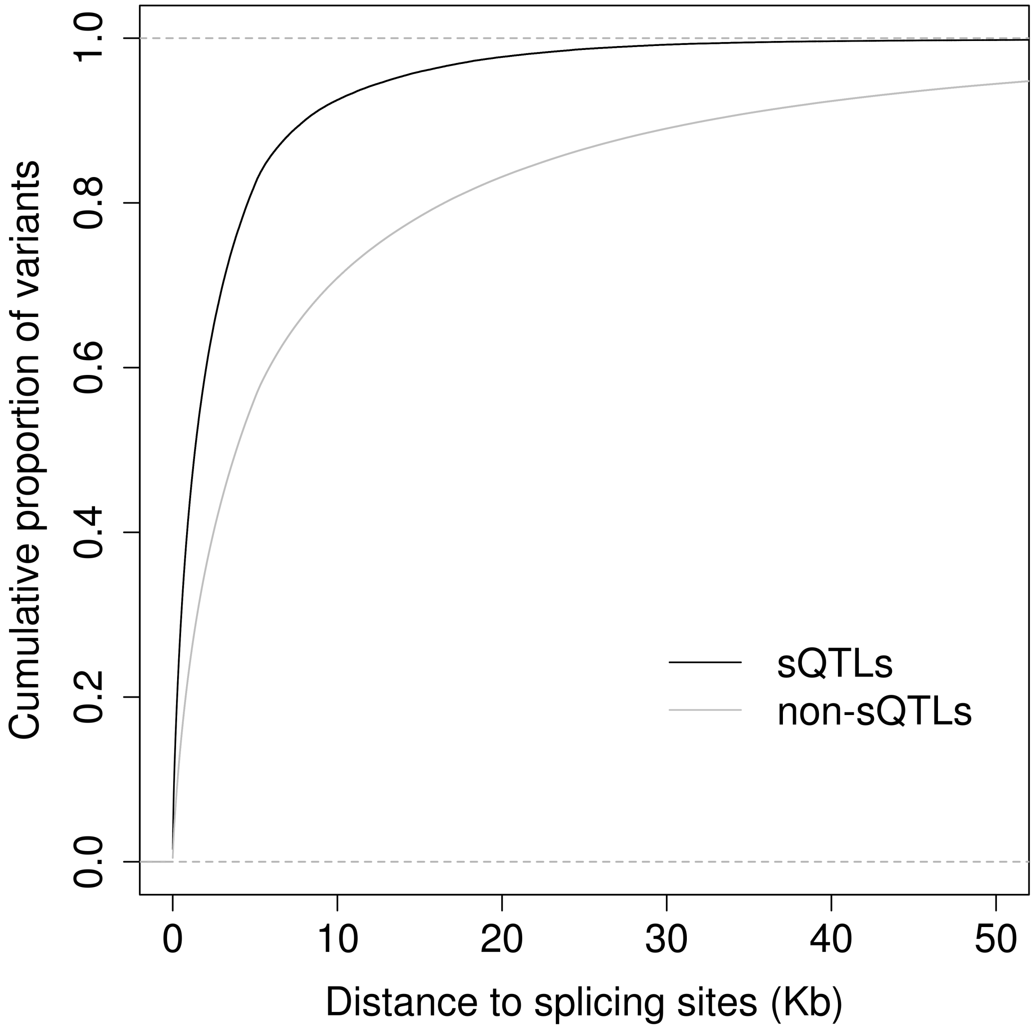

**Fig. S14**. Receiver Operating Characteristic (ROC) curves for 12 RNA-binding proteins (RBPs) corresponding to the classification of their eCLIP peaks by a gapped k-mer support vector machine (gkm-SVM). For each RBP, the solid line and the coloured area correspond, respectively, to the mean and standard error across the cross-validation folds. The mean area under the curve (AUC) for each RBP is shown between parentheses.

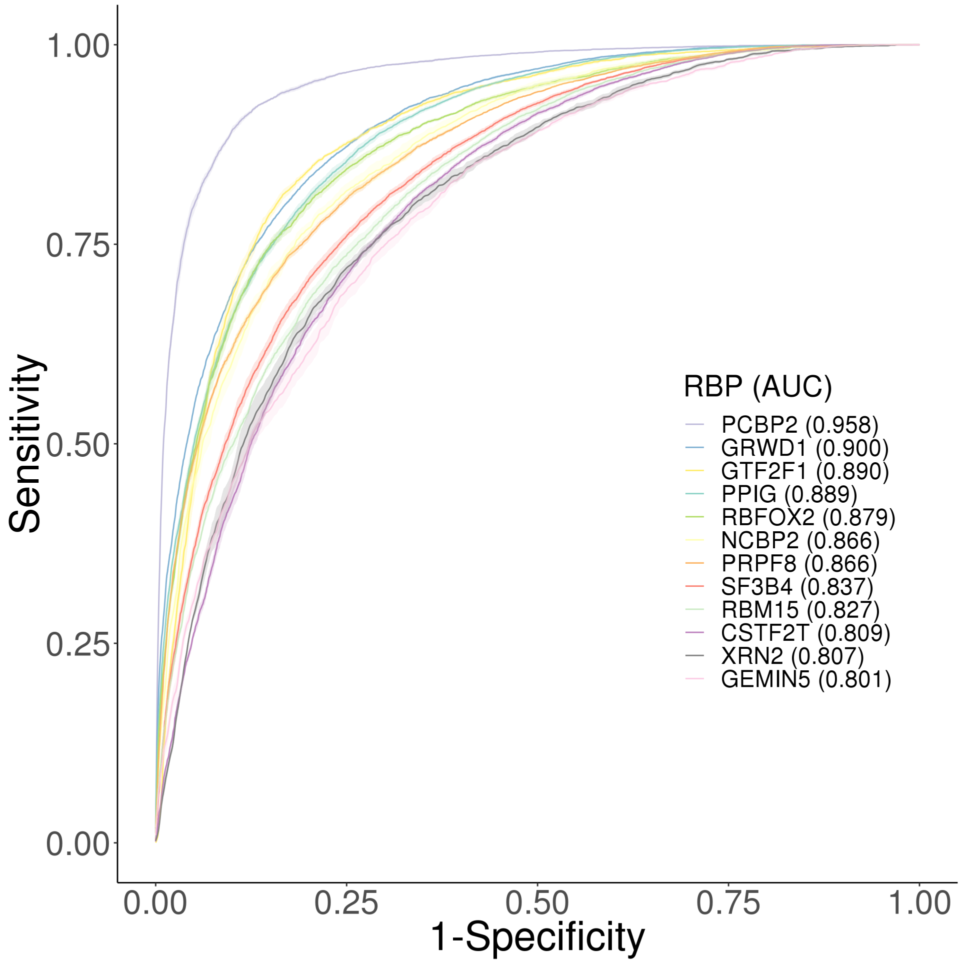

**Fig. S15.** Sequence logos for the predicted binding motifs of 12 RNA-binding proteins (RBPs), derived from the alignment of the 100 highest-scoring gkm-SVM 10-mers for each RBP. The proportion of each nucleotide (y-axis) is shown for each position of the sequence (x-axis).

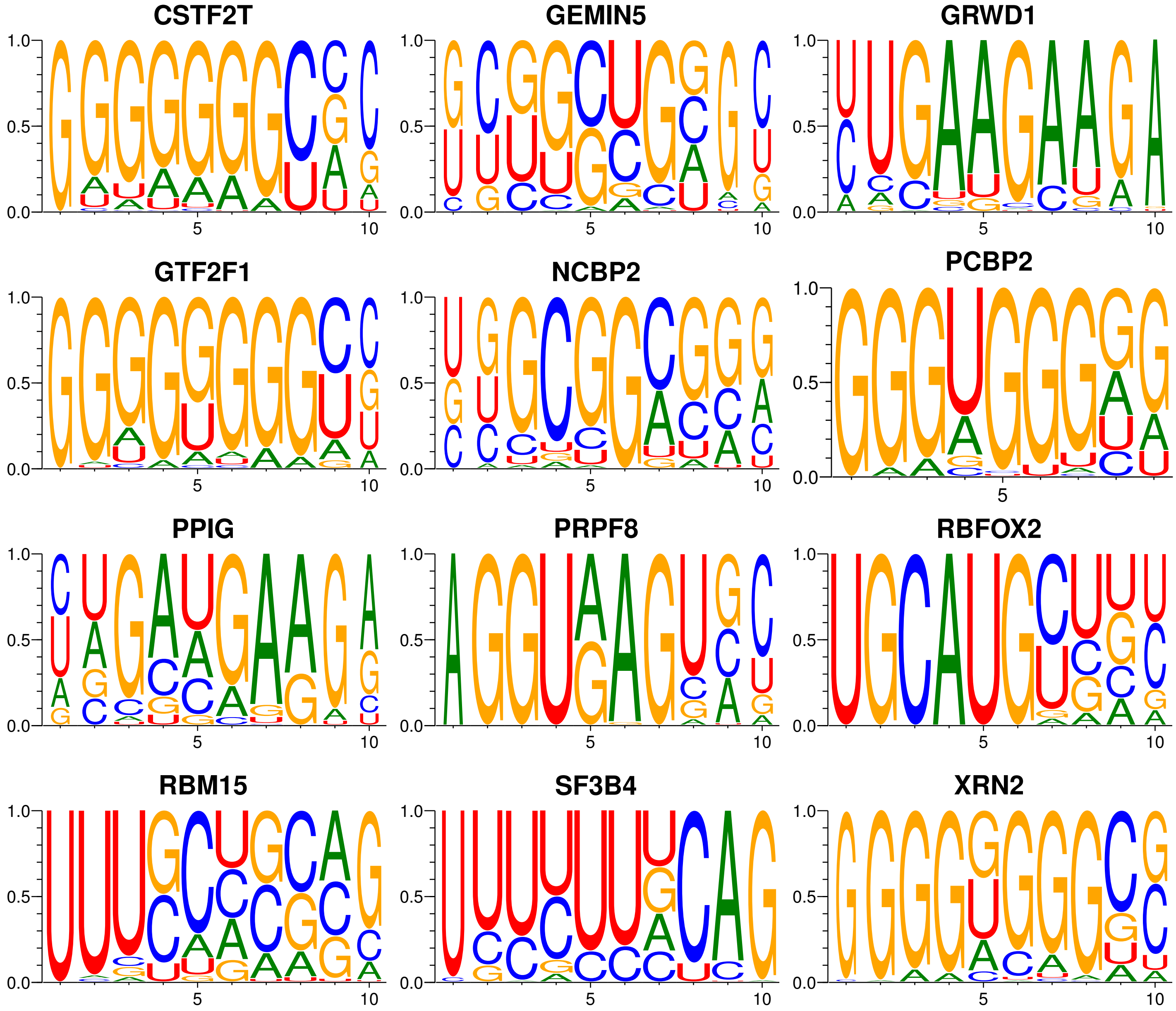

**Fig. S16.** **a)** Relative abundances of the most expressed isoforms in heart (left ventricle) from the gene PSMG4 (chr6:3,231,637-3,303,607, forward strand), for each genotype group at the rs4959783 locus (chr6:3,260,093, G/A in the forward strand). The least abundant isoforms are grouped in *Others*. The number of individuals in each genotype group is shown between parentheses. In reference homozygous individuals at rs4959783 (GG), isoform ENST00000438998 (blue) captures most of the expression of the gene and isoform ENST00000419065 (green) is not expressed. In contrast, in alternative homozygous individuals (AA) both isoforms have comparable splicing ratios. Heterozygous individuals (GA) display an intermediate behaviour. rs4959783 is an sQTL for PSMG4 in a total of 8 tissues ($\overline{MD} = 0.30$). **b)** Analogous representation for the gene TAMM41 (chr3:11,831,916-11,888,393, reverse strand) and the SNP rs9876026 (chr3:11,849,807, T/G in the reverse strand) in cerebellum. In this case, the abundance of isoform ENST00000455809 (blue) increases with the number of copies of the alternative allele (G) at rs9876026. In contrast, isoforms ENST00000457498 (pink) and ENST00000486090 (red) display the opposite behaviour, being less abundant in alternative homozygous individuals. rs9876026 is an sQTL for TAMM41 in a total of 46 tissues ($\overline{MD} = 0.18$).

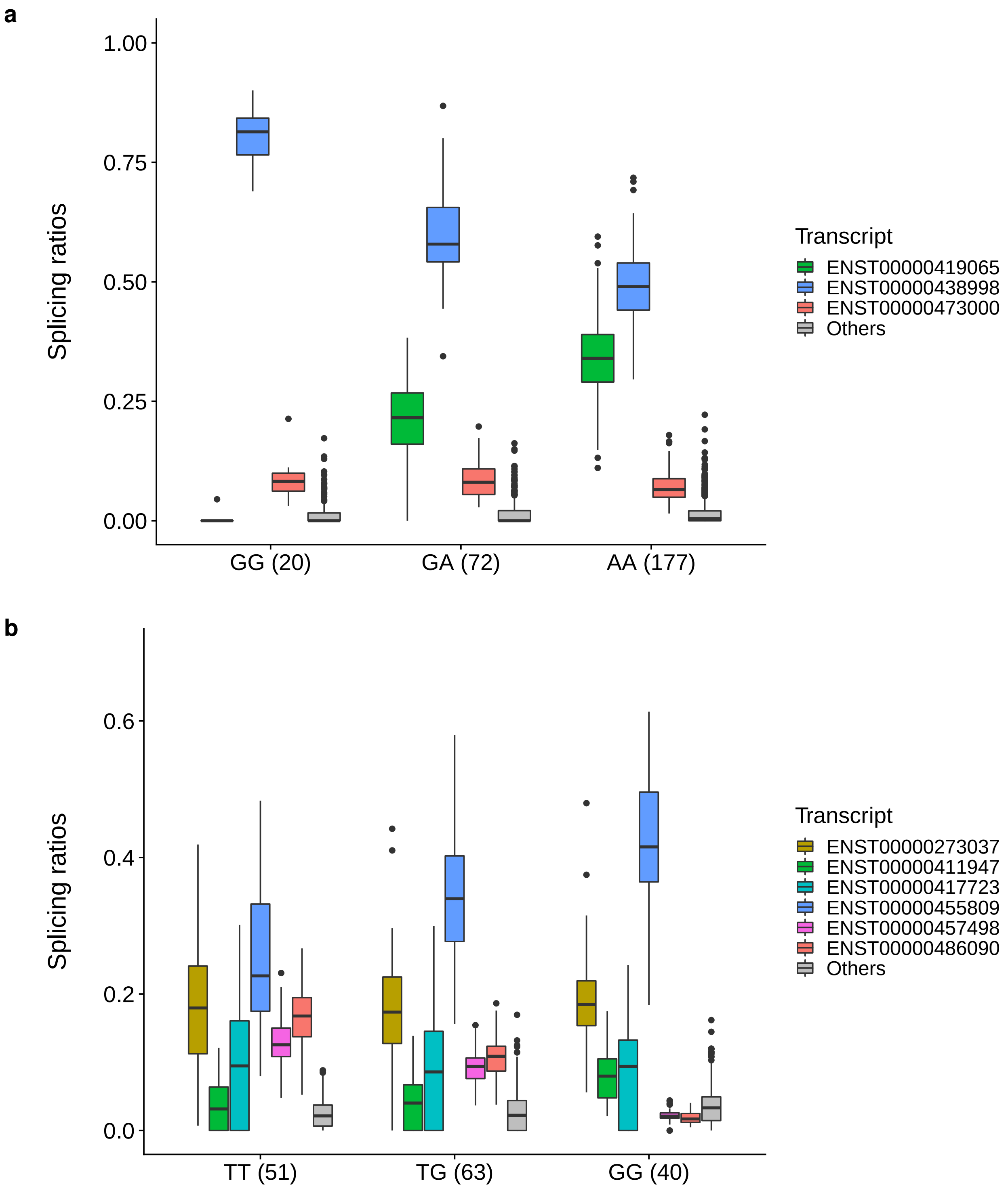

**Fig. S17.** Enrichment of sQTLs *vs* matched non-sQTLs in a set of allele-specific RBP binding (ASB) variants identified by BEAPR (Binding Estimation of Allele-specific Protein-RNA interaction) on the ENCODE eCLIP dataset (Fisher's exact test, FDR $< 0.05$). Estimates and 95% confidence intervals for the enrichment $log_2$ odds-ratio (OR) are displayed. FDR values are color-coded.

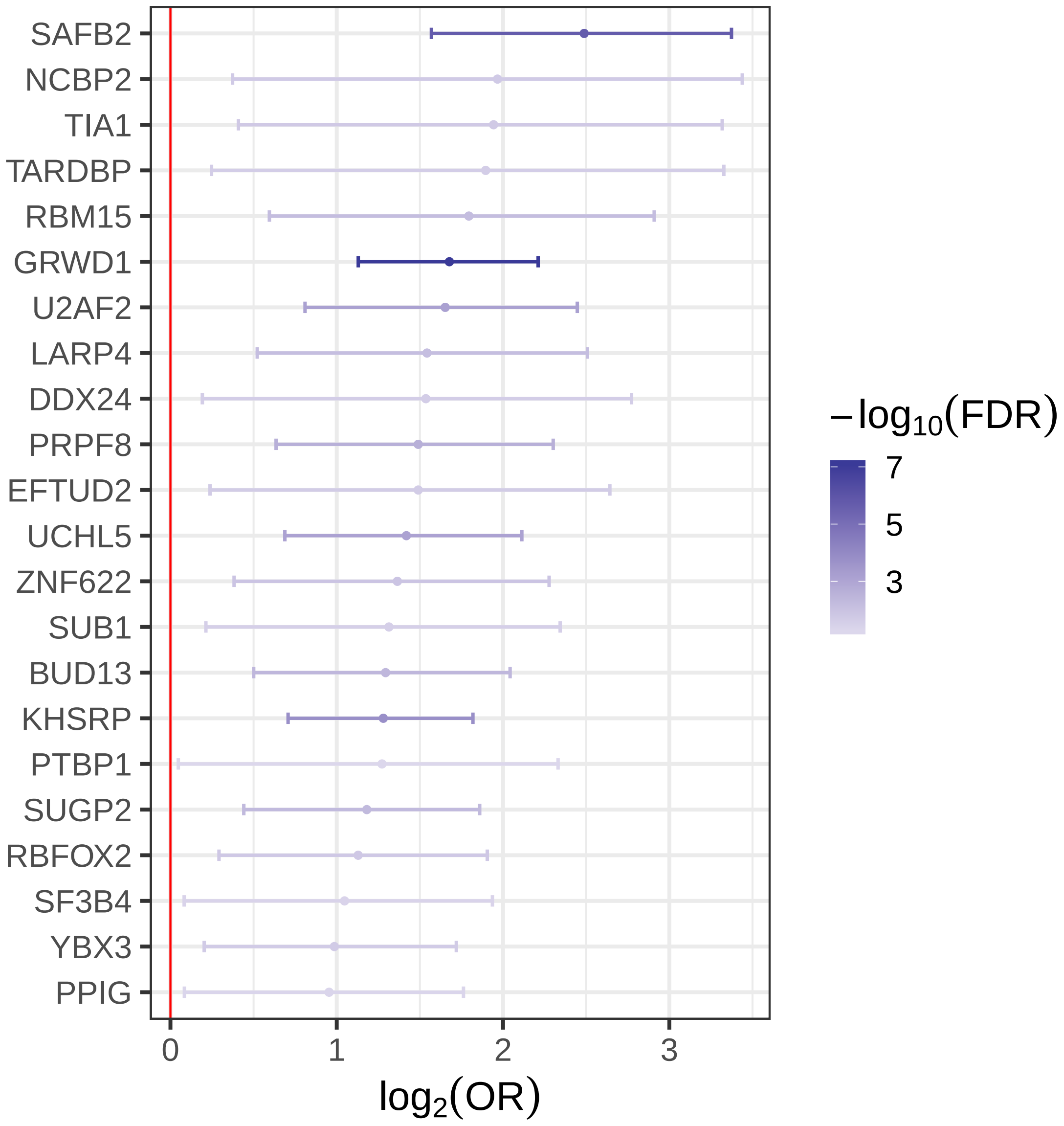

**Fig. S18.** Comparison of the distribution of effect sizes (MD values) for sQTLs in splice sites and RNA-binding protein (RBP) binding sites (pooled RBP eCLIP peaks from ENCODE). Variants falling in both categories have been removed.

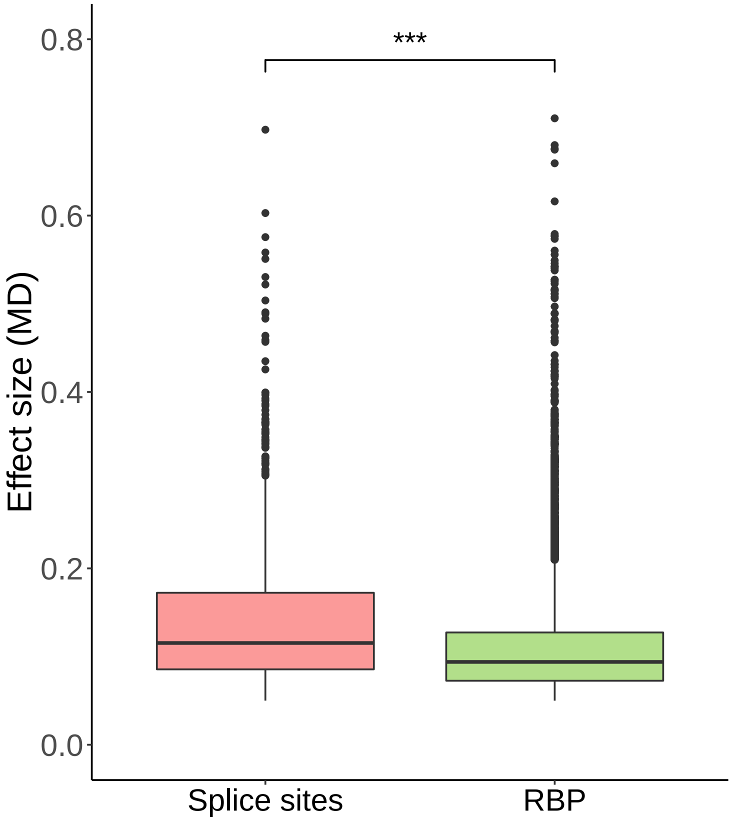

**Fig. S19. a)** Variant density (number of variants per Kb) along co- (blue) and post-transcriptionally spliced (orange) introns. **b)** Enrichment of sQTLs falling in post-transcriptionally spliced introns *vs* sQTLs falling in co-transcriptionally spliced introns (Fisher's exact test, FDR $< 0.05$, odds-ratio $\geq 1.5$). Estimates and 95% confidence intervals for the enrichment $log_2$ odds-ratio (OR) are displayed. FDR values are color-coded. TFBS and RBP correspond to pooled transcription factors and RNA-binding protein binding sites, respectively.

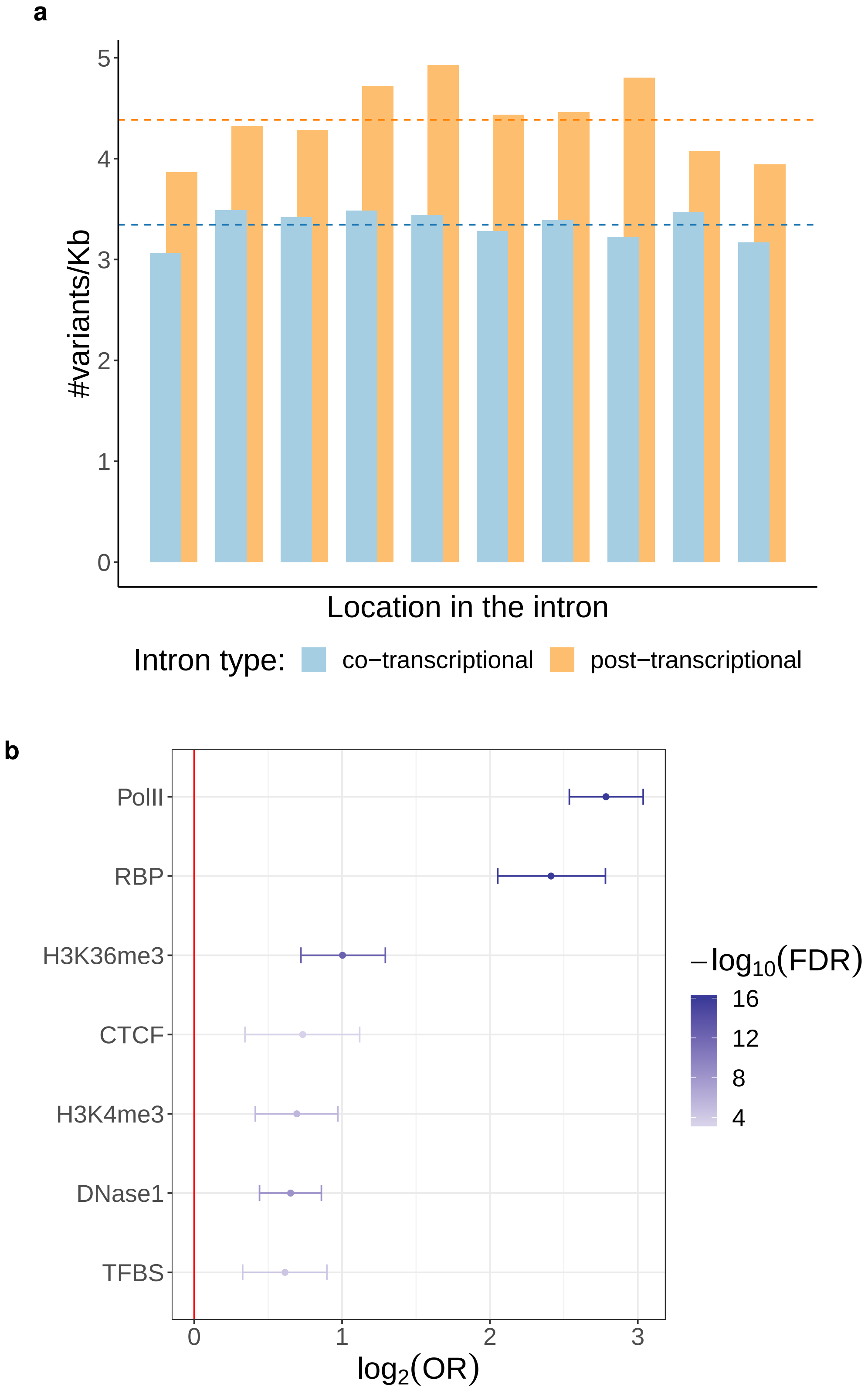

**Fig. S20**. Quantile-quantile (QQ) plots of *p* values for association with several traits and diseases, including asthma, breast cancer, coronary artery disease (CAD), heart rate, height, low-density lipoprotein (LDL) levels, rheumatoid arthritis and schizophrenia, both for sQTLs (black dots), and non-sQTLs (black solid line and grey area, representing the median and the middle 95% values across 10,000 random subsamplings of the non-sQTL set with the same size than the sQTL set). The identity line is shown in red.

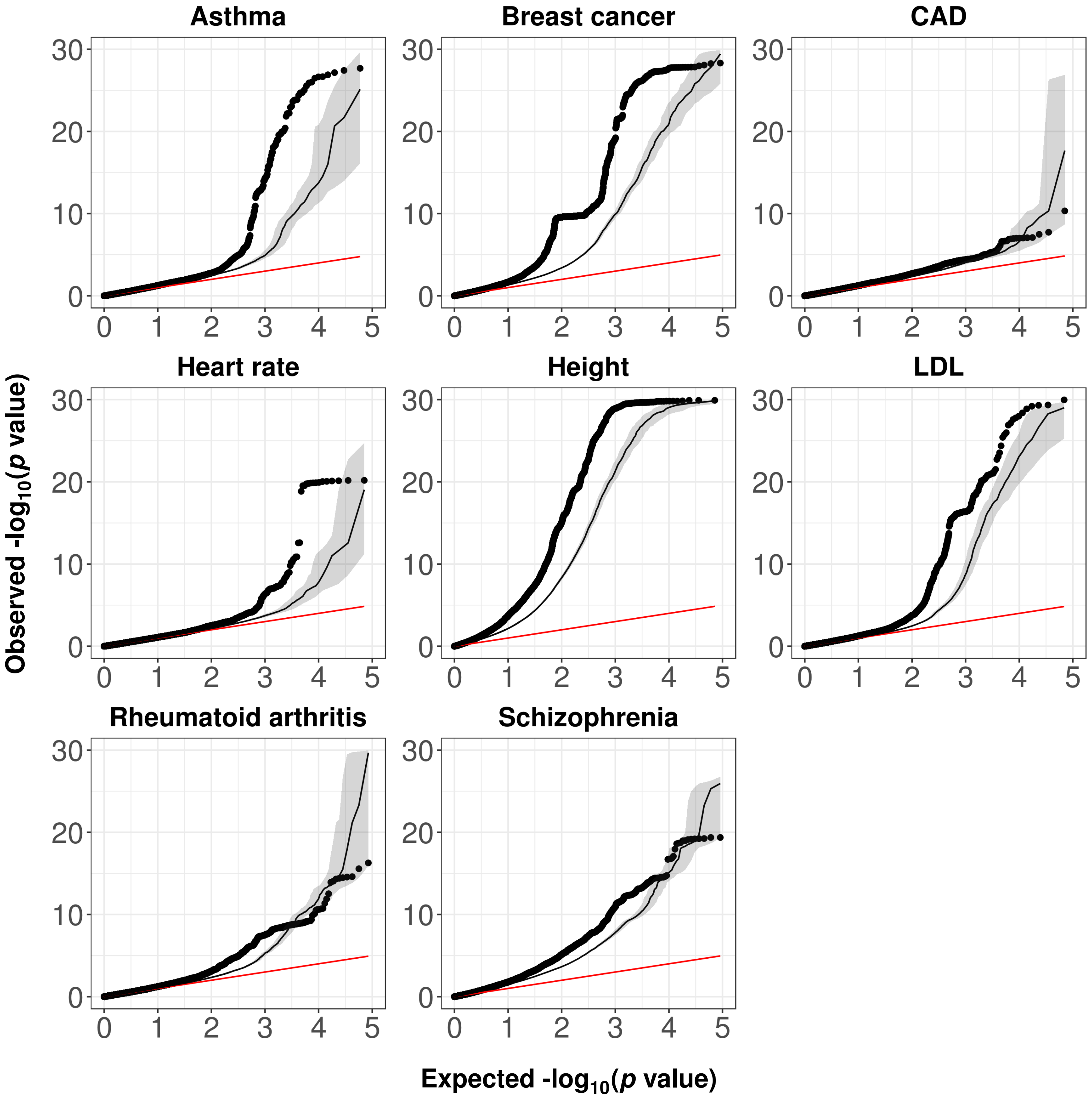

**Fig. S21**. Quantile-quantile (QQ) plots of *p* values for association with several traits and diseases, corresponding to sQTLs within the eCLIP peaks of RBFOX2, PRPF8, GTF2F1, PCBP2, SF3B4, GRWD1, PPIG, GEMIN5, CSTF2T, RBM15 and NCBP2 (coloured dots), and the remaining sQTLs (black dots). The identity line is shown in red.

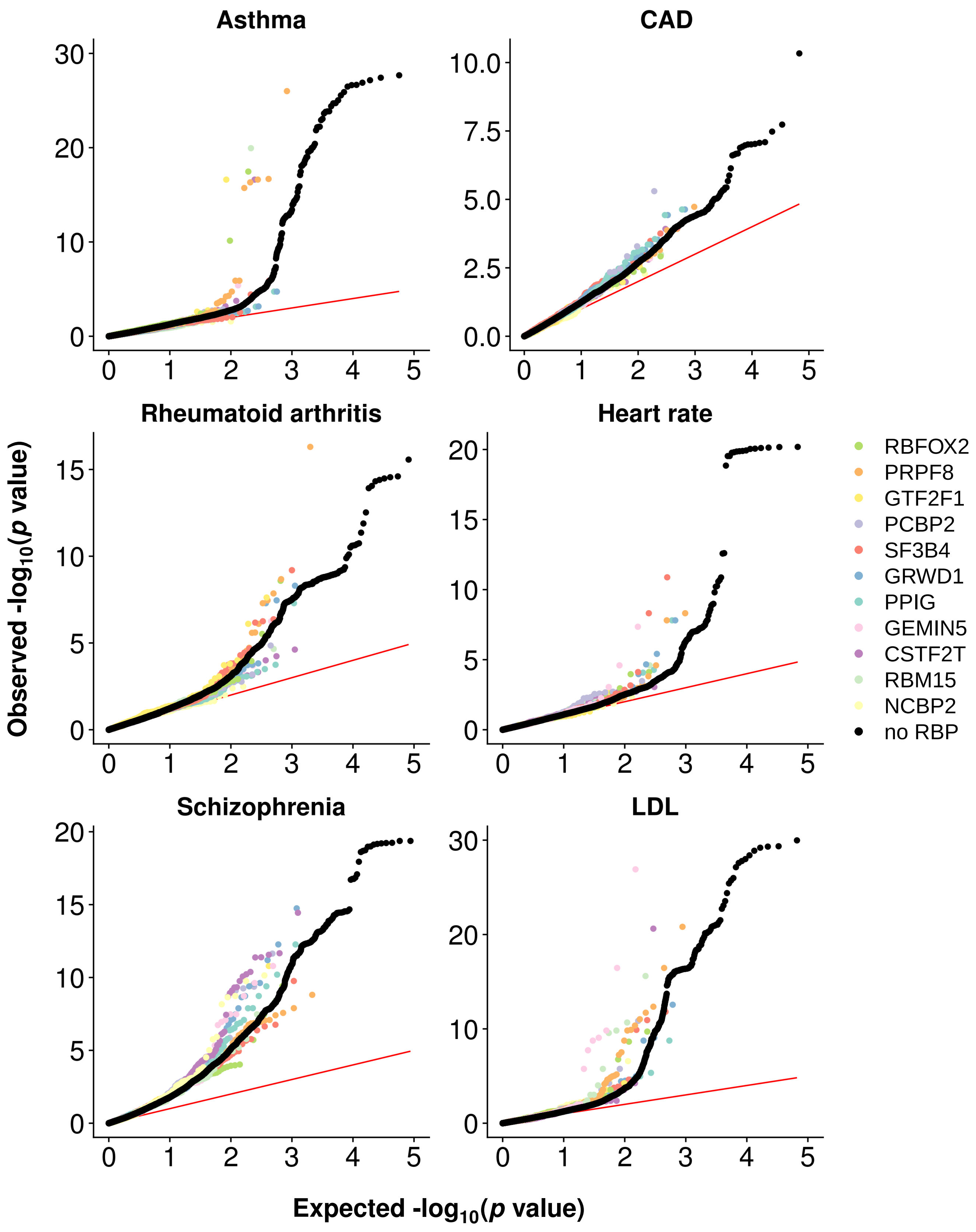

**Fig. S22.** **a)** Relative abundances of the most expressed isoforms in lung from the gene GSDMB (chr17:38,060,848-38,076,107, reverse strand), for each genotype group at the rs11078928 locus (chr17:38,064,469, T/C). Solid and shaded colors correspond to protein coding and non-coding isoforms, respectively. The least abundant isoforms are grouped in *Others*. The number of individuals in each genotype group is shown between parentheses. From all protein coding isoforms, GSDMB-003 (green) is the most abundant in reference homozygous individuals at rs11078928 (TT). In contrast, in alternative homozygous individuals (CC), isoform GSDMB-002 (red) is the most expressed. Isoform GSDMB-001 (blue) shows a pattern analogous to GSDMB-002 (red). Heterozygous individuals (TC) display an intermediate behaviour. The abundances of non-coding isoforms barely change with the genotype at rs11078928. **b)** Exonic structure of the isoforms GSDMB-001, GSDMB-002 and GSDMB-003 and location of the rs11078928 SNP in exon 6 (marked with an arrow).

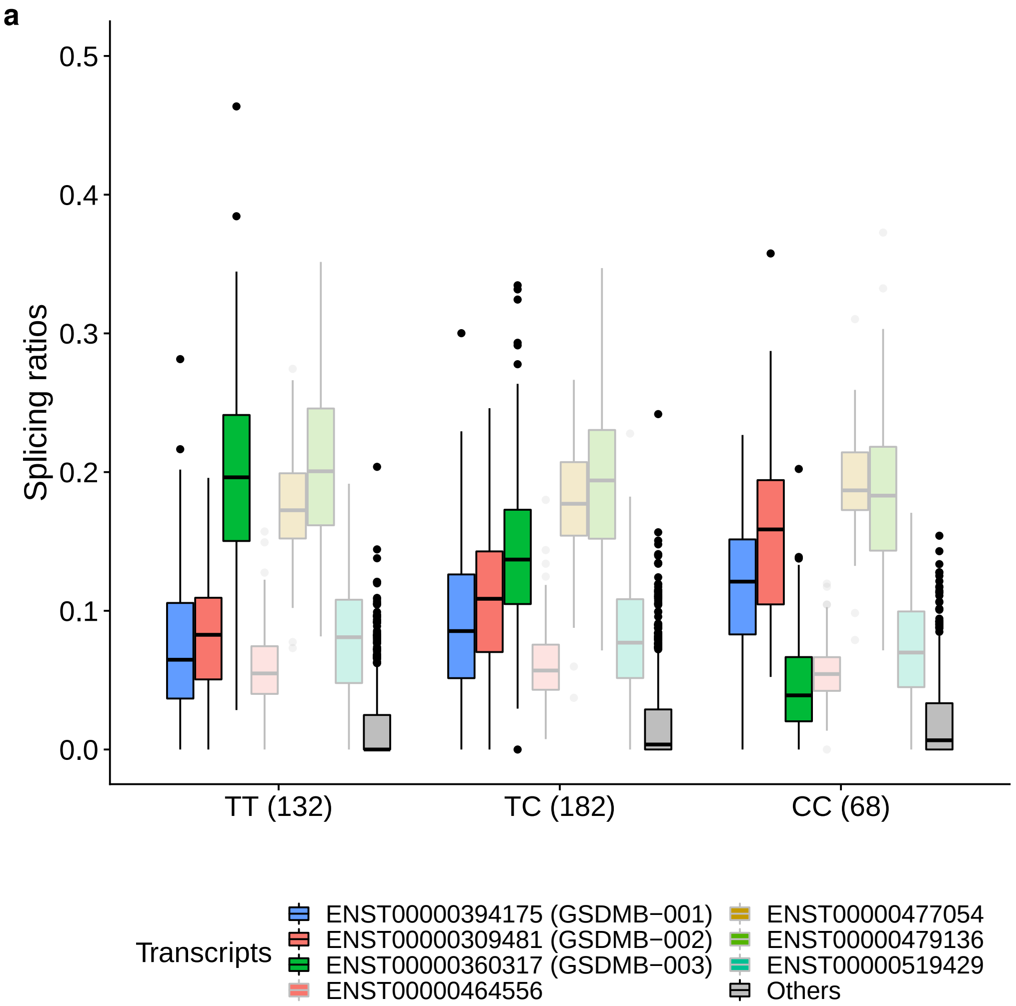


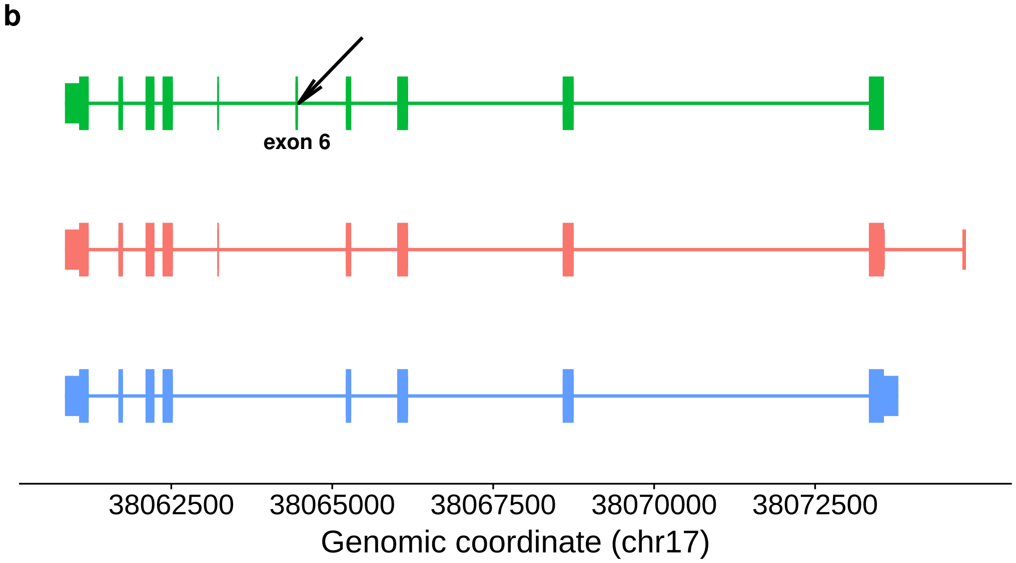

**Fig. S23**. Comparison of the classification performance for all RNA-binding proteins, RBPs (measured as the mean cross-validation gkm-SVM ROC AUC per RBP), between different choices of the `l` (word length) and `k` (number of informative columns) parameters. Default values (`l` = 10, `k` = 6) are marked in bold.

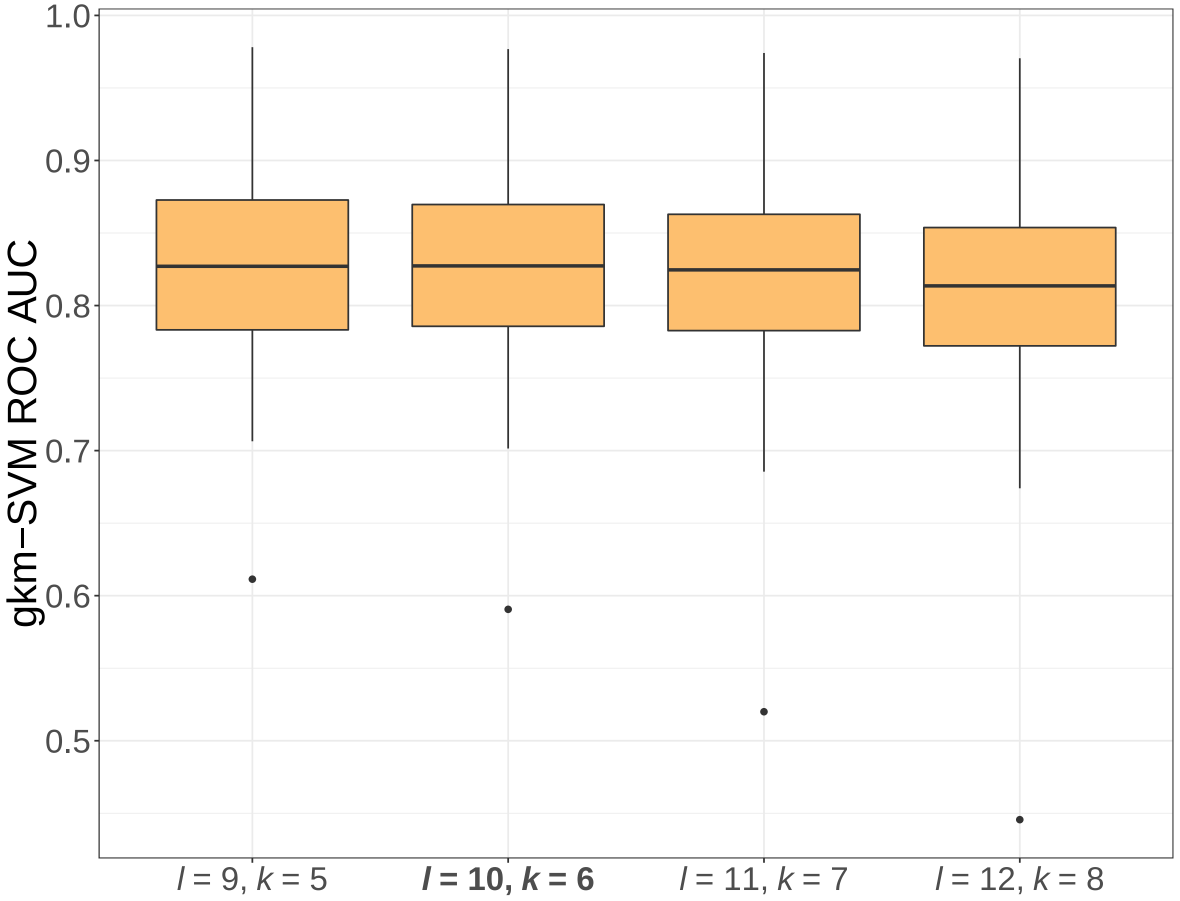

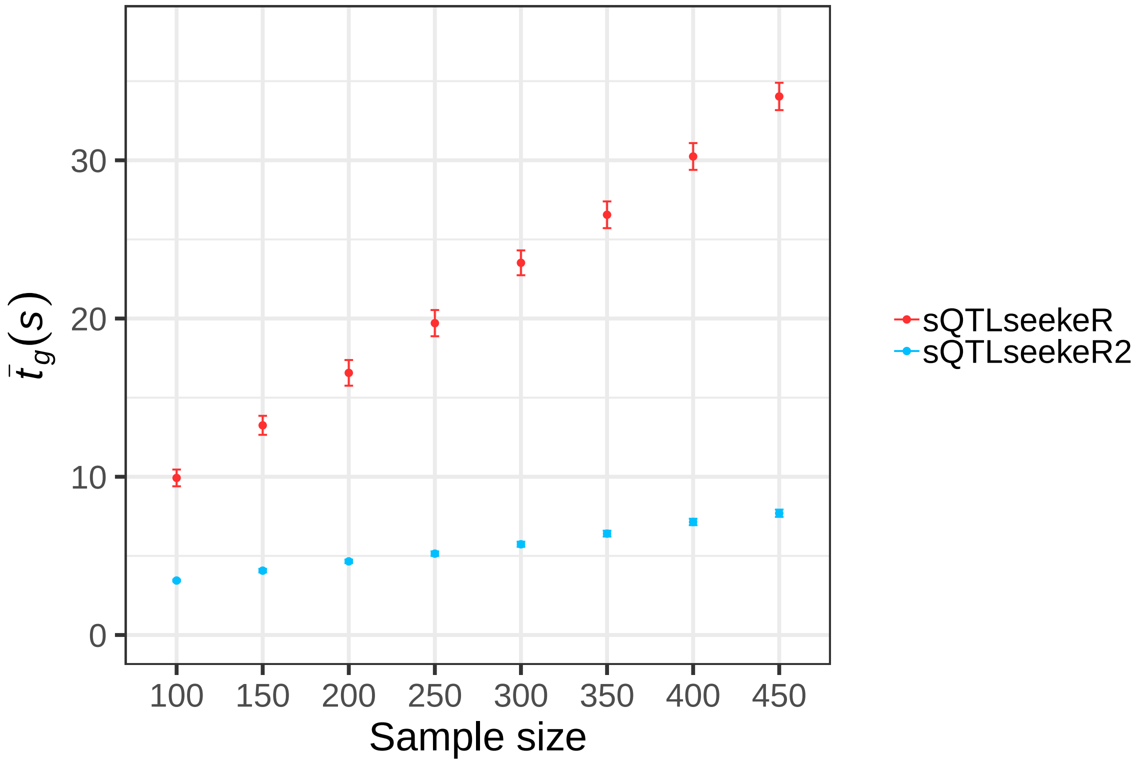


**Fig. S24**. Mean running time per gene ($\bar{t}_g$) of `sQTLseekeR` (red) and `sQTLseekeR2` (blue). Error bars represent standard errors. A nominal pass of both versions of the software was run on 10 sets of 100 randomly selected genes (on average 135 variants tested per gene), across a wide range of sample sizes, obtained by downsampling the GTEx Muscle Skeletal transcript expression dataset (tissue with the largest sample size available). To make results comparable, we set common options to the same values, did not include covariates and did not perform additional filtering steps only available in `sQTLseekeR2`. We asked `sQTLseekeR` to perform $10^7$ Monte Carlo generations.

**Fig. S25.** **a)** Type I error of `sQTLseekeR2` in simulated datasets with different sample size ($n$) and number isoforms studied ($q$). The horizontal green line marks the significance level selected ($\alpha$ = 0.05). **b)** Power of `sQTLseekeR2` across simulated datasets with different values of $n$, $q$ and sQTL effect sizes (MD values). See Supplementary Note 1 for details on the simulation.

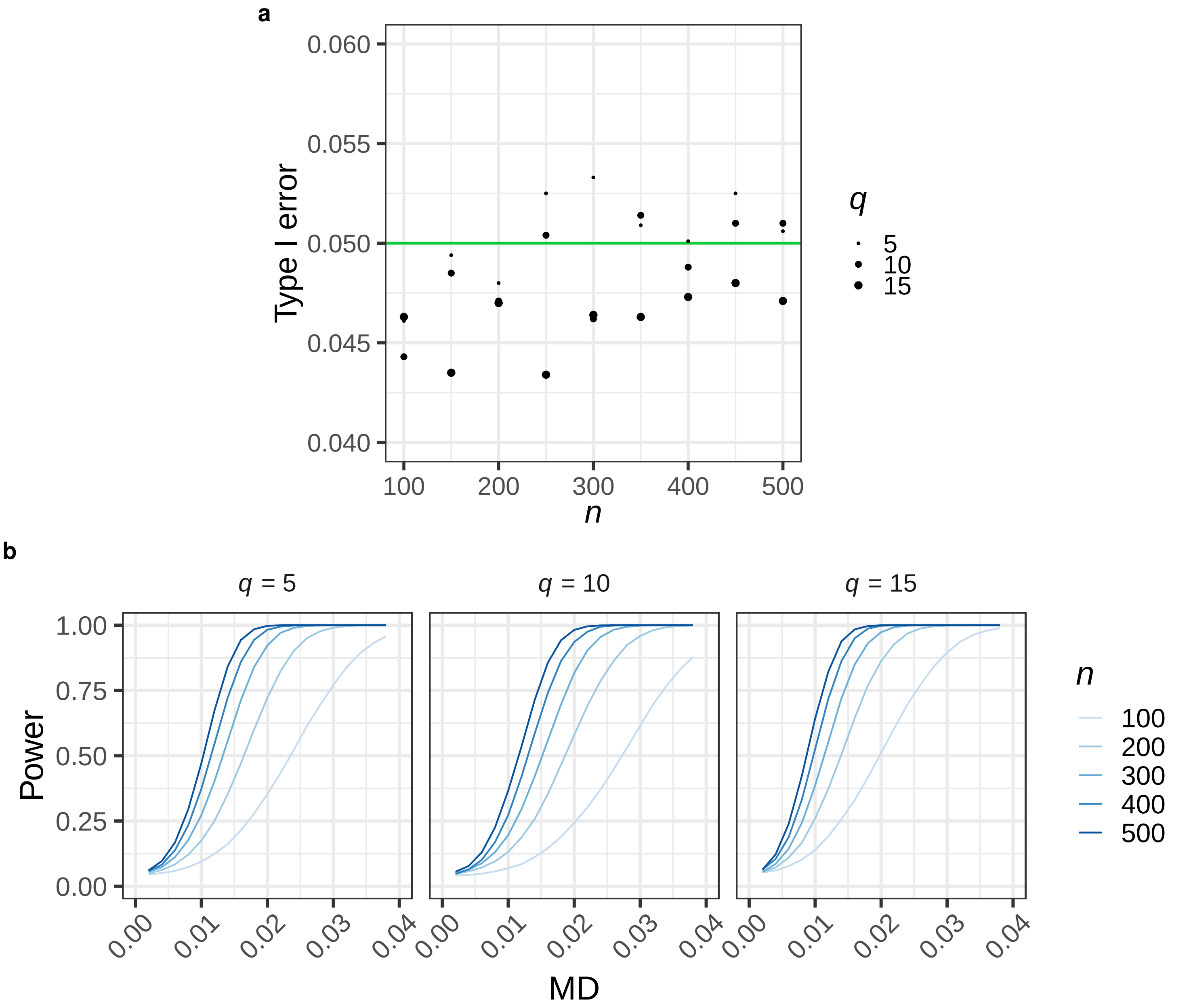

**Fig. S26**. **a)** Overlap between the sQTLs identified by `sQTLseekeR2` using as input `RSEM` transcript quantifications and `LeafCutter` quantifications, measured at the level of sGenes (*Jaccard* index) and pairs sQTL-sGene ($\pi_1$) across tissues. **b)** Proportion of sGenes (y-axis) identified only with transcript quantifications (TQ), only with `LeafCutter` quantifications (LC), or with both (Common) *vs* tissue sample size (x-axis). **c)** Mean number of reads supporting an intron cluster in each tissue for LC-exclusive (y-axis) *vs* Common (x-axis) sQTLs. Error bars represent standard errors. The identity line is shown in black. For visualization purposes, whole blood ($x = 956$, $y = 546$) is not displayed given its outlier behaviour. **d)** Representation in two dimensions, for each tissue, of the vector of proportions of the different types of AS events associated with Common and TC-exclusive sQTLs, obtained by PCA. Tissue color codes are shown in Table S2. **e)** Comparison of the location of TQ-exclusive and LC-exclusive sQTLs.

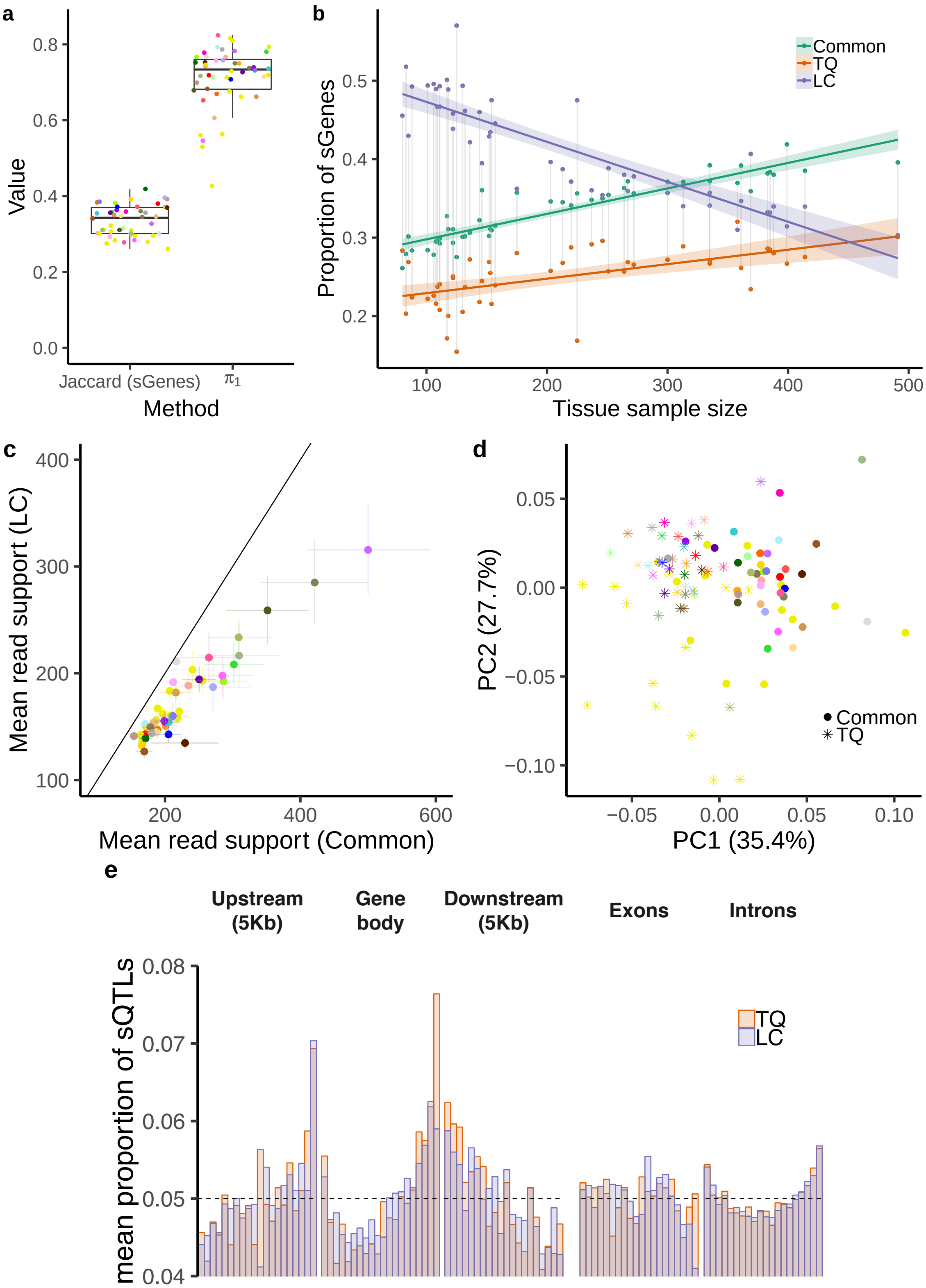

**Fig. S27.** **a)** Proportion of sGenes (over tested genes) per tissue (y-axis) with respect to the tissue sample size (x-axis) identified in GTEx V8. Tissue color codes are shown in Table S9. **b)** Overlap between the sGenes identified by `sQTLseekeR2` in GTEx V7 and V8. Distribution of the *Jaccard* index (left) and the minimum overlap (that is, the number of common sGenes between V7 and V8 over the total number of sGenes identified in the smallest set, i.e. V7, right), computed per tissue. To obtain these metrics only the variant-gene-tissue trios tested in both V7 and V8 are considered. **c)** Relative change in median sQTL effect size (median MD value) between V8 and V7, per tissue (y-axis), *vs* the relative change in tissue sample size (x-axis). **d)** Overlap between the sQTLs identified by `sQTLseekeR2` and the ones obtained using `LeafCutter` + `FastQTL` by the GTEx Consortium, measured at the level of sGenes (*Jaccard* index) and pairs sQTL-sGene ($\pi_1$) across tissues.

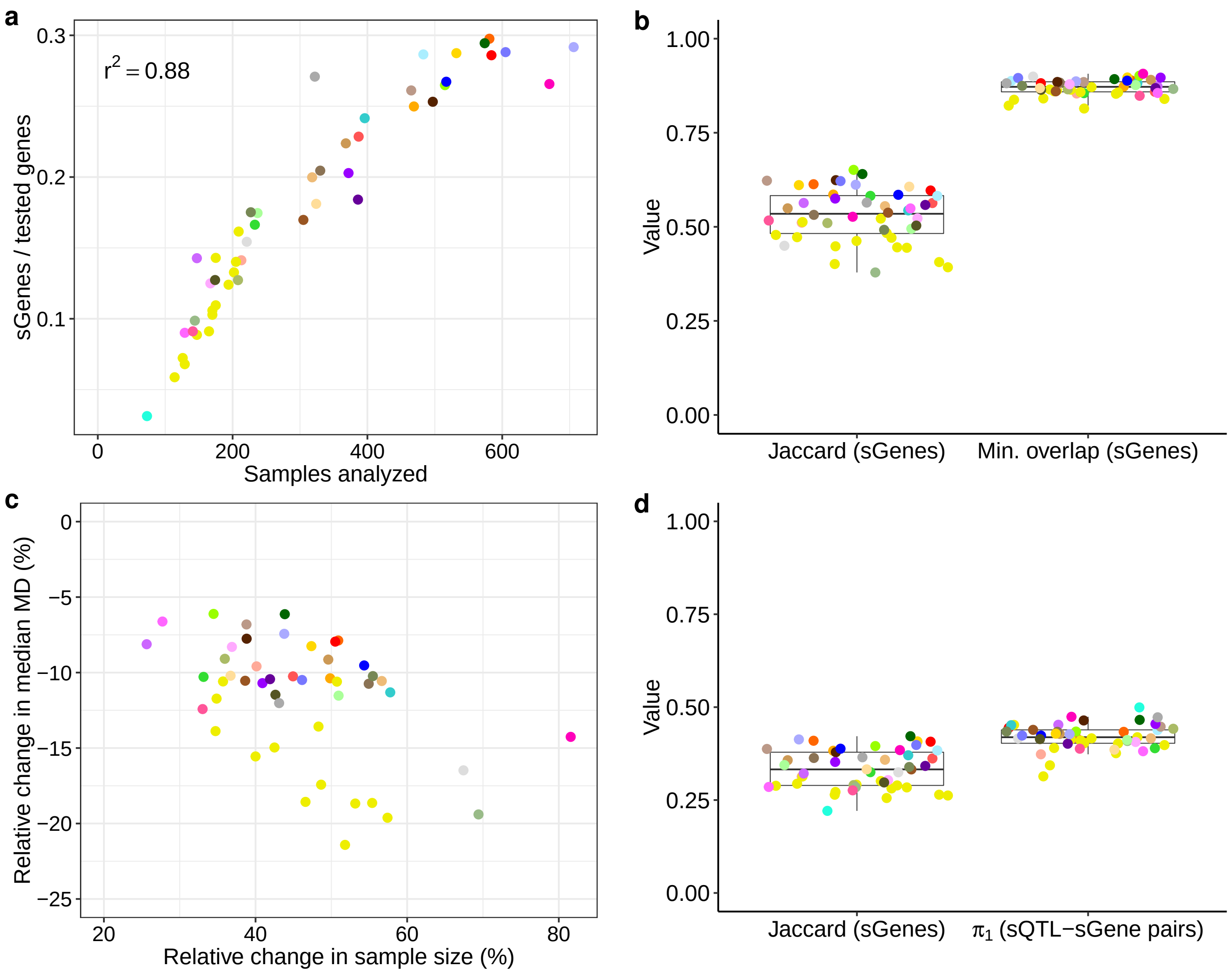
